# Benchmarking, detection, and genotyping of structural variants in a population of whole-genome assemblies using the SVGAP pipeline

**DOI:** 10.1101/2025.02.07.637096

**Authors:** Ming Hu, Penglong Wan, Chengjie Chen, Shuyuan Tang, Jiahao Chen, Liang Wang, Mahul Chakraborty, Yongfeng Zhou, Jinfeng Chen, Brandon S. Gaut, J.J. Emerson, Yi Liao

**Affiliations:** Key Laboratory of Biology and Genetic Improvement of Horticultural Crops (South China), Ministry of Agriculture and Rural Affairs, College of Horticulture, South China Agricultural University, Guangdong 510642, China; Tropical Crops Genetic Resources Institute, Chinese Academy of Tropical Agricultural Sciences & National Key Laboratory for Tropical Crop Breeding & Laboratory of Crop Gene Resources and Germplasm Enhancement in South China, Ministry of Agriculture and Rural Affairs & Key Laboratory of Tropical Crops Germplasm Resources Genetic Improvement and Innovation of Hainan Province, Hainan, 571101, China; Department of Biology, Texas A&M University, College Station, TX, 77843, USA; State Key Laboratory of Integrated Management of Pest Insects and Rodents, Institute of Zoology, Chinese Academy of Sciences, Beijing, 100101, China; Department of Ecology and Evolutionary Biology, University of California, Irvine, CA, 92697, USA

## Abstract

Comparisons of complete genome assemblies offer a direct procedure for characterizing all genetic differences among them. However, existing tools are often limited to specific aligners or optimized for specific organisms, narrowing their applicability, particularly for large and repetitive plant genomes. Here, we introduce SVGAP, a pipeline for structural variant (SV) discovery, genotyping, and annotation from high-quality genome assemblies at the population level. Through extensive benchmarks using simulated SV datasets at individual, population, and phylogenetic contexts, we demonstrate that SVGAP performs favorably relative to existing tools in SV discovery. Additionally, SVGAP is one of the few tools to address the challenge of genotyping SVs within large assembled genome samples, and it generates fully genotyped VCF files. Applying SVGAP to 26 maize genomes revealed hidden genomic diversity in centromeres, driven by abundant insertions of centromere-specific LTR-retrotransposons. The output of SVGAP is well-suited for pan-genome construction and facilitates the interpretation of previously unexplored genomic regions.

## Background

Structural variants (SVs) are commonly defined as genome alterations between individuals in either the order or content of DNA spanning 50 or more base pairs (bp) (1–3). These include simple variants like deletions, insertions, duplications, translocations, or inversions, as well as more complex DNA rearrangements (4–6). Despite being recognized as major sources of functional variation for over a century (7, 8), SVs are still the least well-characterized genetic variants, especially in plants (9–11).

Only recently has a growing emphasis been placed on genome-wide analyses of SVs, driven by significant advancements in detection technologies (12–17). These studies highlight the important roles that SVs play in processes ranging from mutation to evolution (18–20), from ecology to speciation, and from human health and survival to crop genome diversity, domestication, migration history, breeding, and traits (15, 21–23). Furthermore, SVs impact more of the genome than single-nucleotide variations (SNVs) (24), and recent studies demonstrated that incorporating SVs into pangenome graphs significantly enhance the power of genome-wide association studies, capture missing heritability, and empower crop breeding (16). Comprehensive and accurate SV discovery and genotyping is required for the effective study of these important issues.

Detection and analysis of SVs has greatly benefited from the rapid advancement of sequencing technologies. While early studies (e.g., those employing cytogenetic approaches or microarray assays) allowed us to probe variation in genome structure, sequencing-based methods have the potential to identify both large and small SVs at a base pair resolution, can probe any region that can be sequenced, and, perhaps most importantly, permit studies to scale to large sample sizes quickly and inexpensively. Sequencing methods can be broadly categorized into three major groups: short-read reference mapping, long-read reference mapping, and whole genome assembly-based methods (2). To date, nearly 80 pieces of software based on short-read sequencing and 50 based on long-read sequencing have been developed (25, 26). They adopt diverse algorithms for resolving different types of sequencing platforms and experimental scenarios. Benchmark analyses reveal that no single tool achieves optimal performance for SV detection, with tradeoffs between accuracy and completeness being common(27, 28). Furthermore, benchmark experiments based on simulated data in animals revealed that short-read-based methods capture only a small proportion of SVs (i.e. ∼50%) (29), while long-read-based methods achieve a higher but still incomplete rate of between 70% and 84% (30). Benchmark analyses in plants are relatively rare compared to animals. However, due to their highly repetitive and complex nature, it seems unlikely that performance in plant genomes will exceed performance in other organisms. Indeed, SV detection continues to pose significant challenges. Researchers are actively developing new tools employing cutting-edge strategies and algorithms, such as pangenomics (31, 32) and machine learning (33, 34), aiming to achieve a comprehensive description of the entire landscape of SVs across all genomic contexts (35).

A *de novo* assembly-based strategy is widely anticipated to be a superior approach for characterizing SVs (36). Its ability to identify all differences between two haplotypes is theoretically limited only by the ability to accurately assemble the haplotypes (37). This capability has recently seen dramatic improvements in a very short time. Such dramatic advances are already providing access to previously unexplored genomic contexts, including challenging centromeric regions (38). Numerous studies have uncovered a significant proportion of hidden SVs that are missed by short-read and even long-read-based methods (39). However, the extensive effort and cost required to generate high-quality genome assemblies have often led to the neglect of this method. Fortunately, sequencing technologies and computational methods have become increasingly feasible and cost-effective, making reference-grade genome assemblies far more accessible (40). Consequently, population-scale high-quality genome assemblies are now widely attainable, and are becoming routine parts of the toolkit of biologists studying genetic variation (14). In this context, assembly-based variant calling and joint genotyping methods have emerged as attractive options for SV detection, offering indispensable benefits in genetic discovery and serving as valuable complements to pangenome construction (41, 42).

There are several methods available for calling SVs by aligning assembled genomes to a reference (26). These methods typically require a pre-alignment process before SV calling. Some methods, such as SyRI (43), MUM&Co (44), SVMU (39), and Assemblytics (45), rely on whole genome alignments from specific aligners. Others, such as Dipcall(46), SVIM-asm(47), PAV(37), and cuteSV (48), may also employ large contig alignments. Additionally, certain aligners themselves, like minimap2 (49), GSAlign (50), and AnchorWave (51), offer the option to call SVs during the alignment process. However, it is worth noting that most of these SV callers were developed and tested on a narrow range of taxa (typically animal genomes), which naturally raises questions about their effectiveness in plants. Plant genome structures are more diverse than mammalian genomes, often exhibiting variation in ploidy, greater sequence diversity, extensive rearrangements, and a high density of repetitive sequences (52).

When developing a *de novo* assembly-based structural variant (SV) discovery method, three key aspects should be taken into account. First, since this method relies on whole genome alignments as input, it is essential to validate the performance of alignment tools on the genomes being analyzed.

Second, a comprehensive set of SV “truth sets” should be employed for benchmarking purposes. Finally, the tools developed should be suitable for application across populations and leverage population-level data to enhance genotyping; to our knowledge, no such tools currently exist.

In this work, we introduce **SVGAP (S**tructural **V**ariants **G**enotyping of **A**ssemblies on **P**opulation scales), a flexible pipeline to detect, genotype, and annotate SVs in large samples of *de novo* genome assemblies. It compares each sample to a reference genome in whole-genome alignments to call SVs. The SVs identified are subsequently combined across samples to generate a nonredundant call set. Each SV call in this call set can be further re-genotyped by examining local alignment information specific to each sample to produce fully genotyped VCF (Variant Call Format) files. Our pipeline can optionally be applied to detect and report small variants such as indels and SNPs. SVGAP categorizes SVs according to their mutation class, including tandem duplications, transposable element insertions, or gene translocations, among others. This annotation adds valuable information to the detected SVs, enhancing the understanding of their potential biological impact.

To ensure the wide applicability and feasibility of SVGAP, we conducted thorough testing on a diverse range of commonly used aligners for whole genome alignments. We specifically included the aligners that exhibited excellent performance in the SVGAP pipeline, ensuring compatibility with genomes of varying complexity, particularly in plants. Through comprehensive benchmarking analyses using simulated structural variations (SVs) at individual, population, and phylogenetic levels, we show that SVGAP surpasses existing tools in terms of accuracy and completeness, regardless of SV type, sequence divergence, and genomic regions. Our benchmarking quantifies the accuracy of SV genotyping conducted by SVGAP. To evaluate SVGAP’s feasibility for real data, we applied it to 26 maize genomes, highlighting its proficiency in detecting SV at all genomic texts. The resulting VCF files can be used for pangenome construction and facilitate the genomic analysis of previously inaccessible genomic contexts. SVGAP is implemented in Perl and is freely available under the MIT license at [https://github.com/yiliao1022/SVGAP].

## Results

### Overview of SVGAP

The SVGAP pipeline consists of six main steps (**Fig. 1a**): (1) processing whole genome alignments (WGA); (2) constructing syntenic alignments; (3) calling SVs; (4) merging SVs from multiple samples; (5) re-genotyping SVs for each sample; and (6) annotating SVs.These steps are executed sequentially using a set of perl programs (**Fig. 1b**). During the first three steps, each sample is independently processed against the reference genome, though multiple samples can be parallelized for SVGAP. Here, we provide a brief overview of these steps, while additional details can be found in Methods.

**Fig. 1:**
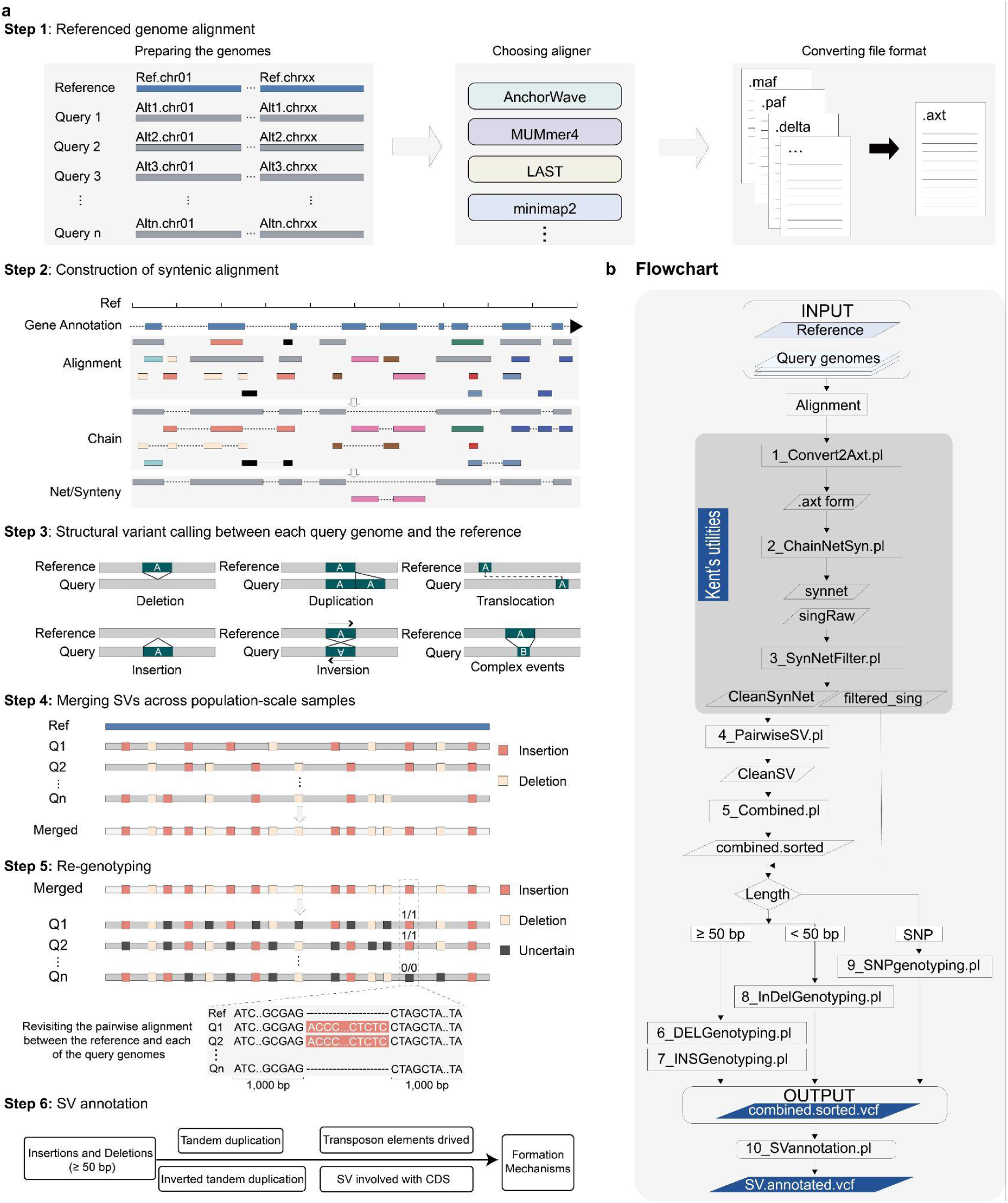
Overview of the SVGAP pipeline. **a**, The six main steps in the SVGAP pipeline: (1) converting WGA results from external alignment tools into the desired AXT format files; (2) employing Kent’s utilities to construct syntenic alignments at chromosome-scale level; (3) identifying SVs between each sample and the reference; (4) merging SVs across samples to generate a unique SV call set. (5) Re-genotyping SVs and producing the fully genotyped VCF files; and (6) annotating SVs for understanding mechanisms underlying their formation. **b**, The flowchart and Perl scripts implemented in each step of the SVGAP pipeline.

The inputs for SVGAP are whole genome pairwise alignments (WGA), including alignments produced by tools like last, MUMmer, minimap2, AnchorWave, and others. As a result, the first step of SVGAP involves converting the output from these tools into a common output format, such as the AXT format defined in UCSC (**Fig. 1a, step 1**). This ensures that SVGAP works with any aligner, provided the alignments can be converted into the required format. This flexibility allows users to choose their preferred aligner while still benefiting from the functionality and capabilities provided by SVGAP.

Detecting SVs from WGA poses two main challenges: distinguishing between orthologous and paralogous alignments, and identifying large SVs from fragmented alignments. SVGAP seeks to address these challenges by constructing alignment chains and nets using UCSC programs (53) (**Fig. 1a, step 2**). Chains represent co-linear local alignments in the same order between two genomes, while gaps within a chain likely indicate large insertions or deletions. Nets, on the other hand, are collections of the best chains hierarchically organized to cover genomic regions with single coverage. This dataframe enables the detection of large SVs and SVs in syntenic regions, even in rearranged genomic regions such as inversions and translocations (**Supplementary** Figure 1).

Using alignment nets, SVGAP can accurately identify various types of SVs. These include small insertions and deletions (<50 bp), large insertions and deletions (>50 bp), tandem duplications, inversions, translocations, and complex genomic loci where the reference and query genomes do not align (displaying a double-sided gap) (**Fig. 1a, step 3**). Additionally, SVGAP offers the flexibility to filter out paralogous alignments if desired, ensuring that only the top or most confident chains are used for SV calling. This filtering can be adjusted based on the complexity of the genomes involved.

As SVs are detected individually for each sample, SVGAP provides a merging function to generate a nonredundant SV call set (**Fig. 1a, step 4**). To identify putatively identical events, SVGAP first combines calls of the same SV type across samples and sorts them based on coordinates. Subsequently, different strategies are employed for each type of SV. For example, deletions and inversions are merged using an adjustable threshold for overlap (e.g., 90%). If the coordinates of two SVs overlap by at least that threshold, they are considered the same event and merged. In the case of insertion events, sequence identity is also taken into account in addition to coordinates. This ensures not only that the coordinates but also that the actual sequence of the inserted fragment is considered for identification and merging purposes.

After generating the non-redundant SV set, SVGAP proceeds to re-genotype each call across all samples using the corresponding filtered pairwise WGA (**Fig. 1a, step 5**). This involves genotyping each SV site in every sample by extracting and examining the local sequence alignments. SVGAP also offers a program to genotype SNVs using the filtered one-to-one alignment files. The outcome of this step is fully genotyped VCF (variant call format) files for each SV type, as well as for SNVs. These files are well-suited for further pangenome construction and evolutionary population genetics studies.

SVGAP also aims to annotate SVs by inferring the mechanisms underlying their formation (**Fig. 1a, step 6**). In other words, SVGAP has the capability of deducing various mutational mechanisms rather than simply identifying alignment gaps as indels. First, it can accurately identify an insertion or deletion event as a duplication or contraction derived from surrounding sequences. It does this by comparing the inserted or deleted sequences to their immediately flanking sequences. Second, SVGAP is proficient in recognizing an insertion event as a transposable element (TE) insertion, because it can compare the SV to TE sequence library. Third, it can determine if an insertion event is the result of gene duplication. This biologically motivated approach enhances accurate identification and characterization of larger insertions involving multiple TE elements.

We provide a detailed and easy-to-follow guide on how to use SVGAP for analyzing population-scale genomes in the **Supplementary Note 1**. In the following sections, we present our extensive benchmarking analyses, which serve as the foundation for constructing the SVGAP pipeline. We also compare SVGAP to existing tools and demonstrate its effectiveness in various scenarios.

### Assessing aligners for whole genome alignment of plant genomes

Although WGAs serve as input for assembly-based SV identification approaches, the consequences of alignment software choice have yet to see much attention, especially in the case of plants. To fill this gap, we conducted a preliminary assessment of 14 commonly used aligners by applying them to two rice (*Oryza sativa*) genomes (∼380 Mb), MH63 and ZS97 (54). We evaluated each aligner for various metrics, including computational speed, memory usage, and alignment quality. Based on these results, we retained six aligners for further consideration: lastz, last, MUMmer4, AnchorWave, GSAlign, and minimap2. Details for running these tools are described in **Supplementary Note 2**.

We next expanded the assessment of the six selected aligners to larger and more repetitive plant genomes (i.e. tomato, maize, and pepper), representing a diverse range of genome complexities (**Supplementary Table 1**). We also included fly (*Drosophila melanogaster*) and human (*Homo sapiens*) genomes for comparison. For each combination of species and aligner, we calculated pairwise alignments across a total of 11 aligner-parameter conditions (**Supplementary Table 2**). Figure 2 illustrates the running time, memory usage, storage space requirements, and alignable portion of the genome for each aligner across the tested species (see **Methods** for more details on evaluation metrics and **Supplementary Table 3** for tool execution details). To summarize their performance, GSAlign, AnchorWave, and MUMmer4 completed the alignments for all species using their default parameters. However, minimap2 exceeded system memory capacity (NGB 1 TB) when aligning the maize and pepper genomes. Additionally, lastz failed for these same two species as well as the human genome due to excessive runtime. To address the memory capacity issue encountered when running minimap2 when aligning the maize and pepper genomes, we employed a solution by dividing the query genome into smaller segments (e.g., using a 20 Mb window with a 2 Mb step) before performing the alignment against the reference genome. Additionally, when using its default parameters, Last generated a significant amount of raw alignments for the human, pepper, and maize genomes (1.0, 3.1, and 5.1 Tb, respectively), which poses a challenge for downstream analysis.

**Fig. 2:**
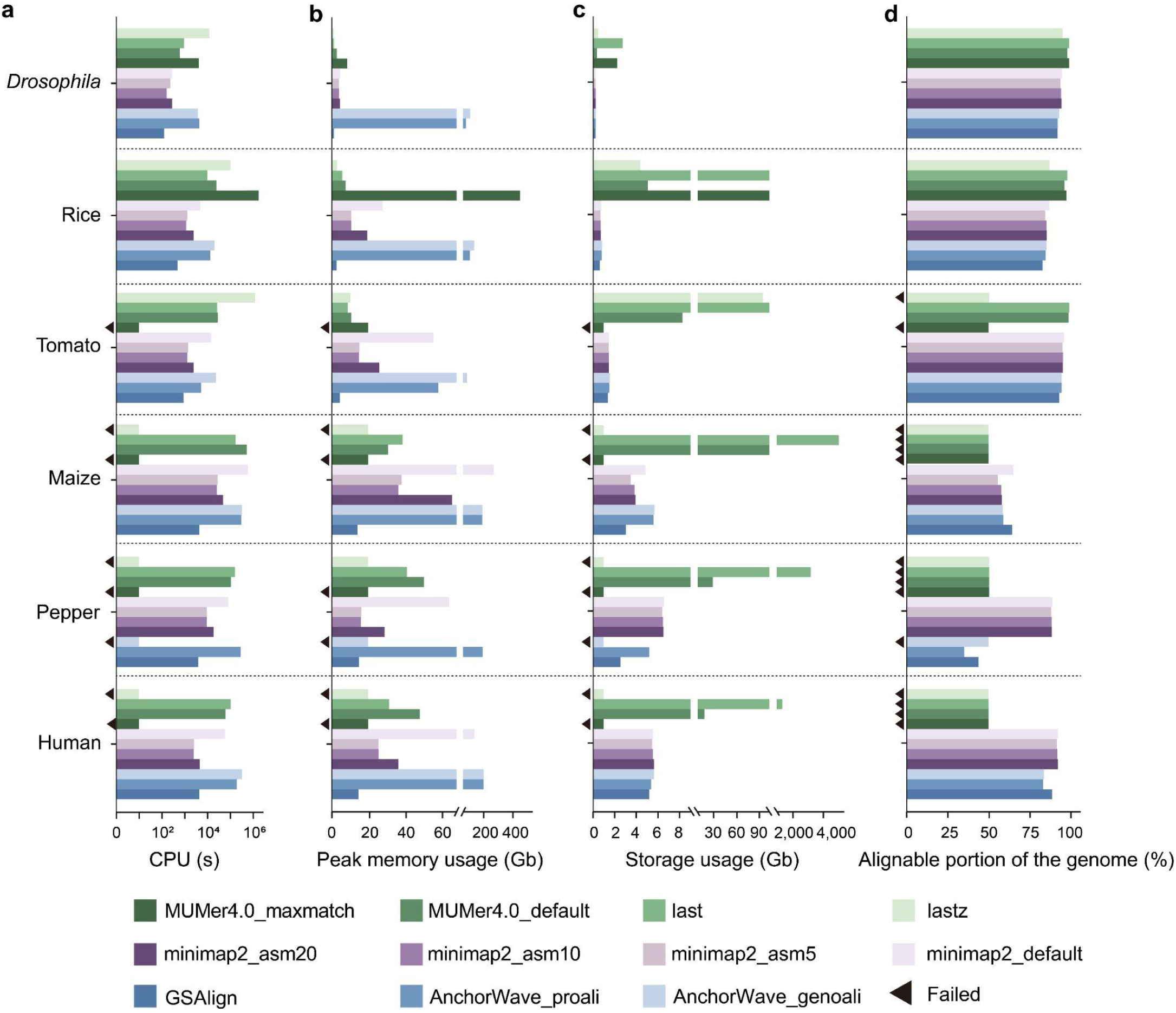
The performance of six widely used WGA tools on genomes of *Drosophila*, rice, tomato, maize, pepper, and human, representing varying levels of complexity. Metrics assessed include **a**. runtime, **b.** peak memory consumption, **c.** volume of raw alignments generated, and **d.** percent coverage of the reference. Each aligner’s performance was measured by aligning two representative genomes within each species as shown in **Supplementary Table 3**. Black triangles mark instances of failed alignment.

MUMmer4 only completed alignments for the *Drosophila* and rice genomes in the ’-maxmatch’ mode. AnchorWave encountered difficulties aligning the pepper genome in the ’-genoAli’ mode likely due to the fragmented genome assembly we used. Otherwise, all other alignments completed without issue and could be assessed. Overall, these results quantify the aligners’ capacity to align genomes, particularly in the context of plants with varying levels of genome complexity. They also highlight the key practical challenges in aligning plant genomes, because at least two species (maize and pepper) had low (<75%) alignable genome portions with most aligners. This information serves as a valuable guide for selecting aligners when developing *de novo* assembly-based methods for SV discovery.

### Performance of SVGAP across different aligners

We supplied the pairwise WGAs obtained from the six best aligners (i.e. AnchorWave, minimap2, Last, Lastz, MUMmer4, and GSAlign) to evaluate the performance of SVGAP by comparing the consistency and variability of SV calls across these aligners. Our analysis focused on deletions and insertions, which are the most common types of SVs, using genomes from *Drosophila*, rice, and tomato. For the rice genomes MH63 and ZH97 (54), SVGAP reported a wide range of SV calls (i.e., mutations ≥50 bp) across aligners, ranging from 3,735 to 11,642 for deletions and 3,821 to 11,542 for insertions. Among the aligners, AnchorWave reported the highest number of SV calls, followed by last, MUMmer4, minimap2, lastz, and GSAlign (**Supplementary Table 4**). The consistency of SV calls from pairwise comparisons varied between 27.4% and 92.5% (**Supplementary** Fig. 2a, for combined insertions and deletions). We used an upset plot (**Fig. 3a**, calculation details are provided in **Methods**) to visualize the consistency between different aligners. In total, these six aligners identified 28,548 unique insertions and deletions. Approximately 47.4% (13,542) of them were reported by at least five aligners, while nearly 23.6% (6,748) were reported by only one aligner. Among the aligner-specific SV calls, the proportion of AnchorWave calls was the highest, accounting for 12.78% (2,943/23,047) of its total calls. minimap2 reported 7.3% (1,455/19,940), Last reported 6.1% (1,274/20,774), Lastz reported 4.2% (786/18,679), GSAlign reported 1.6% (117/7,517), and MUMmer4 reported 1.1% (173/16,016). Notably, the differences among aligners may become more pronounced as genome complexity increases, as demonstrated by additional analyses using the *Drosophila* and tomato genomes (**Supplementary Table 4** and **Supplementary** Fig. 2b-e). This observation indicates that the choice of aligners significantly impacts SV detection.

**Fig 3:**
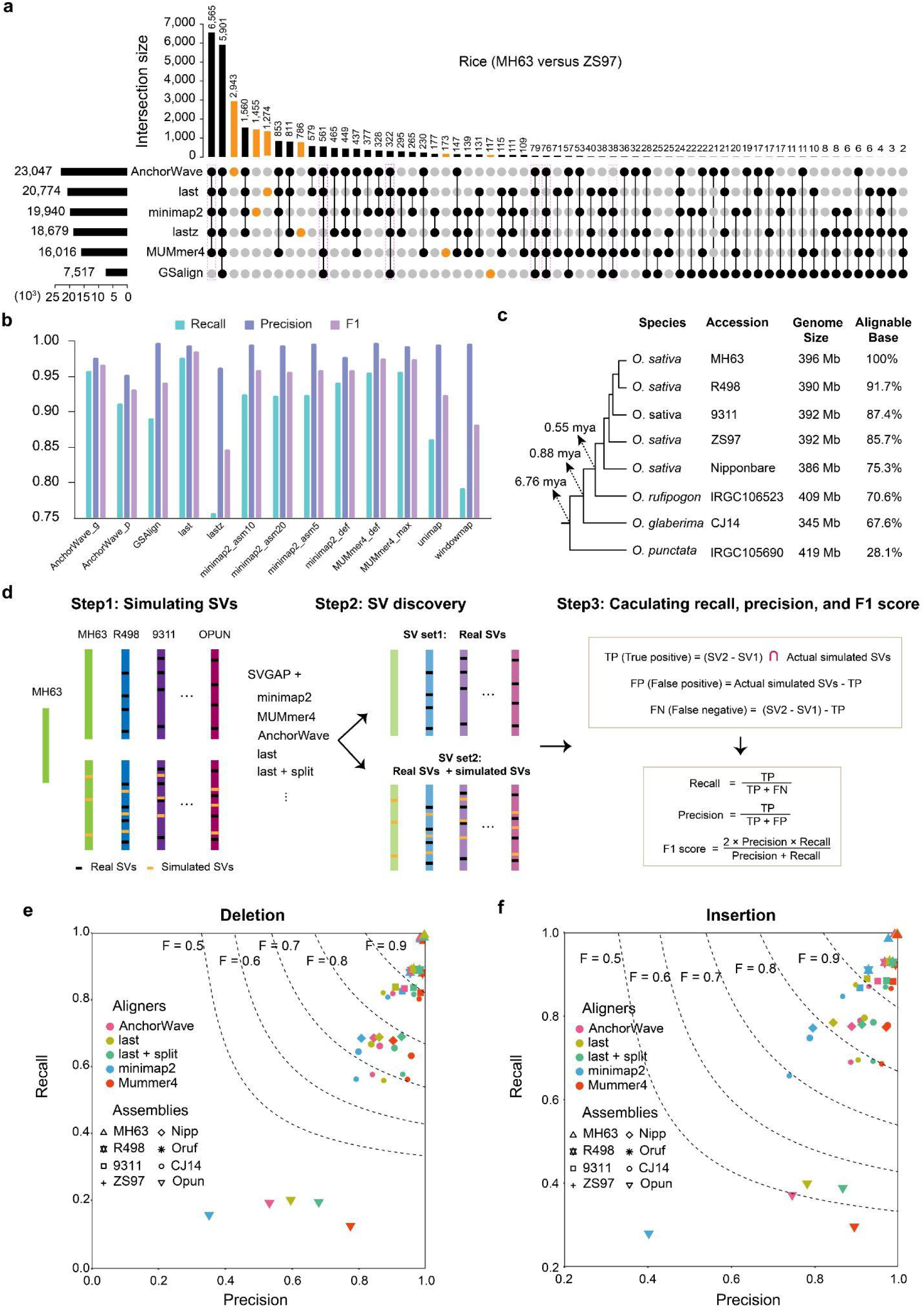
Performance of SVGAP for SV detection across different aligners. a, Analysis of shared SV calls—including deletions and insertions—among various aligners applied to two rice reference genomes. **b,** Comparative assessment of SVGAP performance in detecting SVs between a rice reference genome and its simulated counterpart, which includes introduced SVs. **c,** Phylogenetic relationships and estimated divergence times of selected rice (*Oryza*) genome assemblies, along with their approximate percentage of sequence alignment to the reference genome (MH63). **d,** Strategies for evaluating SVGAP performance across different aligners at varying levels of sequence divergence, along with the formulas used to calculate recall, precision, and F1 score. **e** and **f**, Comparison of SVGAP performance across aligners in detecting SVs, with panel **e** representing deletions and panel **f** insertions, at different levels of sequence divergence within the *Oryza* system.

To assess the performance of SVGAP on a validated truth set, we randomly simulated 7,264, 20,328, and 28,440 deletions and 4,290, 10,046, and 15,245 insertions (≥50 bp) using RSVSim (55) in the reference genomes of *Drosophila* (iso-1), rice (Nipponbare), and tomato (SL4), respectively (**Supplementary Table 5**). The reference genomes were then aligned with their respective simulated versions employing different aligners and parameter settings for SV detection. The resulting SV callsets were cross-referenced with the validated truth set to assess the precision of SV discovery by SVGAP in combination with the different aligners. These comparisons highlight that Last consistently outperformed all the other aligners, achieving the highest recall (97.5%-99.0%), precision (99.0%-99.7%), and F1-score (98.4%-99.4%) for both insertions and deletions (≥50 bp) across all three species (**Supplementary Table 5** and example shown for rice in **Fig. 3b**). While other callers also exhibited comparably high performance, their F1-scores ranged from 97.2% to 99.2% for MUMmer4, 95.1% to 98.1% for minimap2, 91.7% to 97.3% for AnchorWave, 93% to 97.1% for GSalign, and 79.7% to 95% for Lastz. Notably, Lastz showed the poorest performance in plant genomes, suggesting it may not be an ideal choice for SV discovery within the SVGAP workflow when analyzing large and repetitive plant genomes.

We further evaluated the performance of SVGAP across different aligners at various levels of sequence divergence, excluding Lastz and GSalign due to their inferior performance as previously demonstrated. To achieve this, we simulated SVs in genomes with varying phylogenetic distances from a reference genome. We then applied SVGAP with different aligners to identify SVs between each simulated genome and the reference, as well as between each original genome and the reference. The latter served as the background set of SVs, which were subsequently subtracted from the former. The resulting SV sets were used to calculate recall and precision, as illustrated in **Fig. 3c**. For instance, we used MH63 (*O. sativa*) as the reference rice genome and introduced 12,000 deletions and 8,000 insertions, with sizes ranging from 50 to 20,000 base pairs, across eight genome assemblies (see **Methods**). These assemblies included five from *O. sativa* (MH63, R498, 9311, ZS97, and Nipponbare), along with one assembly from each closely related species: *O. rufipogon*, *O. glaberrima* (CJ14), and *O. punctata*. Collectively, these represented an estimated divergence span of 6.78 million years (56), with the percentage of syntenic sequence alignment to the reference ranging from 28% to 100% (**Fig. 3d**; see **Methods** for calculation). Following the aforementioned strategy (**Fig. 3d**), we demonstrated that all tested aligners (last, MUMmer4, minimap2, and AnchroWave) successfully detected SVs across diverged genomes, yielding similar levels of recall and precision (**Supplementary Table 6, Fig. 3e**, **3f** and **Supplementary** Figure 3). Among these, Last and MUMmer4 consistently outperformed the other aligners, likely due to their specialized design for whole-genome alignments. Similar results were also observed in tomatoes (**Supplementary** Figure 4). These findings highlight the ability of SVGAP to accurately identify SVs between diverse and repetitive plant genomes using various aligners.

### Comparison of SVGAP with other genome assembly-based SV callers

We compared SVGAP (based on last alignments) to seven widely SV callers—Assemblytics (45), SyRI (43), SVIM-asm (47), paftools (49), GSAlign (50), MUM&Co (44) and AnchorWave (51)—all of which support the assembly-versus-assembly strategy for SV detection. To evaluate their performance on real data, we applied each tool to identify SVs between two rice genomes, MH63 and ZS97 (see **Supplementary Table 7** for parameters). For simplicity and consistency, we focused only on deletions and insertions, as these are the primary SV types detected by all tools. Among the evaluated methods, AnchorWave identified the highest number of SVs (24,960), followed by SVGAP (21,241), paftools (19,889), MUM&Co (19,514), SVIM-asm (13,402), Assemblytics (11,993), and SyRI (6,571), while GSAlign detected the fewest SVs (2,557)(**Supplementary Table 8**). In addition to large differences in the total number of calls, some methods exhibited distinct biases in SV size and type. For example, MUM&Co identified significantly more large SVs (>5 kb), whereas SyRI and GSAlign reported few, if any, SVs exceeding 1 kb (**Fig. 4a**). Furthermore, with the exception of AnchorWave, SVGAP, and paftools, all other tools showed a bias towards either deletions or insertions (**Fig. 4b**).

**Fig. 4:**
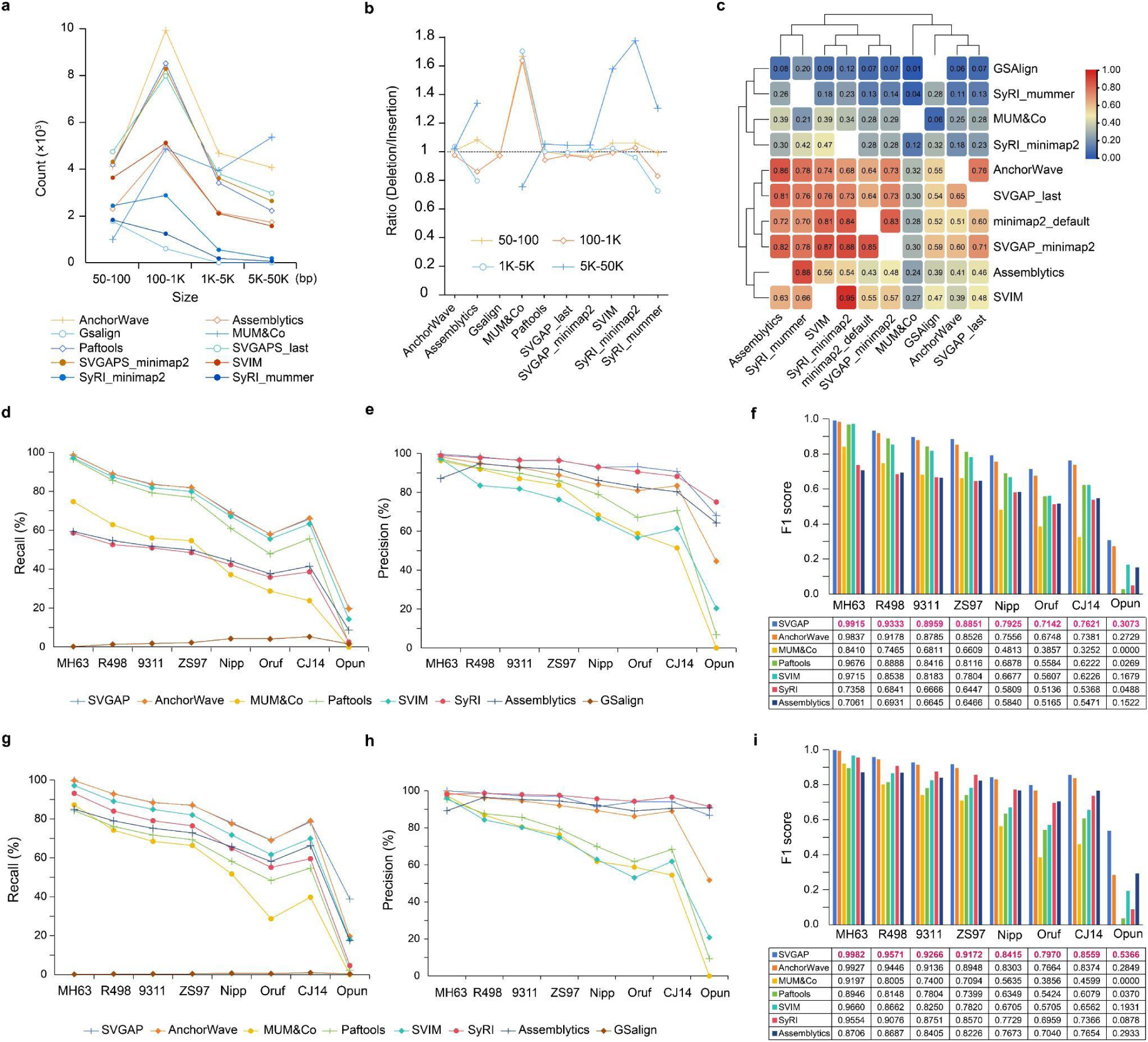
Performance comparison of SVGAP with other methods. a, Number of SV calls reported by different methods across various length ranges when comparing the two rice genomes, MH63 and ZS97. **b,** Ratio of deletions to insertions reported by different methods, again based on the two rice genomes. **c,** Overlap of SVs in pairwise comparisons among different methods. **d,** Recall for the benchmark analysis of deletions when comparing MH63 with 8 other divergent genomes, including simulated SVs. **e,** Precision of deletion detection. **f,** F1 score for deletions. **g,** Recall for the analysis of insertions. **h,** Precision of insertion detection. **i,** F1 score for insertions.

These results reveal substantial variability in detection efficiency and specificity across tools on real data.

We next evaluated the concordance and discrepancies of SV calls across different methods through calculating the overlap rate in pairwise comparisons. This overlap rate was defined as the proportion of SVs from one method that had a reciprocal overlap of at least 50% with the SV calls from another method. Our analysis revealed a wide range of concordance rates between pairs of methods, reflecting both shared and unique SV detections (**Fig. 4c**). Overall, AnchorWave, SVGAP, and paftools had higher overlaps with other methods and with each other, followed by SVIM and Assemblytics. In contrast, MUM&Co, GSAlign, and SyRI showed much lower overlaps with other tools, mainly due to their smaller number of SV calls. Similar results were also obtained in comparing two tomato genomes: SL4 and M82 (57) (**Supplementary** Figure 5a-c). These results reveal substantial variability in SV call reporting across different methods, highlighting the challenge of reconciling SV datasets generated by diverse tools.

To benchmark SVGAP for SV detection and compare its performance with other methods, we used the simulated rice and tomato genomes based on the reference in each species generated in the previous section. By comparing SV calls from each method to the simulated events, SVGAP consistently achieved the highest F1 scores across SV types and species, with at least 99.15% for rice and 99.53% for tomato (**Supplementary Table 6**).

To further evaluate the effect of sequence divergence on the performance of each tool, we applied them to detect SVs between the reference rice genome (MH63) and each of the simulated *Oryza* genomes representing varying levels of divergence used in the previous section (see **Fig. 3c** for details). This allowed us to systematically compare the effectiveness of each method in identifying SVs as sequence divergence increased. Since GSalign reported very few SVs during testing, it was excluded from further comparisons. The benchmark results demonstrated that SVGAP consistently achieved the highest F1 score for detecting both deletions (**Fig. 4d-f**) and insertions (**Fig. 4g-i**) across all divergent genomes, while also leading in recall and precision. AnchorWave achieved the second-best F1 scores. A similar trend was observed in tomato datasets (**Supplementary** Figure 5). These results indicate that SVGAP surpasses existing methods in SV discovery across a wide range of sequence divergence.

### SVGAP enables SV genotyping in large samples of assembled genomes

SVGAP independently identifies SVs for each individual sample. The SV calls from multiple samples are then merged to create a unique set of SVs by eliminating redundancy. However, this merged set lacks complete genotype information, as it only includes positive calls (indicating the presence of variants). To address this limitation, SVGAP provides tools for re-genotyping all SV calls across samples. This re-genotyping process examines pairwise alignments of each genome against the reference, extracting local sequence alignments around the SV sites—specifically focusing on regions extending a certain size (e.g. 2 kb) upstream and downstream of the breakpoints as defined by the user. By revisiting these local alignments, the tool infers the genotype for each SV in every sample, classifying them as positive, negative, or missing. More detailed descriptions of this process are depicted in **Supplementary** Figure 6.

To evaluate the merging and genotyping process, we benchmarked them using population-scale simulated genome assemblies in two plant species, rice and tomato. For rice, we simulated genomes by randomly incorporating SVs into the reference ‘Nipponbare’. These SVs were modified from a truth SV dataset (10,000 SVs) derived from comparisons of ‘Nipponbare’ with 48 different rice strains, including both insertions and deletions, as well as additional insertions resulting from transposable elements (see **Methods**). We simulated 20 genomes, each containing approximately half of the events from the truth SV dataset, along with one genome that included all the SVs (**Fig. 5a**).

**Fig. 5:**
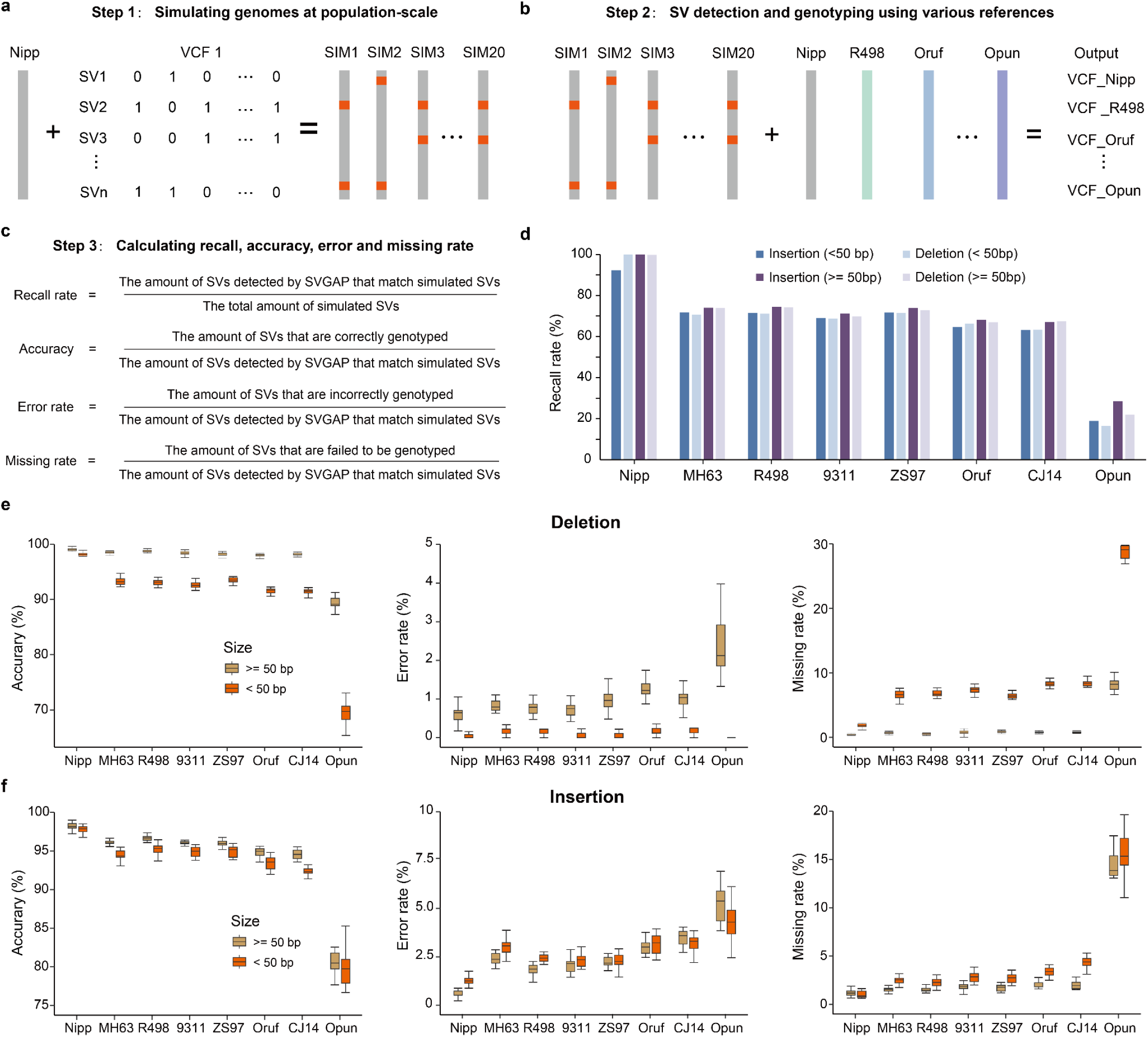
Benchmark analysis of merging and genotyping functions in SVGAP. a, Strategies for simulating population-scale genomes. **b,** SV discovery among population-scale genomes when using eight divergent genomes as the references. **c,** Formulas for computing recall rate, accuracy, error rate, and missing rate. **d,** Recall comparing the reference genome Nipponbare with eight other rice genomes across varying divergence scales, categorized by length. **e,** Accuracy, error rate, and missing rate for deletions. **f,** Accuracy, error rate, and missing rate for insertions across sequence divergence.

After applying SVGAP to identify SVs from these genomes relative to the reference ‘Nipponbare,’ the merging step successfully retrieved 99.8% (3,074 out of 3,079) of the total simulated SVs (> 50 bp), while producing only three false calls (**Fig. 5b** and **Supplementary Table 9**). The genotyping also achieved high accuracy, with an average of 99.0% of deletions and 98.2% of insertions correctly genotyped across samples. The error rate was only 0.5%, and the missing rate was 0.45% (**Supplementary Table 10**). These results demonstrate that SVGAP can accurately merge and genotype nearly all SVs while introducing few errors when comparing genomes with low levels of sequence divergence. Similar results were obtained in tomatoes (**Supplementary Table 9 and 10**).

To further assess the impact of sequence divergence on our merging and genotyping methods, we next identified SVs in the simulated population-scale genome assemblies using eight other divergent genomes as references (see previous section, **Fig. 3c**). As anticipated, the retrieval efficiency, or recall (as defined in **Fig. 5c**), declined as sequence divergence increases (**Fig. 5d**), due to the reduction of alignable regions between genomes. After applying a rough normalization based on the alignable rate between genomes (i.e., normalizing the recall divided by the estimated alignable proportion), SVGAP demonstrated the capability to identify the majority of simulated SVs (> 95%, **Supplementary Table 10**) located within the alignable regions. For genotyping, we focused only on the retrieved SVs and found that accuracy also decreased as sequence divergence increased, with a more pronounced effect on smaller events (< 50 bp) and insertions (**Fig. 5e** and **5f**). Except for *O. punctata* (whose genome has only roughly 28% of its sequence syntenically aligned to ‘Nipponbare’), SVGAP achieved 98.1% and 95.2% genotyping accuracy for deletions and insertions (> 50 bp) for all other references (**Supplementary Table 10**, **Fig. 5e** and **5f**). Accuracy for smaller SV events (< 50 bp) was slightly lower, with 91.2% for deletions and 93.3% for insertions. Correspondingly, the error and missing event rates increased with sequence divergence but remained generally low, with an approximate error rate of no more than 1% and a missing rate of 1.2% for deletions, and an error rate of 2.5% and a missing rate of 2% for insertions (see **Fig. 5e** and **5f**). Similar trends were also obtained in tomato genomes (**Supplementary Table 10** and **Supplementary** Figure 7). These analyses reveal that SVGAP can accurately merge and genotype SVs for population-scale genome assemblies with varying levels of sequence divergence.

### Discovering SVs in maize genomes and uncovering hidden genomic diversity in its centromeres

To assess the applicability of SVGAP in large and repetitive plant genomes, we applied it to 26 diverse maize genomes, including 25 NAM (nested association mapping) founder inbreds (58) and the reference B73 (14). The maize genome is known for its large size, approximately 2 Gb, with a significant proportion (∼85%) consisting of transposable elements (59). Our initial intraspecific comparison reveals that the average alignable portion in synteny between each query genome and the reference B73 accounts for approximately 56% of the entire genome (**Supplementary** Figure 8a and b). This suggests the existence of substantial presence and absence variants (PAVs) within maize genomes, which makes them an excellent model for evaluating the effectiveness of SVGAP.

The alignments obtained from minimap2, which were determined through benchmarking (**Supplementary Table 11** and **Supplementary** Figure 8c and d), were used in combination with SVGAP to detect SVs. By comparing each NAM genotype to B73 revealed a cumulative total of 1,758,685 SVs ≥50 bp in size, with count per genotype ranging between 61,451 and 75,913, except complex loci (**Fig. 6a** and **Supplementary Table 12**). Notably, SVGAP found a significant number of complex loci, where regions between conserved syntenic blocks cannot align to each other, implying rapid sequence turnover across the maize genome. This number of SVs is considerably higher than that reported in the previous study (60) that primarily used long-reads mapping methods to identify SVs. For deletion, approximately 90% of the deletions detected in previous work were also detected by SVGAP, whereas nearly half of deletions detected by SVGAP were not reported previously (**Supplementary** Figure 8e). Of those additional deletion calls detected by SVGAP, we found that a significant portion (45%) are independently detected in at least two genotypes, reflecting a substantial number of them representing false negatives based on the long-reads mapping methods used in the previous work. These missing calls were randomly distributed along chromosomes (**Supplementary** Figure 8f). For insertions, SVGAP detected 8.4-fold (28,009/235,410) more insertions than the previous work, probably reflecting that long read-mapping methods are significantly biased toward detecting deletions (26), but *de novo* assembly methods likely solve this issue. These results indicate that SVGAP can be applied to plant genomes as complex as in maize, providing a more comprehensive set of SVs that were not identified by long-read mapping methods.

**Fig. 6:**
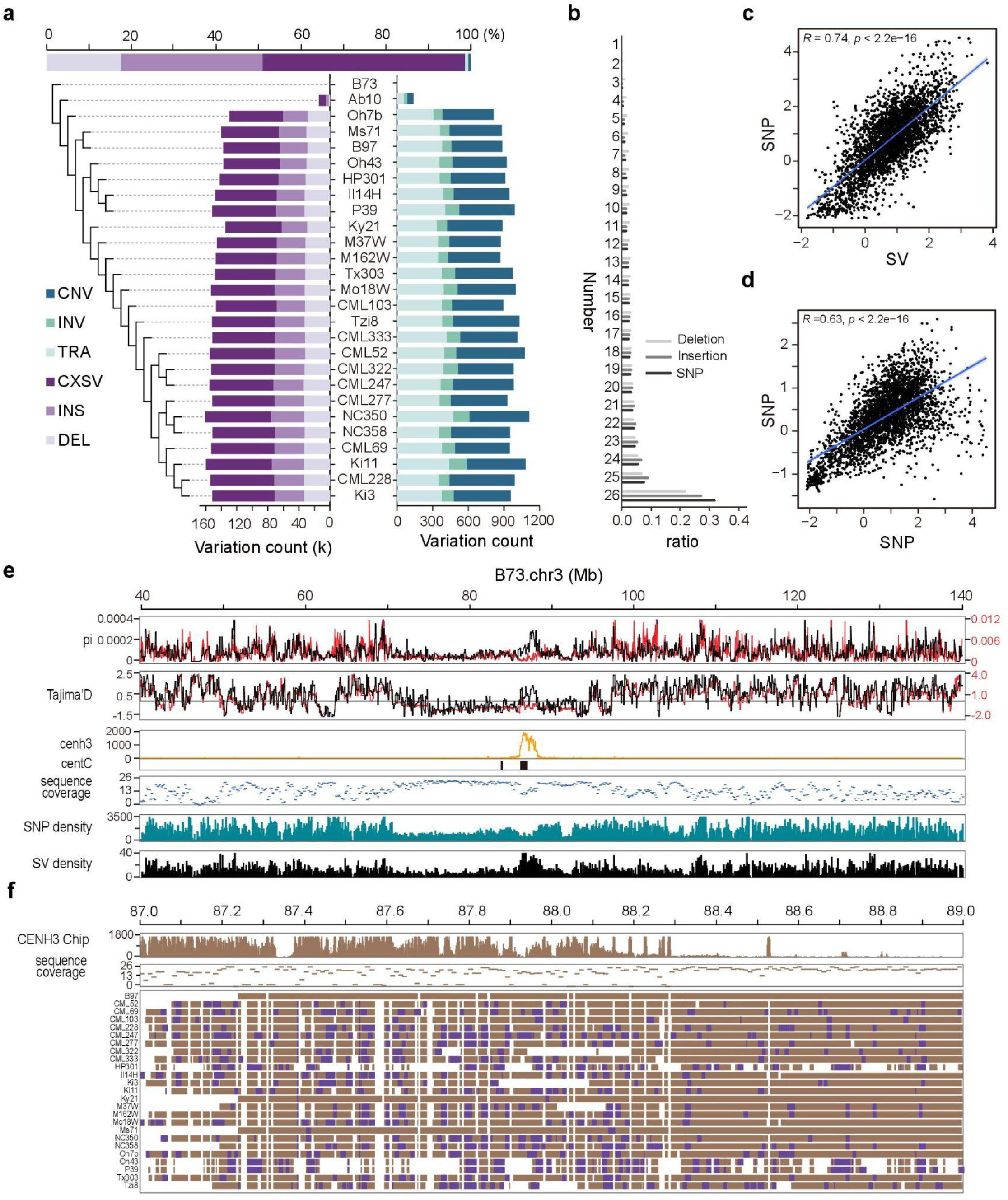
SV discovery in maize and hidden genomic diversity uncovered around its centromeric regions. **a**. SVs detected in 26 diverse maize genomes using SVGAP; **b**. Genotyping frequency for different types of genetic variations including insertion, deletion, and SNV across the samples; **c**. The plot graphs Tajima’s D calculated based on SNP and SV which detected by SVGAP across non-overlapping 100-kb windows of the maize genome, with the line indicating the correlation, which is strongly positive (Pearson r = 0.74, *P* = 2.2e-16); **d**. The plot graphs Tajima’s D calculated based on SNP detected by SVGAP in 26 maize genomes and SNP from a prior study, which identified SNP from 1,515 accessions across non-overlapping 100-kb windows of the maize genome. The line here too indicates a strong positive correlation (Pearson r = 0.63, *P* = 2.2e-16); **e**. Features of genetic variations and their population properties around the centromeric regions of chromosome 3. Panels from up to down indicate: the SVs (black line) and SNVs (red line) average pairwise diversity (π) for non-overlapping 100-kb windows, the SVs (black line) and SNVs (red line) Tajima’s D for non-overlapping 100-kb windows, CenH3 ChiP-seq reads mapping; distribution of maize centromeric specific tandem repeat (CentC), sequence coverage for non-overlapping 10-kb windows across the 26 maize genomes, SNV density, and SV density. **f.** The increased genomic diversity around the cen3 detected by SVs is attributable to the amplification of centromeric-specific retrotransposons.

Merging SVs across all lines revealed a total of 513,339 uniquely located SVs (excluding complex loci), including 192,455 deletions, 312,188 insertions, 2,084 inversions, and 6,612 copy number variations (**Fig. 6a**). We next used SVGAP to genotype deletions and insertions across all samples and generated the full genotyped VCF files for each. Genotyping results revealed that 52.3% of deletions and 63.3% of insertions can be genotyped in at least 20 lines, compared to those observed for SNPs (63.2%) and multiple sequence coverage (**Fig. 6b** and **Supplementary** Figure 8g and h). We next evaluated the quality of the genotyping result by examining the population properties of SVs.

Consistent with previous observations (14), both SVs and SNPs are more prone to occur at the chromosomal ends than middle centromeric regions (**Supplementary** Figure 9). Furthermore, the genomic diversity calculated with SV and SNP data consistently revealed a reduction of diversity across most centromeres, consistent with previous work (20). SV and SNP diversity were significantly correlated across chromosomal windows (*Pearson* r = 0.74, *P* < 2.2e-16; **Fig. 6c**). Also, the SNP diversity calculated with SVGAP data from the 25 diverse maize genomes was significantly correlated across chromosomal windows compared to those calculated with high-quality SNPs from 1,515 maize accessions (*Pearson* r = 0.63, *P* < 2.2e-16; **Fig. 6d**) (61). Together, these results imply that SVGAP can effectively genotype SVs across large samples of plant genomes, including complex genomes like maize.

While there was a substantial correlation between SNP and SV diversity across the majority of chromosomal windows, certain regions, particularly within some centromeres, displayed noticeable disparities (**Fig. 6e** and **Supplementary** Figure 9). Centromeres are among the most dynamic parts of the genome. Visually, centromeres and their flanking regions often display lower genomic diversity than chromosome arms in maize. Surprisingly, these regions contain highly conserved sequences in the population, which are reflected by the significantly higher sequence coverage (i.e. the number of lines that are alignable in synteny to B73 across the 25 maize lines (**Fig. 6e)**. We specifically focused on two centromeric regions (*Cen3* and *Cen9*) that show higher SV but not SNP diversity. The higher SV diversity observed in Cen3 (**Fig. 6f**) and Cen9 (**Supplementary** Figure 10) are the result of a higher occurrence of SVs in these genomic regions. By detailed examination of the SVs that occurred in these regions, we found most were insertions of CRMs (centromere-specific retrotransposons in maize). Of 459 SVs detected in the 2 Mb regions of the centromeres, 267 were CRM insertions (**Fig. 6f**). These results show that frequent insertions of CRM into the centromere regions can significantly increase local genomic diversity. They also demonstrate that SVGAP enables accurate SV detection in recalcitrant repetitive genomic regions and can therefore uncover hidden genomic diversity.

### Execution time and memory efficiency

To evaluate the runtime and memory usage of each step in the SVGAP pipeline, we conducted tests using 49 rice genomes on a high-performance system. Specifically, we utilized a dual CPU AMD EPYC 9654 96-core node running on Linux (Rocky Linux release 9.3), equipped with 1TB of DDR5 RAM and connected to storage through 1GB RAID controllers. After conducting the tests, we found that the SVGAP pipeline, which includes genome alignment, SV discovery, and genotyping steps, could be completed within two days on our system. Additionally, the peak memory usage of our method remained below 150 GB. **Supplementary Table 13** provides a summary of the computational resources, detailing the time and memory usage for each step of the SVGAP pipeline. These results indicate that SVGAP can efficiently leverage reasonable computational resources, making it suitable for routine SV discovery and genotyping in large assembly samples within a population or species.

## Discussion

Recent advancements in sequencing technologies and computational methods have made it possible to routinely generate nearly complete genome assemblies from large samples of the same or closely related species (40). These assemblies provide valuable opportunities to investigate the full spectrum of genetic variations and their functional impacts, especially previously inaccessible ones, across diverse evolutionary contexts and genomic landscapes (62, 63). Consequently, there is a growing demand for flexible and scalable computational tools to analyze population-scale genome assemblies, particularly in plant genomics, where the dynamic and plastic nature of plant genomes presents significant challenges for sequencing, assembly, alignment, and variant discovery (52, 64, 65).

In this study, we introduce SVGAP, a versatile pipeline for population-level identification of SVs from large samples of genome assemblies. SVGAP identifies SVs through pairwise whole-genome alignments, comparing multiple genome assemblies against a reference genome. The pipeline includes tools for combining SVs, genotyping a unique call set across all samples to produce fully genotyped VCF files, and annotating SVs (**Fig. 1**). Benchmark analyses using simulated plant genomes of varying complexities—spanning individual, population, and phylogenetic levels—demonstrated that SVGAP consistently outperforms existing tools in SV discovery (**Figs. 3** and **4**). It also provides accurate population genotyping and exhibits robustness against false variant discoveries (**Fig. 5**). We believe that the accuracy and comprehensive SV data generated by SVGAP make it a useful resource for constructing reference pangenome graphs (32, 66, 67). Although primarily tested on chromosome-scale assemblies within the same species, SVGAP can be easily adapted to accommodate assemblies of varying quality, size, and divergence.

An optimal whole-genome alignment is crucial for SVGAP and other similar SV callers. Selecting appropriate aligners for the species under examination is a prerequisite for ensuring the accuracy and comprehensiveness of SV discovery. Our initial assessment of existing aligners for large and repetitive plant genomes, such as maize and pepper, has uncovered significant challenges, because only a subset of aligners can effectively complete this task (**Fig. 2**). The practical obstacles encountered include long running times (e.g., Lastz and MUMmer4-default), high memory consumption (e.g., Minimap2 and Last-split), incomplete alignments (e.g., GSAlign), storage inefficiencies (e.g., Last-default), and difficulties in resolving specific chromosomal arrangements (e.g., AnchorWave). One approach for aligning large plant genomes involves splitting chromosomes into smaller segments and aligning these segments individually to the reference genome. We demonstrated this strategy in maize and pepper using minimap2 for alignment in the SVGAP pipeline. Our results, however, underscore the pressing need for new alignment tools optimized for plant genomes (52).

Comparing the overlaps of SV callsets from different callers or the same caller using different aligners reveals substantial inconsistencies. Genome complexity plays a key role, because the level of inconsistency increases with the complexity of the genome. Local sequence features are also critical, as we observed that inconsistent SV calls tend to overlap with repetitive regions and low-complexity sequences enriched in GA/AT dinucleotides. There is also an underlying biological complication, in that regions flanking indels demonstrate elevated mutation rates and sequence diversity that varies according to mating system and divergence times (68, 69). Although these boundary-specific dynamics were likely not captured in our simulation approach, our results nonetheless accentuated that a consistent choice of callers and aligners is essential for comparative analyses. Additionally, the efficiency of SV discovery across different genomic regions should be carefully considered in consensus-aware studies, such as comparisons between sex chromosomes and autosomes, or between centromeric regions and other genomic regions.

Genome divergence is an important factor to consider in SV discovery. Our benchmarking analyses indicate that detection recall and accuracy progressively decline as divergence increases, while the false discovery rate rises. This is expected, as genome divergence often leads to synteny decay due to large-scale rearrangements and gain/loss mutations. This restricts variant detection to regions where sequences remain alignable between genomes. Consequently, ancient and newly arising variants, such as SVs, in unalignable regions become increasingly difficult to detect. Furthermore, reduced sequence identity—driven by the accumulation of SNVs and InDels—introduces ambiguity in breakpoint detection for SVs within alignable regions as divergence increases. Employing sequence divergence-aware aligners may further enhance the accurate and reproducible detection of SVs across divergent genomes (49, 51).

While high-quality benchmark datasets with verified SVs exist for animals, comparable datasets for plants are still lacking. Given the distinct genomic features of plants, creating such datasets is crucial for evaluating existing SV callers and developing optimized ones, though this process is often challenging and time-consuming (70). In this study, we developed an efficient simulation approach to partially address this gap by simulating SVs in genomes with varying divergence from a reference, preserving the characteristics of sequence divergence. Our evaluation focuses on SVs simulated in alignable regions during genome comparisons, demonstrating the efficiency of this approach for evaluating aligners and callers. Further studies with high-quality benchmark datasets or novel simulation applications can improve our assessment of existing methods and aid in developing new ones.

Deciphering the mechanisms of SV formation is crucial after their discovery and can in turn guide the development of new tools. Understanding these mechanisms enhances SV detection accuracy, improves functional annotation, and offers insights into their evolutionary and biological significance. SVGAP has addressed simpler mechanisms like tandem duplications and TE or gene-related insertions, but further improvements are needed to resolve mechanisms that lead to more complex SV events, for example, nested TE insertions which are prevalent in plant genomes (71).

In maize we observed increased genomic diversity at maize centromeres, mainly due to the amplification of centromere-specific LTR-retrotransposons. This observation benefitted from previously assembled high-quality genomes, which allows SVGAP to enhance population genomic approaches and improve the interpretation of previously inaccessible genomic regions. The practical implications for maize suggest that future improvements to SVGAP should focus on increasing speed, accuracy, efficiency, scalability, and user-friendliness.

While the primary aim of SVGAP is to generate complete and accurate reference-based variant calls from large samples of high-quality genome assemblies, its output can also be valuable for pangenome construction. The resulting VCF files can be directly used to create VCF-derived pangenome graphs, such as those implemented in vg (32) and Paragraph (72), and for further genotyping with these tools. The strategy for whole genome alignment of large and repetitive plant genomes has potential implications for pangenome construction using tools based on whole-genome alignments or multiple genome alignments, such as pggb (73), PSVCP (42) and Minigraph-Cactus (66, 74, 75). Given the computational challenges posed by large, repetitive, and complex plant genomes, additional efforts may be needed to optimize these tools for plants.

## Methods

### Overview of SVGAP

SVGAP is designed to identify SVs (≥ 50 bp), InDels (< 50 bp), and SNVs from whole-genome pairwise assembly-assembly alignments between a reference genome and multiple query genomes. These alignments can be generated using tools such as minimap2 (49), MUMmer (76), Last (77, 78), AnchorWave (51), and others. SVGAP takes alignments from these external tools as input and follows a multi-step workflow consisting of six main steps (**Fig. 1**): (1) Alignment files are converted into the desired format for SVGAP; (2) Construction of syntenic and orthologous alignments; (3) SV discovering between each query genome and the reference; (4) Merging SVs across all pairwise comparisons; (5) SV re-genotyping by extracting local alignments from whole-genome alignments, and (6) SV annotation. The output of SVGAP consists of standard VCF files that contain confidently called SVs and other variants.

#### Conversion of alignment files

Alignment files from external aligners must be converted to AXT format (https://genome.ucsc.edu/) for downstream processing with SVGAP. MUMmer’s output *.delta* files are converted using the delta2maf (79) and mafToAxt (https://hgdownload.soe.ucsc.edu/admin/exe/linux.x86_64/) programs. Minimap2’s output *.paf* files are converted using paftools.js (distributed with Minimap2) and mafToAxt, with minor modifications required for the intermediate *.maf* files. Output *.maf* files from AnchorWave, Last, and Lastz are also converted using mafToAxt, with additional adjustments needed for compatibility. Other tools involved in the conversion process include maf_sort and maf_order from TBA (80), along with scripts developed for SVGAP in this study.

#### Construction of whole-genome synteny alignment

The converted alignment files were processed into structures known as chains and nets (https://genome.ucsc.edu/) with a series of programs developed by Kent et al. (53). Genome FASTA files were converted to 2bit format with faToTwoBit, and chromosome length files were generated using faSize with the -detailed parameter. Chains were constructed using axtChain with the -linearGap=medium option and filtered with chainPreNet. Nets were created using chainNet with the -minSpace=1 option and subsequently annotated with netSyntenic. The resulting *.synnet* files were converted to pairwise *.maf* files using netToAxt and axtToMaf, which were then used to generate *.maf* files with single coverage in both target and query genomes using single_cov2 from the TBA program (80).

#### Variant detection

SVs and InDels are identified for each query genome relative to the reference genome using the *.synnet* files with the program *PairASSYSV.pl*. These files can be further filtered with *SynNetFilter.pl* to enhance accuracy, although this may result in a slight loss of sensitivity. This step is particularly recommended when the compared genomes exhibit significant sequence divergence and rearrangements, ensuring that variants are identified only from reliable alignment portions, specifically syntenic and orthologous genomic regions, typically represented as top chains (the highest-scoring chained alignments). Gaps within the top chains may correspond to insertions, deletions, inversions, duplications, or translocations. Therefore, variants may also be retrieved from lower-scoring chains (i.e., secondary chains) if they fill large gaps in the top chains and indicate inversions or translocations (**Supplementary** Figure 1). SVGAP currently reports six types of SVs: insertions, deletions, tandem duplications, inversions, translocations, and complex events. Additionally, SNVs are jointly called from pairwise single-coverage *.maf* files using *SNPgenotyping.pl*. All aforementioned Perl scripts are included in SVGAP.

#### SV merging

SVGAP independently identifies SVs for each sample relative to the reference genome. Due to sequence divergence, the same SV event from different samples may exhibit slight differences in their breakpoint coordinates. To create a non-redundant SV dataset, SVGAP uses the program *Combined.pl* to merge SVs across samples. SVs from all samples are first combined for each type and sorted based on reference coordinates. For deletions, inversions, and translocations, events with identical coordinates or reciprocal overlap of at least the specified cutoff threshold (default 90%) are merged into a single event. For insertions and duplications, events with a breakpoint shift range of no more than 12 bp (or as defined by the user), inserted lengths varying by no more than 20%, and sequence identity above 50% (adjustable) are considered a single event.

#### SV re-genotyping

The merged SV dataset lacks complete genotyping information, as only samples where the event was reported have clear genotypes, while the status in samples that did not report the event remains uncertain. This uncertainty may arise from sequence gaps, sequence loss due to divergence, or genotypes identical to the reference. To address this issue and obtain a comprehensive genotype status across samples, SVGAP provides DELgenotyping.pl, INSgenotyping.pl, and InDelGenotyping.pl to conduct a second round of genotyping for deletions, insertions, and InDels, respectively (see **Supplementary** Figure 6). These tools extract and analyze the local alignment around each SV locus from pairwise whole genome alignments to infer the genotype status (e.g., reference type, alternative type, or missing data) and generate a standard VCF file for each SV type. For deletions and InDels, local alignments extracted from the single coverage .maf file are directly examined to verify the target event. For insertions, local sequences around the SV locus are first extracted from each query genome and realigned to the reference sequence using Stretcher (81). The resulting alignment is then used to infer the insertion event based on SV length and sequence identity.

#### SV annotation

To annotate SVs, users need to provide FASTA files for full-length coding DNA sequences (CDS) and intact transposable element (TE) sequences for the species under investigation. SVGAP extracts sequences of deletions and insertions from the VCF files and aligns them to these reference sequences using Minimap2, instead of directly comparing SVs with annotation files (e.g., GFF). This approach facilitates the identification of gene deletions or duplications, as well as TE insertions. Additionally, SVGAP compares each inserted sequence with its flanking regions to determine if the insertions result from local tandem duplications. All these analyses are performed by *SVannotation.pl*.

### Assessing and selecting aligners for plant whole-genome alignment

We selected 14 sequence aligners to evaluate their effectiveness in aligning plant genomes. These tools can be categorized into three groups: 1) aligners designed for genome-scale alignments, including Lastz [100], last (77, 78), MUMmer4 (76), GSAlign (50), and AnchorWave (51); 2) aligners for long-read mapping and alignment, such as minimap2 (49, 51), MECAT2 (82), Blasr (83), BWA-MEM [101], Ngmlr (84), and GrapMap (85); and 3) aligners optimized from minimap2 for specific targets, including Pbmm2 (https://github.com/PacificBiosciences/pbmm2/), Unimap (https://github.com/lh3/unimap), and winnowmap2 (86). To evaluate these aligners, we aligned two rice genomes, MH63 and ZS97 (54), using their default or recommended parameters. Their performance of these aligners is detailed in **Supplementary Note 2**.

Based on the evaluated results, we selected six aligners for further evaluation: Last, Lastz, minimap2, MUMmer4, GSAlign, and AnchorWave. These aligners were used to align two individual genomes from five additional species: human (CHM13 (87) *vs.* GRCh38 (88)), fruit fly (iso-1(89) *vs.* A4 (39, 89)), maize (B73 (90) *vs.* Nc350 (60)), tomato (SL4.0 (91) *vs.* M82 (57)), and pepper (Ca59 (92) *vs.* Zunla-1 (93)). In total, we tested 11 aligner-parameter combinations: minimap2 with four settings (-asm5, -asm10, -asm20, and default), MUMmer4 with two options (-maxmatch and default), AnchorWave with -genoAli and -proAli modes, and Last, Lastz, and GSAlign using their default configurations.

We evaluated the aligners across species based on runtime, peak memory usage, storage requirements for raw alignments, and the proportion of the reference genome covered by alignable regions. Runtime and peak memory usage were measured using the command ‘/usr/bin/time -v’. Storage requirements were calculated from the size of the *.maf* file, which was converted from the default output format generated by each aligner. The proportion of the reference genome covered by alignable regions was calculated from the single-coverage *.maf* files generated using the single_cov2 program. The results are summarized in **Supplementary Table 3**.

Specifically, SVGAP includes the program *SplitFa.pl*, which divides large chromosomes into smaller segments with user-defined sizes and step intervals. We applied this split-genome approach to large and repetitive plant genomes (e.g., maize and pepper) when testing minimap2, as it often encounters memory exhaustion issues. This strategy effectively reduces peak memory usage. The coordinates of the segments in their original genomic positions can later be restored using the *Convert2Axt.pl* program.

### SV simulation and generation of benchmark datasets

We created three benchmark datasets containing simulated SVs for both rice and tomato, providing a ground truth for comparing the accuracy of SV detection across different aligners and callers. The first dataset was generated by introducing SVs into the reference genomes of each species using RSVSim (55). For rice, we simulated 10,000 insertions (1 to 20 kb), 15,000 deletions (1 to 40 kb), 200 tandem duplications (200 bp to 20 kb), and 100 inversions (5 kb to 200 kb) based on the Nipponbare genome (94, 95). For tomato, we simulated 15,000 insertions, 20,000 deletions, 500 tandem duplications, and 100 inversions, with similar size ranges, using the SL4.0 genome assembly (91). We refer to these as **reference-based** simulated datasets.

The second dataset was generated by simulating SVs among genomes at varying levels of divergence, based on the phylogenies of *Oryza* and *Solanum*. We refer to these as **phylogeny-based** simulated datasets. For *Oryza*, we independently simulated 10,000 deletions (ranging from 50 bp to 20 kb) and 10,000 insertions (ranging from 50 bp to 40 kb) using RSVSim across eight phylogenetically divergent genomes, representing approximately 6.76 million years of divergence (56). These genomes include five from *Oryza sativa*: MH63 (54), R498 (12), 9311 (12), ZS97 (54), and Nipponbare (94), along with three from other species: *Oryza rufipogon* (IRGC106523) (96), *Oryza glaberrima* (CJ14) (12), and *Oryza punctata* (IRGC105690) (96). For *Solanum*, we also selected 8 phylogenetically divergent genomes and independently simulated 15,000 deletions and 15,000 insertions (ranging from 1 bp to 50 kb) in each one. These genomes include three from *S. lycopersicum*: SL4.0 (91), M82 (57), and ZY65 (63), as well as five from closely related species: *S.galapagense* (ZY56), *S. pimpinellifolium* (ZY57), *S. chmielewskii* (ZY60), *S. peruvianum* (ZY61), and *S. habrochaites* (ZY59) (63). These tomato genomes span approximately or less than 7.5 million years of divergence (63), with the proportion of sequence alignable to the reference SL4.0 ranging from 41.5% to 100% (**Supplementary** Figure 4a). This proportion was calculated from each single-coverage *.maf* file between the query genome and the SL4.0.

The third dataset was generated by simulating SVs in population-scale genome assemblies for rice and tomato. A program called *Simulator_pop.pl* was developed to introduce three types of SVs—deletions, insertions, and duplications—into the reference genomes, allowing for the simultaneous generation of multiple genomes. This program requires three inputs: a reference genome, a set of SVs, and the number of genomes you wish to generate. The set of SVs should include the following information: coordinates, genotypes across samples (where 0 indicates similarity to the reference and ‘1’ denotes an alternative), and sequences for the insertion events. We prepared the set of SVs for both rice and tomato using real datasets from previous studies (12, 92), with coordinates reassigned. For simplicity, these SVs were randomly selected from four chromosomes (Chr01, Chr03, Chr08, and Chr12 in rice; Chr01, Chr02, Chr03, and Chr04 in tomato), along with a subset of transposable elements-associated insertions. In total, there are 2,977 deletions, 2,232 insertions, and 139 duplications for rice, and 3,017 deletions, 3,382 insertions, and 152 duplications for tomato, with sizes ranging from 50 to 10,000 bp. We simulated 21 genomes for each species based on the reference genome Nipponbare for rice and SL4.0 for tomato. In each simulated genome, approximately half of the SVs are present, while one genome contains all the SV events. We refer to these as **population-based** simulated datasets.

### SV overlapping analysis

To assess the consistency of SV calls obtained from different aligners and callers, we compared the overlap of SV calls across various methods. We applied the following approach to generate a data frame for the UpSet plot (see **Fig. 3a**): 1) SVs (deletions and insertions) detected under different methods were combined and categorized into three size groups: less than 5 Kb, 5–10 Kb, and larger than 10 Kb, for each type; 2) SVs in each size group were merged independently using BEDTools (v2.30.0) to remove redundancy; 3) The three size groups were then combined to create a unified reference SV set; 4) SVs identified by each method were compared against the reference set to determine their overlap. An SV was considered a positive call if its coordinates had 100% overlap with any SV in the reference set. This approach provides a comprehensive list of unique SVs generated from all methods and their detectable status across those methods. The UpSet plot was generated using TBtools-II (97).

We also created a similarity matrix based on the pairwise overlap rate among methods. The pairwise overlap between methods was calculated as the proportion of SVs exhibiting at least 80% reciprocal overlap. We used the Jaccard index distance to generate cluster groups and assess the agreement (see **Fig. 4c**) between methods using TBtools-II (97).

### Benchmarking aligners for SV detection with SVGAP

Pairwise whole-genome alignments generated using AnchorWave (v1.0.1), Minimap2 (2.24-r1122), MUMmer (version 4), GSAlign (v1.0.22), Last (version 1406), Lastz (version 1.04.22), Unimap (0.1-r41), and Winnowmap (version 2.03), with default or recommended settings (**Supplementary Table 2**), were used as inputs for SVGAP to detect SVs. We benchmarked the aligners using the first two simulated datasets (see the section “SV Simulation and Generation of Benchmark Datasets”) for rice and tomato, as well as a reference-based simulated SV dataset for *Drosophila*. For tomato and *Drosophila*, we excluded Unimap and Winnowmap from comparison because their performance did not outperform Minimap2 in our analysis of rice.

The SV call sets obtained from SVGAP using different aligners were compared against the ground-truth SV set to assess recall, precision, and F1 score. An SV call (deletion or insertion) was considered a true positive (TP) if it exhibited at least 80% reciprocal overlap with the corresponding call in the ground-truth SV set. For the reference-based simulated dataset, SVs were identified between the simulated genome (denoted as A’) and its original reference genome (denoted as A). In contrast, for the phylogeny-based simulated dataset, two SV datasets were generated for each comparison. The first dataset was identified between the reference genome (A) and the original query genome (B), serving as the background. The second dataset was identified between the reference genome (A) and the simulated query genome (B’). The comparison SV dataset was derived by subtracting the first dataset from the second, which was then used to evaluate against the ground-truth SV set for calculating recall, precision, and F1 score. Detailed formulas for these calculations are shown in **Fig. 3d**, and the specific commands and scripts are provided in **Supplementary Note 2**.

### Benchmarking callers for SV detection

We benchmarked SVGAP alongside seven popular assembly-versus-assembly SV callers: AnchorWave (v1.0.1), SyRI (v1.6), SVIM-asm (version 1.0.3), Assemblytics (https://github.com/MariaNattestad/Assemblytics), GSAlign (v1.0.22), MUM&Co (v3.8), and paftools (used with minimap2 2.24-r1122) for SV detection, utilizing the same benchmark simulated SV datasets previously employed for benchmarking callers in both rice and tomato. AnchorWave was executed with the ‘-v’ option to generate SV calls. For SyRI, alignments from MUMmer4 and minimap2 were employed to call SVs. SVIM-asm utilized alignments from minimap2 with the options “-a -x asm5 --cs -r2k” for SV calling. GSAlign was used for SV calling based on its alignment process.

Assemblytics relied on alignments from MUMmer4.0 with the options “--maxmatch” and default settings. For MUM&Co, alignments from MUMmer4.0 with default options were used for SV calls. Paftools was invoked alongside minimap2. All methods were executed with default or recommended parameters, as detailed in **Supplementary Table 7**. The resulting SV call sets were compared against the ground-truth SV set to calculate recall, precision, and F1 score using the methods described in the previous section.

### Benchmarking SV genotyping for population-scale genome assemblies

In order to evaluate SVGAP’s efficacy in SV genotyping for population samples, we carried out benchmarking tests on genome assemblies from both rice and tomato. Considering SVGAP employs two distinct procedures to manage this task - the first merges SVs from all individuals to produce a unique set of population-specific SV calls, and the second re-genotypes each call separately for each individual by revisiting the whole genome alignment between the respective individual and the reference - we conducted separate assessments for each of these steps. Prior to that, we simulated a set of population-scale genome assemblies (i.e. 21 individuals) for each of the rice and tomato, by introducing three types (that is, deletion, insertion, and duplication) of SVs into their reference genomes (Nipponbare for rice, and SL4 for tomato). The following processes were applied to complete this task. First, a list of SVs with coordinates and affected sequences extracted from the real SV datasets derived from population samples from both rice (this study) and tomato (98) were generated. The coordinates were randomly placed. These SVs include 2,977 deletions, 2,232 insertions, and 139 duplications for rice with size range between 50 and 10,000 bp, and 3,017 deletions, 3,382 insertions, and 152 duplications for tomatoes with size range between 50 and 10,000 bp. For simility, these SVs were randomly distributed across four chromosomes (Chr01, Chr03, Chr08, and Chr12 in rice, and Chr01-Chr04 in tomato) of the reference genome for each species. Second, a ‘10’ matric was constructed, in which rows represent SV calls, and columns represent the genotypes of individuals. In each column, the number of ‘1’ or ‘0’ (i.e. genotypes) are ensured to be roughly similar so as to make sure each individual can harbor roughly half of the SV calls. Third, a perl script named *PopGsimulate.pl* was developed to simulate population-scale genome assemblies with input of the ‘10’ matrix and a reference genome. A simulated VCF file was simulated based on the real SV spectrum of the real dataset.

The simulated genome assemblies constructed at the population scale were then individually mapped onto the reference genome, along with seven other divergent genomes, exhibiting varying levels of phylogenetic distance, as previously described. This was accomplished using MUMmer4 with its default parameters. These alignments were input into SVGAP for detecting structural variations (SVs) between each simulated individual and their corresponding reference genomes. SVGAP then combines SVs from all individuals to produce a unique set of SVs. The SV callset contains not only the simulated SVs but also a set of pre-existing SVs between the reference and other genomes, which should be filtered out. Any SVs in the callset with a genotype frequency greater than 19/21 were considered real SVs and were excluded, except when the assemblies were mapped to the reference where they originated. Post filtration, any called SVs with a simulated target match (i.e., minimum reciprocal overlap of 50%) were classified as true positives (TP), while those without a match were deemed false positives (FP). Evaluation metrics used to measure the first combination step included: completeness, calculated as the percentage of TP divided by the total number of simulated SVs times 100%; accuracy, calculated as the percentage of TP divided by the total number of all predicted SVs; and error rate, calculated as the percentage of FP divided by the total number of all predicted SVs.

For the true positive SV calls, we performed an assessment of genotyping accuracy, error rate, and missing rate for each individual. Accuracy was quantified as the percentage of correctly genotyped SVs out of the total reported SVs. Similarly, the error rate represented the percentage of incorrectly genotyped SVs among the total reported SVs. Additionally, the missing rate was determined as the percentage of ungenotyped SVs compared to the total reported SVs. More details are described in **Supplementary Methods**.

### SV discovery using SVGAP in maize genomes and subsequent analysis

Benchmarks for SV identification with different aligners were first conducted with simulated SVs. We used B73 (version 5) as the reference genome, while the other 25 NAM lines and the A10 assembly served as query genomes for SV identification. Each query genome was initially split into 20 Mb segments with a 2 Mb overlap using the ***SplitFa.pl*** program. These segments were then aligned to the B73 genome using minimap2 (2.24-r1122) with default settings to minimize memory usage. The coordinates of the segments were converted back to their original genomic coordinates using ***Convert2Axt.pl***. Following the SVGAP workflow, we called SVs and other genomic variants. The output includes VCF files for deletions and insertions (greater than 50 bp) as well as SNVs.

The VCF files were used to calculate the genotyping rate across samples for each variant, indicating how many samples could be genotyped for that variant. For each variant, the number of samples with a valid genotype (i.e., 1/1 or 0/0 in the corresponding VCF file) was counted. Tajima’s D and nucleotide diversity (π) were calculated in non-overlapping 100-kb windows along the chromosomes based on VCF files for SVs and SNVs using VCFtools (v0.1.16). An additional SNP VCF file from 1, 515 maize accessions was used for comparison, which was obtained from a previous study (61) and downloaded from https://www.maizegdb.org/. SNP and SV densities were also calculated for non-overlapping 100-kb windows along the chromosomes. Sequence-alignable coverage was defined as the number of query genomes aligned to the B73 reference genome over at least half the length of each fixed window. This metric was calculated for each 10-kb window from the single-coverage .maf file for each sample.

Maize CENH3 ChIP-seq data from a previous study (20) was downloaded from GenBank (accession no. SRP067358). The Illumina paired-end reads were mapped to the B73 reference genome (version 5) using Bowtie with the following parameters: -X 2000 --chunkmbs 3000 -k 3 --strata --best -v 2 -q.

### Runtime and memory requirements

We evaluated each step of the SVGAP pipeline using 49 rice genomes (12, 54, 99) on a high-performance computing system. The system consisted of a dual-CPU AMD EPYC 9654 96-core node running Rocky Linux 9.3, equipped with 1 TB of DDR5 RAM and connected to storage via 1 GB RAID controllers. Runtime and memory usage were monitored using the Linux time -v command.

Genome alignment was performed with MUMmer4.0 using default settings.

## Data availability

All data used in this study are publicly available and cited in the appropriate places where mentioned.

## Code availability

SVGAP and user manuals are publicly available at http://github.com/yiliao1022/SVGAP under the MIT License. We used the v1.0 version for SV discovery and benchmark in the manuscript. Key custom Perl scripts used in the manuscript can be found at http://github.com/yiliao1022/SVGAP/Utils.

## Acknowledgements

This work was funded by the GuangDong Basic and Applied Basic Research Foundation (Grant No. 2024A1515010362) and Research Start-up Funding from South China Agricultural University to Yi Liao. J.J. Emerson was partly supported by National Institutes of Health (NIH) awards R01GM123303 and R35GM153327. Mahul Chakraborty was partly supported by NIH award R00GM129411 and start-up funding from Texas A&M University. Chengjie Chen was partly supported by the National Science Foundation of China (Grant No. 32102320), the Invigorate the Seed Industry of Guangdong Province (2023-NJS-00-012), and the Central Public-interest Scientific Institution Basal Research Fund for the Chinese Academy of Tropical Agricultural Sciences (1630032024026). Jinfeng Chen was partly supported by the National Natural Science Foundation of China (Grant No. 32470607).

## Author contributions

Y.L. and J.J.E. conceived the project; Y.L. developed the software; M.H., C.C., and S.T. revised the software; Y.L., M.H., P.W., C.C., J.H.C., and L.W. tested the software and analyzed the data; Y.L. wrote the original manuscript; Y.L., M.H., P.W., C.C., J.F.C., M.C., B.S.G., and J.J.E. revised the manuscript. All authors read and approved the final version of the manuscript.

## Additional information

Supplementary material is included with the main text.

## References

1. Alkan, C., Coe, B.P. and Eichler, E.E. (2011) Genome structural variation discovery and genotyping. Nat. Rev. Genet., 12, 363–376.

2. Mahmoud, M., Gobet, N., Cruz-Dávalos, D.I., Mounier, N., Dessimoz, C. and Sedlazeck, F.J. (2019) Structural variant calling: the long and the short of it. Genome Biol., 20, 246.

3. Gaut, B.S., Seymour, D.K., Liu, Q. and Zhou, Y. (2018) Demography and its effects on genomic variation in crop domestication. Nat Plants, 4, 512–520.

4. Escaramís, G., Docampo, E. and Rabionet, R. (2015) A decade of structural variants: description, history and methods to detect structural variation. Brief. Funct. Genomics, 14, 305–314.

5. Li, Y., Roberts, N.D., Wala, J.A., Shapira, O., Schumacher, S.E., Kumar, K., Khurana, E., Waszak, S., Korbel, J.O., Haber, J.E., et al. (2020) Patterns of somatic structural variation in human cancer genomes. Nature, 578, 112–121.

6. Hadi, K., Yao, X., Behr, J.M., Deshpande, A., Xanthopoulakis, C., Tian, H., Kudman, S., Rosiene, J., Darmofal, M., DeRose, J., et al. (2020) Distinct Classes of Complex Structural Variation Uncovered across Thousands of Cancer Genome Graphs. Cell, 183, 197–210.e32.

7. Bridges, C.B. (1936) THE BAR ‘GENE’ A DUPLICATION. Science, 83, 210–211.

8. Sturtevant, A.H. (1913) The linear arrangement of six sex? linked factors in Drosophila, as shown by their mode of association. J. Exp. Zool., 14, 43–59.

9. Kou, Y., Liao, Y., Toivainen, T., Lv, Y., Tian, X., Emerson, J.J., Gaut, B.S. and Zhou, Y. (2020) Evolutionary Genomics of Structural Variation in Asian Rice (Oryza sativa) Domestication. Molecular Biology and Evolution, 10.1093/molbev/msaa185.

10. Saxena, R.K., Edwards, D. and Varshney, R.K. (2014) Structural variations in plant genomes. Brief. Funct. Genomics, 13, 296–307.

11. Yuan, Y., Bayer, P.E., Batley, J. and Edwards, D. (2021) Current status of structural variation studies in plants. Plant Biotechnol. J., 19, 2153–2163.

12. Qin, P., Lu, H., Du, H., Wang, H., Chen, W., Chen, Z., He, Q., Ou, S., Zhang, H., Li, X., et al. (2021) Pan-genome analysis of 33 genetically diverse rice accessions reveals hidden genomic variations. Cell, 10.1016/j.cell.2021.04.046.

13. Chen, J., Liu, Y., Liu, M., Guo, W., Wang, Y., He, Q., Chen, W., Liao, Y., Zhang, W., Gao, Y., et al. (2023) Pangenome analysis reveals genomic variations associated with domestication traits in broomcorn millet. Nat. Genet., 55, 2243–2254.

14. Hufford, M.B., Seetharam, A.S., Woodhouse, M.R., Chougule, K.M., Ou, S., Liu, J., Ricci, W.A., Guo, T., Olson, A., Qiu, Y., et al. (2021) De novo assembly, annotation, and comparative analysis of 26 diverse maize genomes. bioRxiv, 10.1101/2021.01.14.426684.

15. Liu, Y., Du, H., Li, P., Shen, Y., Peng, H., Liu, S., Zhou, G.-A., Zhang, H., Liu, Z., Shi, M., et al. (2020) Pan-Genome of Wild and Cultivated Soybeans. Cell, 182, 162–176.e13.

16. Zhou, Y., Zhang, Z., Bao, Z., Li, H., Lyu, Y., Zan, Y., Wu, Y., Cheng, L., Fang, Y., Wu, K., et al. (2022) Graph pangenome captures missing heritability and empowers tomato breeding. Nature, 606, 527–534.

17. Chen, S., Wang, P., Kong, W., Chai, K., Zhang, S., Yu, J., Wang, Y., Jiang, M., Lei, W., Chen, X., et al. (2023) Gene mining and genomics-assisted breeding empowered by the pangenome of tea plant Camellia sinensis. Nat Plants, 10.1038/s41477-023-01565-z.

18. Carvalho, C.M.B. and Lupski, J.R. (2016) Mechanisms underlying structural variant formation in genomic disorders. Nat. Rev. Genet., 17, 224–238.

19. Stuart, K.C., Edwards, R.J., Sherwin, W.B. and Rollins, L.A. (2023) Contrasting Patterns of Single Nucleotide Polymorphisms and Structural Variation Across Multiple Invasions. Mol. Biol. Evol., 40.

20. Schneider, K.L., Xie, Z., Wolfgruber, T.K. and Presting, G.G. (2016) Inbreeding drives maize centromere evolution. Proc. Natl. Acad. Sci. U. S. A., 113, E987–96.

21. Li, S., Lin, D., Zhang, Y., Deng, M., Chen, Y., Lv, B., Li, B., Lei, Y., Wang, Y., Zhao, L., et al. (2022) Genome-edited powdery mildew resistance in wheat without growth penalties. Nature, 602, 455–460.

22. Gao, L., Gonda, I., Sun, H., Ma, Q., Bao, K., Tieman, D.M., Burzynski-Chang, E.A., Fish, T.L., Stromberg, K.A., Sacks, G.L., et al. (2019) The tomato pan-genome uncovers new genes and a rare allele regulating fruit flavor. Nat. Genet., 51, 1044–1051.

23. Fuentes, R.R., Chebotarov, D., Duitama, J., Smith, S., De la Hoz, J.F., Mohiyuddin, M., Wing, R.A., McNally, K.L., Tatarinova, T., Grigoriev, A., et al. (2019) Structural variants in 3000 rice genomes. Genome Res., 29, 870–880.

24. Hämälä, T., Wafula, E.K., Guiltinan, M.J., Ralph, P.E., dePamphilis, C.W. and Tiffin, P. (2021) Genomic structural variants constrain and facilitate adaptation in natural populations of, the chocolate tree. Proc. Natl. Acad. Sci. U. S. A., 118.

25. Kosugi, S., Momozawa, Y., Liu, X., Terao, C., Kubo, M. and Kamatani, Y. (2019) Comprehensive evaluation of structural variation detection algorithms for whole genome sequencing. Genome Biol., 20, 117.

26. Ahsan, M.U., Liu, Q., Perdomo, J.E., Fang, L. and Wang, K. (2023) A survey of algorithms for the detection of genomic structural variants from long-read sequencing data. Nat. Methods, 20, 1143–1158.

27. Liu, Z., Xie, Z. and Li, M. (2024) Comprehensive and deep evaluation of structural variation detection pipelines with third-generation sequencing data. Genome Biol., 25, 188.

28. Cameron, D.L., Di Stefano, L. and Papenfuss, A.T. (2019) Comprehensive evaluation and characterisation of short read general-purpose structural variant calling software. Nat. Commun., 10, 3240.

29. Cleal, K. and Baird, D.M. (2022) Dysgu: efficient structural variant calling using short or long reads. Nucleic Acids Res., 50, e53.

30. Dierckxsens, N., Li, T., Vermeesch, J.R. and Xie, Z. (2021) A benchmark of structural variation detection by long reads through a realistic simulated model. Genome Biol., 22, 342.

31. Chin, C.-S., Behera, S., Khalak, A., Sedlazeck, F.J., Sudmant, P.H., Wagner, J. and Zook, J.M. (2023) Multiscale analysis of pangenomes enables improved representation of genomic diversity for repetitive and clinically relevant genes. Nat. Methods, 20, 1213–1221.

32. Hickey, G., Heller, D., Monlong, J., Sibbesen, J.A., Sirén, J., Eizenga, J., Dawson, E.T., Garrison, E., Novak, A.M. and Paten, B. (2020) Genotyping structural variants in pangenome graphs using the vg toolkit. Genome Biol., 21, 35.

33. Popic, V., Rohlicek, C., Cunial, F., Hajirasouliha, I., Meleshko, D., Garimella, K. and Maheshwari, A. (2023) Cue: a deep-learning framework for structural variant discovery and genotyping. Nat. Methods, 20, 559–568.

34. Lin, J., Wang, S., Audano, P.A., Meng, D., Flores, J.I., Kosters, W., Yang, X., Jia, P., Marschall, T., Beck, C.R., et al. (2022) SVision: a deep learning approach to resolve complex structural variants. Nat. Methods, 19, 1230–1233.

35. Denti, L., Khorsand, P., Bonizzoni, P., Hormozdiari, F. and Chikhi, R. (2023) SVDSS: structural variation discovery in hard-to-call genomic regions using sample-specific strings from accurate long reads. Nat. Methods, 20, 550–558.

36. Chen, Y., Wang, A.Y., Barkley, C.A., Zhang, Y., Zhao, X., Gao, M., Edmonds, M.D. and Chong, Z. (2023) Deciphering the exact breakpoints of structural variations using long sequencing reads with DeBreak. Nat. Commun., 14, 1–12.

37. Ebert, P., Audano, P.A., Zhu, Q., Rodriguez-Martin, B., Porubsky, D., Bonder, M.J., Sulovari, A., Ebler, J., Zhou, W., Serra Mari, R., et al. (2021) Haplotype-resolved diverse human genomes and integrated analysis of structural variation. Science, 372.

38. Logsdon, G.A., Rozanski, A.N., Ryabov, F., Potapova, T., Shepelev, V.A., Catacchio, C.R., Porubsky, D., Mao, Y., Yoo, D., Rautiainen, M., et al. (2024) The variation and evolution of complete human centromeres. Nature, 629, 136–145.

39. Chakraborty, M., VanKuren, N.W., Zhao, R., Zhang, X., Kalsow, S. and Emerson, J.J. (2018) Hidden genetic variation shapes the structure of functional elements in Drosophila. Nat. Genet., 50, 20–25.

40. De Coster, W., Weissensteiner, M.H. and Sedlazeck, F.J. (2021) Towards population-scale long-read sequencing. Nat. Rev. Genet., 22, 572–587.

41. Cochetel, N., Minio, A., Guarracino, A., Garcia, J.F., Figueroa-Balderas, R., Massonnet, M., Kasuga, T., Londo, J.P., Garrison, E., Gaut, B.S., et al. (2023) A super-pangenome of the North American wild grape species. Genome Biol, 24, 290.

42. Wang, J., Yang, W., Zhang, S., Hu, H., Yuan, Y., Dong, J., Chen, L., Ma, Y., Yang, T., Zhou, L., et al. (2023) A pangenome analysis pipeline provides insights into functional gene identification in rice. Genome Biol., 24, 19.

43. Goel, M., Sun, H., Jiao, W.-B. and Schneeberger, K. (2019) SyRI: finding genomic rearrangements and local sequence differences from whole-genome assemblies. Genome Biol., 20, 277.

44. O’Donnell, S. and Fischer, G. (2020) MUM&Co: accurate detection of all SV types through whole-genome alignment. Bioinformatics, 36, 3242–3243.

45. Nattestad, M. and Schatz, M.C. (2016) Assemblytics: a web analytics tool for the detection of variants from an assembly. Bioinformatics, 32, 3021–3023.

46. Li, H., Bloom, J.M., Farjoun, Y., Fleharty, M., Gauthier, L., Neale, B. and MacArthur, D. (2018) A synthetic-diploid benchmark for accurate variant-calling evaluation. Nat. Methods, 15, 595–597.

47. Heller, D. and Vingron, M. (2021) SVIM-asm: structural variant detection from haploid and diploid genome assemblies. Bioinformatics, 36, 5519–5521.

48. Jiang, T., Liu, Y., Jiang, Y., Li, J., Gao, Y., Cui, Z., Liu, Y., Liu, B. and Wang, Y. (2020) Long-read-based human genomic structural variation detection with cuteSV. Genome Biol., 21, 189.

49. Li, H. (2018) Minimap2: pairwise alignment for nucleotide sequences. Bioinformatics, 34, 3094–3100.

50. Lin, H.-N. and Hsu, W.-L. (2020) GSAlign: an efficient sequence alignment tool for intra-species genomes. BMC Genomics, 21, 182.

51. Song, B., Marco-Sola, S., Moreto, M., Johnson, L., Buckler, E.S. and Stitzer, M.C. (2022) AnchorWave: Sensitive alignment of genomes with high sequence diversity, extensive structural polymorphism, and whole-genome duplication. Proc. Natl. Acad. Sci. U. S. A., 119.

52. Song, B., Buckler, E.S. and Stitzer, M.C. (2023) New whole-genome alignment tools are needed for tapping into plant diversity. Trends Plant Sci., 10.1016/j.tplants.2023.08.013.

53. Kent, W.J., Baertsch, R., Hinrichs, A., Miller, W. and Haussler, D. (2003) Evolution’s cauldron: duplication, deletion, and rearrangement in the mouse and human genomes. Proc. Natl. Acad. Sci. U. S. A., 100, 11484–11489.

54. Song, J.-M., Xie, W.-Z., Wang, S., Guo, Y.-X., Koo, D.-H., Kudrna, D., Gong, C., Huang, Y., Feng, J.-W., Zhang, W., et al. (2021) Two gap-free reference genomes and a global view of the centromere architecture in rice. Mol. Plant, 14, 1757–1767.

55. Bartenhagen, C. and Dugas, M. (2013) RSVSim: an R/Bioconductor package for the simulation of structural variations. Bioinformatics, 29, 1679–1681.

56. Stein, J.C., Yu, Y., Copetti, D., Zwickl, D.J., Zhang, L., Zhang, C., Chougule, K., Gao, D., Iwata, A., Goicoechea, J.L., et al. (2018) Genomes of 13 domesticated and wild rice relatives highlight genetic conservation, turnover and innovation across the genus Oryza. Nat. Genet., 50, 285–296.

57. Alonge, M., Wang, X., Benoit, M., Soyk, S., Pereira, L., Zhang, L., Suresh, H., Ramakrishnan, S., Maumus, F., Ciren, D., et al. (2020) Major Impacts of Widespread Structural Variation on Gene Expression and Crop Improvement in Tomato. Cell, 182, 145–161.e23.

58. Flint-Garcia, S.A., Thuillet, A.-C., Yu, J., Pressoir, G., Romero, S.M., Mitchell, S.E., Doebley, J., Kresovich, S., Goodman, M.M. and Buckler, E.S. (2005) Maize association population: a high-resolution platform for quantitative trait locus dissection. Plant J., 44, 1054–1064.

59. Ou, S., Collins, T., Qiu, Y., Seetharam, A.S., Menard, C.C., Manchanda, N., Gent, J.I., Schatz, M.C., Anderson, S.N., Hufford, M.B., et al. (2022) Differences in activity and stability drive transposable element variation in tropical and temperate maize. bioRxiv, 10.1101/2022.10.09.511471.

60. Hufford, M.B., Seetharam, A.S., Woodhouse, M.R., Chougule, K.M., Ou, S., Liu, J., Ricci, W.A., Guo, T., Olson, A., Qiu, Y., et al. (2021) De novo assembly, annotation, and comparative analysis of 26 diverse maize genomes. Science, 373, 655–662.

61. Grzybowski, M.W., Mural, R.V., Xu, G., Turkus, J., Yang, J. and Schnable, J.C. (2023) A common resequencing-based genetic marker data set for global maize diversity. Plant J., 113, 1109–1121.

62. Altemose, N., Logsdon, G.A., Bzikadze, A.V., Sidhwani, P., Langley, S.A., Caldas, G.V., Hoyt, S.J., Uralsky, L., Ryabov, F.D., Shew, C.J., et al. (2022) Complete genomic and epigenetic maps of human centromeres. Science, 376, eabl4178.

63. Li, N., He, Q., Wang, J., Wang, B., Zhao, J., Huang, S., Yang, T., Tang, Y., Yang, S., Aisimutuola, P., et al. (2023) Super-pangenome analyses highlight genomic diversity and structural variation across wild and cultivated tomato species. Nat Genet, 55, 852–860.

64. Murat, F., Van de Peer, Y. and Salse, J. (2012) Decoding plant and animal genome plasticity from differential paleo-evolutionary patterns and processes. Genome Biol. Evol., 4, 917–928.

65. Reneker, J., Lyons, E., Conant, G.C., Pires, J.C., Freeling, M., Shyu, C.-R. and Korkin, D. (2012) Long identical multispecies elements in plant and animal genomes. Proc. Natl. Acad. Sci. U. S. A., 109, E1183–91.

66. Li, H., Feng, X. and Chu, C. (2020) The design and construction of reference pangenome graphs with minigraph. Genome Biol., 21, 265.

67. Garrison, E., Sirén, J., Novak, A.M., Hickey, G., Eizenga, J.M., Dawson, E.T., Jones, W., Garg, S., Markello, C., Lin, M.F., et al. (2018) Variation graph toolkit improves read mapping by representing genetic variation in the reference. Nat. Biotechnol., 36, 875–879.

68. Tian, D., Wang, Q., Zhang, P., Araki, H., Yang, S., Kreitman, M., Nagylaki, T., Hudson, R., Bergelson, J. and Chen, J.-Q. (2008) Single-nucleotide mutation rate increases close to insertions/deletions in eukaryotes. Nature, 455, 105–108.

69. Hollister, J.D., Ross-Ibarra, J. and Gaut, B.S. (2010) Indel-associated mutation rate varies with mating system in flowering plants. Mol Biol Evol, 27, 409–416.

70. Sarkar, A., Yang, Y. and Vihinen, M. (2020) Variation benchmark datasets: update, criteria, quality and applications. Database, 2020, baz117.

71. Sigman, M.J. and Slotkin, R.K. (2016) The First Rule of Plant Transposable Element Silencing: Location, Location, Location. Plant Cell, 28, 304–313.

72. Chen, S., Krusche, P., Dolzhenko, E., Sherman, R.M., Petrovski, R., Schlesinger, F., Kirsche, M., Bentley, D.R., Schatz, M.C., Sedlazeck, F.J., et al. (2019) Paragraph: a graph-based structural variant genotyper for short-read sequence data. Genome Biol., 20, 291.

73. Garrison, E., Guarracino, A., Heumos, S., Villani, F., Bao, Z., Tattini, L., Hagmann, J., Vorbrugg, S., Marco-Sola, S., Kubica, C., et al. (2023) Building pangenome graphs. bioRxiv, 10.1101/2023.04.05.535718.

74. Armstrong, J., Hickey, G., Diekhans, M., Fiddes, I.T., Novak, A.M., Deran, A., Fang, Q., Xie, D., Feng, S., Stiller, J., et al. (2020) Progressive Cactus is a multiple-genome aligner for the thousand-genome era. Nature, 587, 246–251.

75. Hickey, G., Monlong, J., Ebler, J., Novak, A.M., Eizenga, J.M., Gao, Y., Human Pangenome Reference Consortium, Marschall, T., Li, H. and Paten, B. (2024) Pangenome graph construction from genome alignments with Minigraph-Cactus. Nat. Biotechnol., 42, 663–673.

76. Marçais, G., Delcher, A.L., Phillippy, A.M., Coston, R., Salzberg, S.L. and Zimin, A. (2018) MUMmer4: A fast and versatile genome alignment system. PLoS Comput. Biol., 14, e1005944.

77. Kiełbasa, S.M., Wan, R., Sato, K., Horton, P. and Frith, M.C. (2011) Adaptive seeds tame genomic sequence comparison. Genome Res., 21, 487–493.

78. Frith, M.C. and Kawaguchi, R. (2015) Split-alignment of genomes finds orthologies more accurately. Genome Biol., 16, 106.

79. Delcher, A.L., Phillippy, A., Carlton, J. and Salzberg, S.L. (2002) Fast algorithms for large-scale genome alignment and comparison. Nucleic Acids Res., 30, 2478–2483.

80. Blanchette, M., Kent, W.J., Riemer, C., Elnitski, L., Smit, A.F.A., Roskin, K.M., Baertsch, R., Rosenbloom, K., Clawson, H., Green, E.D., et al. (2004) Aligning multiple genomic sequences with the threaded blockset aligner. Genome Res., 14, 708–715.

81. Myers, E.W. and Miller, W. (1987) Optimal Alignments in Linear Space.

82. Xiao, C.-L., Chen, Y., Xie, S.-Q., Chen, K.-N., Wang, Y., Han, Y., Luo, F. and Xie, Z. (2017) MECAT: fast mapping, error correction, and de novo assembly for single-molecule sequencing reads. Nat. Methods, 14, 1072–1074.

83. Chaisson, M.J. and Tesler, G. (2012) Mapping single molecule sequencing reads using basic local alignment with successive refinement (BLASR): application and theory. BMC Bioinformatics, 13, 238.

84. Sedlazeck, F.J., Rescheneder, P., Smolka, M., Fang, H., Nattestad, M., von Haeseler, A. and Schatz, M.C. (2018) Accurate detection of complex structural variations using single-molecule sequencing. Nat. Methods, 15, 461–468.

85. Sović, I., Šikić, M., Wilm, A., Fenlon, S.N., Chen, S. and Nagarajan, N. (2016) Fast and sensitive mapping of nanopore sequencing reads with GraphMap. Nat. Commun., 7, 11307.

86. Jain, C., Rhie, A., Hansen, N.F., Koren, S. and Phillippy, A.M. (2022) Long-read mapping to repetitive reference sequences using Winnowmap2. Nat. Methods, 19, 705–710.

87. Nurk, S., Koren, S., Rhie, A., Rautiainen, M., Bzikadze, A.V., Mikheenko, A., Vollger, M.R., Altemose, N., Uralsky, L., Gershman, A., et al. (2022) The complete sequence of a human genome. Science, 376, 44–53.

88. Lander, E.S., Linton, L.M., Birren, B., Nusbaum, C., Zody, M.C., Baldwin, J., Devon, K., Dewar, K., Doyle, M., FitzHugh, W., et al. (2001) Initial sequencing and analysis of the human genome. Nature, 409, 860–921.

89. Hoskins, R.A., Carlson, J.W., Wan, K.H., Park, S., Mendez, I., Galle, S.E., Booth, B.W., Pfeiffer, B.D., George, R.A., Svirskas, R., et al. (2015) The Release 6 reference sequence of the Drosophila melanogaster genome. Genome Res., 25, 445–458.

90. Schnable, P.S., Ware, D., Fulton, R.S., Stein, J.C., Wei, F., Pasternak, S., Liang, C., Zhang, J., Fulton, L., Graves, T.A., et al. (2009) The B73 maize genome: complexity, diversity, and dynamics. Science, 326, 1112–1115.

91. Hosmani, P.S., Flores-Gonzalez, M., van de Geest, H., Maumus, F., Bakker, L.V., Schijlen, E., van Haarst, J., Cordewener, J., Sanchez-Perez, G., Peters, S., et al. (2019) An improved de novo assembly and annotation of the tomato reference genome using single-molecule sequencing, Hi-C proximity ligation and optical maps. bioRxiv, 10.1101/767764.

92. Liao, Y., Wang, J., Zhu, Z., Liu, Y., Chen, J., Zhou, Y., Liu, F., Lei, J., Gaut, B.S., Cao, B., et al. (2022) The 3D architecture of the pepper genome and its relationship to function and evolution. Nat. Commun., 13, 3479.

93. Qin, C., Yu, C., Shen, Y., Fang, X., Chen, L., Min, J., Cheng, J., Zhao, S., Xu, M., Luo, Y., et al. (2014) Whole-genome sequencing of cultivated and wild peppers provides insights into Capsicum domestication and specialization. Proc. Natl. Acad. Sci. U. S. A., 111, 5135–5140.

94. Shang, L., He, W., Wang, T., Yang, Y., Xu, Q., Zhao, X., Yang, L., Zhang, H., Li, X., Lv, Y., et al. (2023) A complete assembly of the rice Nipponbare reference genome. Mol Plant, 16, 1232–1236.

95. Kawahara, Y., de la Bastide, M., Hamilton, J.P., Kanamori, H., McCombie, W.R., Ouyang, S., Schwartz, D.C., Tanaka, T., Wu, J., Zhou, S., et al. (2013) Improvement of the Oryza sativa Nipponbare reference genome using next generation sequence and optical map data. Rice, 6, 4.

96. Zhou, Y., Yu, Z., Chebotarov, D., Chougule, K., Lu, Z., Rivera, L.F., Kathiresan, N., Al-Bader, N., Mohammed, N., Alsantely, A., et al. (2023) Pan-genome inversion index reveals evolutionary insights into the subpopulation structure of Asian rice. Nat. Commun., 14, 1567.

97. Chen, C., Wu, Y., Li, J., Wang, X., Zeng, Z., Xu, J., Liu, Y., Feng, J., Chen, H., He, Y., et al. (2023) TBtools-II: A ‘one for all, all for one’ bioinformatics platform for biological big-data mining. Mol Plant, 16, 1733–1742.

98. Liao, Y., Wang, J., Zhu, Z., Liu, Y., Chen, J., Zhou, Y., Liu, F., Lei, J., Gaut, B.S., Cao, B., et al. (2021) The 3D architecture of the pepper (Capsicum annum) genome and its relationship to function and evolution. bioRxiv, 10.1101/2021.12.10.470457.

99. Zhou, Y., Chebotarov, D., Kudrna, D., Llaca, V., Lee, S., Rajasekar, S., Mohammed, N., Al-Bader, N., Sobel-Sorenson, C., Parakkal, P., et al. (2020) A platinum standard pan-genome resource that represents the population structure of Asian rice. Sci Data, 7, 113.

100. Harris, R.S. (2007) Improved pairwise alignment of genomic DNA. Ph.D. Thesis, The Pennsylvania State University.

101. Li H. (2013) Aligning sequence reads, clone sequences and assembly contigs with BWA-MEM. arXiv:1303.3997v2 [q-bio.GN].

