## Supplementary Material for "Benchmarking, detection, and genotyping of structural variants in a population of whole-genome assemblies using the SVGAP pipeline"

#### Contents

1. Supplementary Notes
2. Supplementary Methods
3. Supplementary Tables
4. Supplementary Figures
5. References

### Supplementary Notes

#### Supplementary Note 1: A practical guide to using SVGAP

##### Section 1: Preparing the genome assemblies

###### Assembly quality requirement

The SVGAP pipeline was originally designed for genome assemblies of chromosome-level quality; however, it can also be effectively applied to contig-level assemblies. Theoretically, there are no specific quality requirements for the genome assembly, such as N50 or contig numbers. Nevertheless, we highly recommend that users include only chromosomes and long contigs when preparing the input genome assemblies. Currently, SVGAP is optimized for haploid genomes. If you have phased genomes with multiple haplotypes, the recommended approach is to treat each haplotype as an individual assembly.

###### Naming

To effectively utilize the SVGAP pipeline, users must adhere to specific naming guidelines for genome assemblies and their chromosomes/contigs. Here are some key points to keep in mind:

1) Assembly names do not need to include file extensions such as ".fasta" or ".fa".

2) Avoid naming assemblies solely using numbers. For example, naming an assembly "9311" could potentially cause issues in certain situations. Instead, consider using "N9311".

3) Do not name the query assembly in a way that completely matches or overlaps with the names of reference assemblies. For instance, if the reference name is "MH63", it would be advisable to avoid naming any query assembly "H63" or "MH6".

4) Chromosome names should follow the format "GenomeID.ChromosomeID", such as "MH63.chr01".

###### Splitting (Optional)

If you have an extreme size of plant genomes, such as maize and pepper, you may need to split the genome into small pieces, and then conduct the pairwise genome alignment. Here we provide a script called ***0_SplitFa.pl*** to facilitate this splitting work.

You just need to split the query one in the genome assemblies folder, here is an example:

```
$ perl 0_SplitFa.pl ZH97 2000000 400000
```

This script will split a FASTA file into fixed-sized windows with a specified step size. The first parameter "ZH97" refers to the input FASTA file, the second parameter "2000000" indicates the size of the fixed-sized windows, and the third parameter "400000" denotes the step size for moving along the sequence.

##### Section 2: a step-by-step tutorial

In this section, we used 49 rice genomes to demonstrate how to use SVGAP step by step:

###### Constructing a working folder

To start the project, we first constructed a folder named rice:

```
$ mkdir rice
$ mkdir rice/genome
```

###### Moving working genomes to subfolder

Once the working genomes have been named and filtered for short contigs, they are moved to the rice/genome subfolder. Please be note that even if you split the query genomes, the rice/genome should only contain unsplit files.

###### Step 0: Whole genome alignment

Currently, SVGAP supports alignment results from a variety of aligners, including Lastz, Last, MUMmer, minimap2, GSAlign, and AnchorWave:

1) For small and medium-sized genomes (<500 Mb), we recommend utilizing Last and MUMmer.

2) For larger genomes, particularly in the case of plants, it is advisable to use minimap2 or AnchorWave instead.

3) If you opt for AnchorWave, please ensure that a dependable gene annotation GFF3 file is accessible for the reference genome and that there are no considerable rearrangements between the reference and query genomes.

In the case of our rice data, we opted to employ MUMmer4 (with default parameters) to conduct alignments between the reference genome (MH63) and each of the 48 query genomes.

Here's an example:

```
$ nucmer -t 4 --prefix=MH63vsBasm ./genome/MH63 ./genome/Basm
```

If you would like to use other alternative aligners, you may refer the following commands:

#### minimap2

```
$ minimap2 -c --cs=long ./genome/MH63 ./genome/Basm > MH63vsBasm.chr.paf
```

#### last

```
$ lastdb -P4 -uNEAR MH63dat ./genome/MH63
$ last-train -P4 --revsym -E0.05 -C2 MH63dat ./genome/Basm > MH63vsBasm.train
$ lastal -p MH63vsBasm.train MH63dat Basm > MH63vsBasm.maf
```

**Step 1: Converting the alignment output from a particular aligner to the AXT format alignment files**

SVGAP incorporates a series of UCSC programs (e.g., axtChain, ChainNet, and netSyntenic developed by Jim Kent) to preprocess alignment files before identifying genomic variants. Hence, it is necessary to convert alignment files from other aligners with different formats, such as *.delta* from MUMmer4, *.paf* from minimap2, and *.maf* from LAST/LASTZ/AnchorWave, to the AXT alignment format (the *.axt* format is described in http://genome.ucsc.edu/goldenPath/help/axt.html) prior to processing them within the pipeline. To facilitate this conversion, SVGAP provides a script called ***1_Convert2Axt.pl*** to handle this task. To find the detailed usage information about the *1_Convert2Axt.pl* script, you can run:

```
$ perl 1_Convert2Axt.pl --help
```

In our case of rice data, we saved all *delta* alignment files produced by MUMmer4 to a single output folder rice/alignment and created a working folder rice/chainnet/Target_Ref.

**Rearrange output files**

```
$ mkdir  ./rice/alignment
$ mkdir ./rice/chainnet
$ mv *.delta ./rice/alignment
```

**Run 1_Convert2Axt.pl**

```
$ perl 1_Convert2Axt.pl -ali mummer -input ./rice/alignment -wk ./rice/chainnet
```

Please note that if you split query genomes, you should provide a genome size file (tab-delimited form) named “genomeID.sizes” in the working folder where the *.axt* file will be saved before running *1_Convert2Axt.pl*.

Here is an example of the genome size file:

```
MH63.chr01	23513712
MH63.chr02	32079331
```

After running the above commands, the resulting .axt files have been saved in the rice/chainnet folder and ready for further processing.

**Step 2: Constructing chains and nets from pairwise alignments**

To accurately identify genetic variants between the reference and query genomes, the raw alignments (.axt file) must be processed for synteny, and paralogous regions need to be removed. This ensures that only the orthologous regions are retained for analysis. This task can be accomplished by employing a set of UCSC programs (find necessary information from http://genomewiki.ucsc.edu/index.php/Whole_genome_alignmen_howto) that are sequentially invoked in the script 2_ChainNetSyn.pl within the SVGAP pipeline which will generate a filtered net file for syntenic alignments only and a single-coverage (on the reference genome) collection of pairwise alignments.

To find more detailed usage about 2_ChainNetSyn.pl script, run:

```
$ perl 2_ChainNetSyn.pl --help
```

In our case we run the command as following:

```
$ perl 2_ChainNetSyn.pl --gd ../genome/ --ad chainnet/Target_Ref/ --lst ../genome/g.lst --sing
```

The filtered *.net* files for syntenic alignments will be saved in ./rice/synnet/ and the sing-coverage alignment *.maf* files will be saved in ./rice/singRaw/.

In most cases, especially when comparing two highly divergent genomes, it is recommended to utilize the additional script ***3_SynNetFilter.pl*** to filter the syntenic alignment *.net* files.This script enhances precision while slightly reducing sensitivity, resulting in improved accuracy for such comparisons.

To see how to use the 3_SynNetFilter.pl script, run:

```
$ perl 3_SynNetFilter.pl --help
```

We run the following command for the rice test data:

```
$ perl 3_SynNetFilter.pl --synnet synnet/ --gd ../genome/ --chain chainnet/Target_Ref
```

The final filtered *.net* files for syntenic alignments are saved in ./rice/CleanSynNet/ which will be used for SV calling and the cleaned single-coverage alignment *.maf* files are saved in ./rice/filtered_sing/ which will be used for SNP calling and SV genotyping.

**Step 3: Identification of structural variants (SVs)**

The SVGAP pipeline infers SVs directly from the final filtered *net* files (the *net* format is described in http://genome.ucsc.edu/goldenPath/help/net.html). Currently, five common SV types between pairwise genome comparisons can be routinely identified using the *script* ***4_PairwiseSV.pl*** within the SVGAP pipeline, including insertions, deletions, tandem duplications, inversions, and translations. Additionally, local highly divergent regions (i.e. with gaps on both target and query sequences) are defined as a complex type.

To see how to use the *4_PairwiseSV.pl* script, you can run:

```
$ perl 4_PairwiseSV.pl --help
```

Below is the command we run for the rice data:

```
$ perl 4_PairwiseSV.pl --syndir ./rice/CleanSynNet --gd ./rice/genome/ --outdir ./rice/CleanSV
```

The detected SV calls are stored in the directory ./rice/CleanSV/. For each pairwise comparison, two sets of SVs are reported, with coordinates that correspond to both the reference and query genomes.

**Step 4: Merge and remove redundant SV calls**

Since SVs are identified between the reference genome and each of the query genomes, it is essential to merge the SVs from all comparisons. This merging process aims to generate a non-redundant dataset of SV calls that encompasses all the comparisons. By doing so, each SV can be further genotyped across all individuals involved in the study, enabling construction of the pan-genome. The script ***5_Combined.pl*** within the SVGAP pipeline can work for this task.

To see how to use the *5_Combined.pl* script, you can run:

```
$ perl 5_Combined.pl --help
```

We run the following command to generate the non-redundant SV calls for all 48 pairwise companions.

```
$ perl 5_Combined.pl --sv ./rice/CleanSV --refname MH63
```

Executing the above command will generate a folder called ./rice/CombinedSV/. Within this folder, five types of non-redundant SVs will be generated. It is worth mentioning that the *insertions* and *deletions* have been categorized into two classes according to their lengths; specifically, events that are equal to or longer than 50 bp (>50 bp) are in one class, while the rest are in another.

**Step 5: Re-genotyping the merged non-redundant SV calls across all samples based on pairwise (single coverage) alignments**

Although the above step has provided preliminary genotyping information, the information still lacks completeness. For instance, the detection of SVs was limited to specific individuals, leaving uncertainty as to whether the absence of these SVs in other individuals is genuine or merely a result of inaccurate sequencing and assembling, or sequence loss in those regions. Therefore, additional methods are necessary to achieve complete genotyping information for all SVs across all samples. The SVGAP utilizes pairwise single coverage alignment files to enhance the genotyping of each SV by extracting and comparing the local alignment in order to determine the SV status in each sample. The scripts ***6_DELGenotyping.pl*** and ***7_INSGenotyping.pl*** within the SVGAP pipeline are utilized to perform genotyping for deletions and insertions (with a length greater than or equal to 50 base pairs), respectively.

**Genotyping Deletions (≥50 bp)**

To see how to use the 6_DELGenotyping.pl script, you can run:

```
$ perl 6_DELGenoptyping.pl --help
```

We run the tested rice data with the following command:

```
$ perl 6_DELGenoptyping.pl --del ./rice/CombinedSV/All.DELs.50bplarge.bed.combined.sorted.txt \
--singmaf ./rice/filtered_sing/ --refseq ./rice/genome/MH63 --refsize ./rice/genome/MH63.sizes \
--refname MH63 -t 12
```

This step will result in a *vcf* file named All.DELs.50bplarge.bed.combined.sorted.txt.vcf in the folder where the All.DELs.50bplarge.bed.combined.sorted.txt file is kept.

**Genotyping Insertions (≥50 bp)**

To see how to use the *7_INSGenotyping.pl* script, you can run:

```
$ perl 7_INSGenotyping.pl --help
```

We run the tested rice data with the following command:

```
$ perl 7_INSGenoptyping.pl --ins ./rice/CombinedSV/All.INTs.50bplarge.bed.combined.sorted.txt \
--query ../genome/ -sing filtered_sing/ --stretcher ../../SV-GAPS/pub/Stretcher/ --refseq ../genome/MH63 \
--refsize ../genome/MH63.sizes --refname MH63
```

This step will result in a *vcf* file named All.INTs.50bplarge.bed.combined.sorted.txt.vcf in the folder where the All.INTs.50bplarge.bed.combined.sorted.txt file is kept.

**Genotyping InDels**

Small deletions and insertions (<50 bp) are separately genotyped with *8_InDelGenotyping.pl*.

To see how to use the *8_InDelGenotyping.pl* script, you can run:

```
$ perl 8_InDelGenotyping.pl --help
```

Genotyping small deletions in our case:

```
$ perl 8_InDelGenoptyping.pl --indel ./rice/CombinedSV/All.DELs.50bpsmall.bed.combined.sorted.txt \
--indeltype del --refseq ./rice/genome/MH63 --refsize ./rice/genome/MH63.sizes --refname MH63 \
--query ./rice/genome/ --singmaf ./rice/filtered_sing/ --outdir MH63_del --t 12
```

This step will result in a *vcf* file named All.DELs.50bpsmall.bed.combined.sorted.txt.vcf in the folder where the All.DELs.50bpsmall.bed.combined.sorted.txt file is kept.

Genotyping small insertions in our case:

```
$ perl 8_InDelGenoptyping.pl --indel ./rice/CombinedSV/All.INTs.50bpsmall.bed.combined.sorted.txt \
--indeltype ins --refseq  ./rice/genome/MH63 --refsize ./rice/genome/MH63.sizes --refname MH63 \
--query ./rice/genome/ --singmaf ./rice/filtered_sing/ --outdir MH63_ins --t 12
```

This step will result in a *vcf* file named All.INTs.50bpsmall.bed.combined.sorted.txt.vcf in the folder where the All.INTs.50bpsmall.bed.combined.sorted.txt file is kept.

**Genotyping SNPs**

SVGAP also offers a script called 9_SNPgenotyping.pl, which allows for the direct calling of SNPs from the single-coverage *.maf* files. It generates a *.vcf* file that includes all samples and their respective SNP information.

To see how to use the *9_SNPgenotyping.pl* script, you can run:

```
$ perl 9_SNPgenotyping.pl --help
```

The command we used for the test data is:

```
$ perl 9_SNPgenotyping.pl --refseq ./rice/genome/MH63 --refsize ./rice/genome/MH63.sizes \
--refname MH63 --singmaf ../rice/filtered_sing/ --outdir SNPVCF --t 8
```

The SNP genotyped file All.SNP.vcf will report in the SNPVCF folder.

**Step 6: Annotating structural variants**

When we acquire precise catalogs of structural variants, it is crucial to elucidate the potential mechanisms driving the formation of SVs and their functional impacts. We offer a script called ***10_SVannotation.pl*** to infer the primary common factors in plant genomes, such as tandem duplication, inverted tandem duplication, transposon elements (TEs) insertion, nest TEs insertion, and complete/partial gene duplication, that may be involved in SV formation.

To see how to use the *10_SVannotation.pl* script, you can run:

```
$ perl 10_SVannotation.pl --help
```

The command we used for the test data is:

```
$ perl 10_SVannotation.pl --ref ./rice/genome/MH63 --vcf All.INTs.50bplarge.bed.combined.sorted.txt.vcf \
--svtype INS --te TE.EDTA.lab.fa --cds MSU7.cds.fa --out INS.annotated.vcf
$ perl 10_SVannotation.pl --ref ./rice/genome/MH63 --vcf All.DELs.50bplarge.bed.combined.sorted.txt.vcf \
--svtype DEL --te TE.EDTA.lab.fa --cds MSU7.cds.fa --out INS.annotated.vcf
```

The annotated *.vcf* file(s) will report in the current folder.

### Supplementary Note 2: Evaluation of execution time and memory usage for different genome aligners

##### Installation of genome aligners

We selected 14 genome aligners to evaluate their execution time and memory usage for plant genome alignment, including minimap2, AnchorWave, MUMmer4, Blasr, BWA-MEN, GraphMap, Last, Lastz, MECAT2, Ngmlr, Pbmm2, GSAlign, Unimap and winnowmap. We installed most of them through ***conda*** with one exception.

#### Thirteen of them can be installed with conda

```
$ conda install -c bioconda minimap2
$ conda install -c bioconda -c conda-forge anchorwave
$ conda install bioconda::mummer4
$ conda install bioconda::blasr
$ conda install -c bioconda bwa
$ conda install bioconda::graphmap
$ conda install bioconda::last
$ conda install bioconda::lastz
$ conda install ngmlr -c bioconda
$ conda install -c bioconda pbmm2
$ conda install bioconda::gsalign
$ conda install bioconda::unimap
$ conda install bioconda::winnowmap

MECAT2 was installed from GitHub :
$ git clone https://github.com/xiaochuanle/MECAT2.git
$ cd MECAT2
$ make
```

##### Testing for genome alignment

The command we used for the test data is:

1. **minimap2** ***(version 2.22-r1101)***

*## (2min 42s, 7.64 GB)*

```
$ time minimap2 -x asm5 ~/MH63RS3.fasta ~/ZS97RS3.fasta > map.paf
```

2. ***AnchorWave (v1.2.3)***

*## Extract CDS (0.20 min, 959.61 Mb)*

```
$ time anchorwave gff2seq -i MH63RS3.gff3 -r MH63RS3.fasta -o cds.fa
```

*## Align CDS to the reference genome (3.20 min, 12.85 Gb)*

```
$ time minimap2 -x splice -t 10 -k 12 -a -p 0.4 -N 20 MH63RS3.fasta cds.fa > ref.sam
```

*## Align CDS to the query genome (3.53 min, 13.47 Gb)*

```
$ time minimap2 -x splice -t 10 -k 12 -a -p 0.4 -N 20 S97RS3.fasta cds.fa > query.sam
```

*## Perform genome alignment (78.52 min, 17.13 Gb)*

```
$ time anchorwave genoAli -i MH63RS3.gff3 -as cds.fa -r MH63RS3.fasta -a query.sam -ar ref.sam \
-s ZS97RS3.fasta -n SZ.anchors -o anchorwave.maf -f anchorwave.f.maf > SZ.log
```

##### *3. MUMmer4 (4.0.0rc1)*

The command we used for the test data is:

*## Alignment (103.85 min,4.65 Gb)*

```
$ time nucmer -p out MH63RS3.fasta ZS97RS3.fasta
```

##### *4. Blasr (5.3.5-5.3.5)*

*Blasr* is used to align PacBio reads to the reference genome, but it keeps throwing errors during usage, and the cause is currently unclear.

The command we used for the test data is:

```
$ time blasr ZS97RS3.fasta MH63RS3.fasta --out test.out -m 5 --header --nproc 50
```

We tried modifying the parameters, but the errors persist:

```
$ time blasr ZS97RS3.fasta MH63RS3.fasta --out test.out --maxMatch 10000000000 --maxLCPLength 10000000000
```

##### *5. BWA-MEM (0.7.17-r1188)*

The command we used for the test data is:

*## Construct genome index (4.22 min, 569.72 Gb)*

```
$ time bwa index MH63RS3.fasta
```

*## Alignment (time-consuming, 10.69 Gb)*

```
$ time bwa mem MH63RS3.fasta ZS97RS3.fasta > output.sam
```

##### *6. GraphMap (v0.6.3)*

*GraphMap* is used to align Nanopore reads to the reference genome. The command we used for the test data is:

```
$ time graphmap2 align -r MH63RS3.fasta -d ZS97RS3.fasta -o graphmap_out.sam
```

Unfortunately, the result cannot be output due to our limited memory.

##### 7. *Last (1584)*

The command we used for the test data is:

*### Simple version*

*## Construct reference genome database (2.07 min, 1.89 Gb)*

```
$ time lastdb MHdb MH63RS3.fasta
```

*## Alignment (97.40 min, 3.44 Gb)*

```
$ time lastal MHdb ZS97RS3.fasta > myalns.maf
```

*## Make a dotplot (64.25 min, 118.68 Gb)*

```
$ time last-dotplot myalns.maf
```

*### Detailed version*

*## Construct reference genome database (0.48 min,1.53 GB)*

```
$ time lastdb -P8 -c -uRY128 MH63db MH63RS3.fasta
```

*## Training (0.07 min, 651 Mb)*

```
$ time last-train -P8 --revsym -C2 MH63db ZS97RS3.fasta > zs.train
```

*## Mang to one alignment (1.17 min,13.99 GB)*

```
$ time lastal -P8 -D1e9 -C2 --split-f=MAF+ -p zs.train MH63db ZS97RS3.fasta > many-to-one.maf
```

#### One to one filtration *(0.40 min, 2.31 GB)*

```
$ time last-split -r many-to-one.maf > one-to-one.maf
```

*## Make a dotplot (0.20 min, 98.39 Mb)*

```
$ time last-dotplot one-to-one.maf
```

##### 8. *Lastz (1.04.22)*

*LastZ* only supports pairwise alignment of one chromosome to another, so we used a looping method for whole genome alignment:

*## Extract each chromosome to a new FASTA file*

```
$ awk '/^>/ {if (f) close(f); f = "MH63" substr($0, 2,5) ".fa"; print > f} !/^>/ {print > f}' MH63RS3.fasta
$ awk '/^>/ {if (f) close(f); f = "ZS" substr($0, 2,5) ".fa"; print > f} !/^>/ {print > f}' ZS97RS3.fasta
```

*## "Looping" alignment (8.47 min, 162.79 Mb)*

```
$ for file in ZS*; do name="${file:2:5}"; lastz "MH63${name}.fa" "${file}" --notransition --step=20 --nogapped \
--format=maf > "${name}.maf"; done
```

##### 9. *MECAT2 (2.0.0)*

*MECAT2* is used to align SMRT reads to the reference genome. We used 80 threads because the execution was too slow:

*## Alignment (45.47 min, 21.28 Gb)*

```
$ time mecat2map -num_threads 80 ZS97RS3.fasta MH63RS3.fasta > result.m4
```

##### 10. *Ngmlr (0.2.7)*

*Ngmlr* is also used to align SMRT reads to the reference genome.

The command we used for the test data is:

*## Alignment (time-consuming)*

```
$ time ngmlr -r ~/MH63RS3.fasta -q ~/ZS97RS3.fasta -o test.sam
```

##### 11. *Pbmm2 (1.14.99)*

*Pbmm2* is also used to align SMRT reads to the reference genome. We also used 80 threads because the execution was too slow:

*## Construct genome index (0.10 min, 2.39 Gb)*

```
$ time pbmm2 index MH63RS3.fasta ref.mmi
```

*## Alignment (57.88 min,438.5 Gb)*

```
$ time pbmm2 align ref.mmi ZS97RS3.fasta ref.movie.bam -j 80
```

##### 12. *GSAlign (v1.0.22)*

The command we used for the test data is:

*## Alignment (5.45 min, 2.46 Gb)*

```
$ time GSAlign -r MH63RS3.fasta -q ZS97RS3.fasta -o output
```

***13. Unimap (0.1-r41)***

The command we used for the test data is:

*### Index-free genome alignment (2.28 min,9.28 Gb)*

```
$ time unimap -b24 MH63RS3.fasta ZS97RS3.fasta > out.paf
```

*### Index-depended genome alignment*

*## Construct genome index (0.67 min, 7.31 Gb)*

```
$ time unimap -b24 -d out.umi ~/MH63RS3.fasta
```

*## Alignment (2.23 min,6.21 Gb)*

```
$ time unimap -c out.umi ~/ZS97RS3.fasta > out1.paf
```

***14. winnowmap (2.03)***

The command we used for the test data is:

*## Meryl count（1.82 min,2.21 Gb）*

```
$ time meryl count k=19 output merylDB MH63RS3.fasta
```

*## Meryl print (0.12 min, 1.08 Gb)*

```
$ time meryl print greater-than distinct=0.9998 merylDB > repetitive_k19.txt
```

*## Alignment (6.45 min, 6.44 Gb)*

```
$ time winnowmap -W repetitive_k19.txt -x asm20 MH63RS3.fasta ZS97RS3.fasta > output.paf
```

### Supplementary Methods

#### Approaches for benchmarking analysis using a phylogeny-based simulated dataset

To evaluate the extent to which sequence divergence impacts SV detection, we propose the following strategy for benchmark analysis. ***First***, we selected eight divergent genomes, using one as the reference, from both the *Oryza* and *Solanum* phylogenies. The genomes exhibit varying levels of sequence divergence relative to the reference. ***Second***, for each phylogeny-based genome set, we simulated SVs, including insertions and deletions, in all eight genomes using RSVSim [1]. ***Third***, we identified SVs between the reference and the eight simulated genomes, as well as between the reference and the eight original genomes. ***Fourth***, the latter SV set (background calls) was subtracted from the former to obtain the set of induced SVs. ***Finally***, these remaining SV calls were compared against the simulated SV set to measure the recall, precision, and *F1*-score for each genome.

##### Step 1: Simulating SVs in the genomes

We used the following script, run_RSVSim.Rscript, to simulate SVs for each genome.

```
#!/usr/bin/env Rscript
library(argparser, quietly = TRUE)
## Create a parser
p <- arg_parser("RSVSim runner")
## Add command line arguments
p <- add_argument(p, "--genome", help = "genome_path", type = "character")
p <- add_argument(p, "--output", help = "output_path", type = "character")
p <- add_argument(p, "--delnum", help = "delnum", type = "character")
p <- add_argument(p, "--insnum", help = "insnum", type = "character")
## Parse the command line arguments
argv <- parse_args(p)
genome_file <- argv$genome
output_file <- argv$output
delnum <- as.numeric(argv$delnum)
insnum <- as.numeric(argv$insnum)
library("RSVSim")
## Setting SV size for DELs and INTs
delSizes = estimateSVSizes(n = delnum, minSize = 50, maxSize = 20000, default = "deletions", hist = TRUE)
intSizes = estimateSVSizes(n = insnum, minSize = 50, maxSize = 20000, default = "insertions", hist = TRUE)
## With additional breakpoint mutations
simulateSV(output = output_file, genome = genome_file, sizeDels = delSizes, dels = delnum, sizeIns = intSizes, ins = insnum,
percCopiedIns = 1)
```

For example, we used the following command to introduce 10,000 deletions and 10,00 insertions into the CJ14 genome.

```
$ Rscript run_RSVSim.Rscripts --genome CJ14 --output CJ14_Simulated –delnum 10000 --insnum 10000
```

The output are files recording the variant's coordinates: **deletions.csv** and **insertions.csv**, and the simulated genome: **genome_rearranged.fasta**, all have been saved in the folder CJ14_Simulated.

Since the coordinates in deletions.csv and insertions.csv are based on the original genomes, they need to be lifted over to the simulated genomes for comparisons between the reference and the simulated genomes. The following commands and the script SimulationReformatSV.pl were used for this purpose.

```
$ cat deletions.csv | grep -v Name | cut -f1-5 > simulatedA.DELs.bed
$ cat insertions.csv | grep -v Name | cut -f1,5-8 > simulatedA.INTs.bed
$ cat simulatedA.DELs.bed simulatedA.INTs.bed | sort -k2,2 -k3,3n > simulatedA.All.variant.sort.bed
$ perl SimulationReformatSV.pl simulatedA.All.variant.sort.bed
```

This step will generate .bed files containing SV coordinates based on the simulated genomes.

**Step 2: Whole genome alignment**

The generated simulated data can be used for benchmarking aligners and callers. Here we show how we benchmark callers with SVGAP,  including Last, MUMmer4, minimap2 and AnchorWave.

```
### LAST
## Construct reference genome database
$ lastdb -P8 -uNEAR Adatabase ./genome/A
## Training
$ last-train -P8 -E0.05 --revsym -C2 Adatabase ./genome/B > AvsB.train
## Alignment
$ lastal --split-f=MAF -p AvsB.train Adatabase ./genome/B > AvsB.chr.maf

### MUMmer4
$ nucmer -t 4 --prefix=AvsB ./genome/A .genome/B

### minimap2
$ minimap2 -t 2 -c --cs=long -x asm5 ./genome/A ./genome/B > AvsB.chr.paf

### AnchorWave
## Extract CDS
$ anchorwave gff2seq -i ./genome/A.gff -r ./genome/A -o ./genome/A.CDS.fa
## Align CDS to the query genome
$ minimap2 -x splice -t 10 -k 12 -a -p 0.4 -N 20 ./genome/B ./genome/A.CDS.fa > B.sam
## Align CDS to the reference genome
$ minimap2 -x splice -t 10 -k 12 -a -p 0.4 -N 20 ./genome/A ./genome/A.CDS.fa > A.sam
## Perform genome alignment
$ anchorwave genoAli -i ./genome/A.gff -as ./genome/A.CDS.fa -r ./genome/A -a B.sam -ar A.sam -s ./genome/B -n B.anchors \
-o B.maf -f AvsB.maf > B.log
```

**Step 3: Calling SVs**

Using the alignment results from the four tested aligners, we applied the SVGAP pipeline to call SVs between each genome and the reference. The single-coverage alignment *.maf* files (saved in the singRaw directory) and the detected SV calls (stored in the CleanSV directory) were used for subsequent analysis.

We used the following command to calculate the alignable proportion of each genome relative to the reference.

```
$ grep -v '#' singRaw/A.simulatedB.sing.maf | awk -v term="simulatedB" '$2 ~ term' | \
awk '{sum +=$4} END {print  sum}' > AvssimulatedB.alignedLength

## Calculate the total genome size
$ seqkit faidx genome/simulated
$ awk '{sum +=$2} END{print sum}' simulatedB.fai > simulatedB.totallength

$ echo "$(cat AvssimulatedB.alignedLength)/$(cat simulatedB.totallength)" | bc -l | \
awk -v prefix="mappingrate=" '{print prefix $0}'
```

**Step 4 : Benchmarking**

We calculated recall, precision, and F1 score using the following formula:

```
Precision = TP / (TP + FP)
Recall = TP / (TP + FN)
F1 Score = (2 * Precision * Recall) / (Precision + Recall)
```

, where TP, FP, FN and TN are defined as following:

*# TP (True positive) SVGAP predicted there is a SV and it actually has.*

*# FP (False positive): SVGAP predicted there is a SV but it actually hasn't.*

*# FN (False negative): SV-GAPS predicted there isn't a SV but it actually has.*

*#TN (True negative): SVGAP predicted there isn't a SV and it actually hasn't.*

***Benchmark aligners for DEL (deletion) between genome A and simulated genome A'***

```
## Select DELs ≥50 bp
$ cat CleanSV/A.simulatedA.rawSV/AvssimulatedA.chain.filter.tnet.synnet.refiltered.synnet.out.indel.sort | grep -v "#" | awk
'$8>49 {print $5"\t"$6"\t"$6+$8}' > AvssimulatedA.DELs.bed
## Calculation for TP
$ bedtools intersect -wa -wb -a simulatedA.All.variant.sort.bed.del.bed -b AvssimulatedA.DELs.bed -f 0.8 -F 0.8 | cut -f1-3 |
sort | uniq | wc -l | awk -v prefix="TP=" '{print prefix $0}'
## Calculation for FP
$ bedtools intersect -b simulatedA.All.variant.sort.bed.del.bed -a AvssimulatedA.DELs.bed -f 0.8 -F 0.8 -v | sort | uniq | wc -l |
awk -v prefix="FP=" '{print prefix $0}'
## Calculation for FN
$ bedtools intersect -a simulatedA.All.variant.sort.bed.del.bed -b AvssimulatedA.DELs.bed -f 0.8 -F 0.8 -v | sort | uniq | wc -l |
awk -v prefix="FN=" '{print prefix $0}'
```

***Benchmark aligners for INT (insertion) between genome A and simulated genome A'***

```
## Select INTs ≥50 bp
$ cat CleanSV/simulatedA.A.rawSV/AvssimulatedA.chain.filter.qnet.synnet.refiltered.synn et.out.indel.sort | awk '$8>49' \
> AvssimulatedA.INTs.bed
## Calculation for TP
$ bedtools intersect -wa -wb -a simulatedA.All.variant.sort.bed.ins.bed -b AvssimulatedA.INTs.bed -f 0.8 -F 0.8 | cut -f1-3 | \
```

```
sort | uniq | wc -l | awk -v prefix="TP=" '{print prefix $0}'
## Calculation for FP
$ bedtools intersect -b simulatedA.All.variant.sort.bed.ins.bed -a AvssimulatedA.INTs.bed -f 0.8 -F 0.8 -v | sort | uniq | wc -l | \
awk -v prefix="FP=" '{print prefix $0}'
## Calculation for FN
$ bedtools intersect -a simulatedA.All.variant.sort.bed.ins.bed -b AvssimulatedA.INTs.bed -f 0.8 -F 0.8 -v | sort | uniq | wc -l | \
awk -v prefix="FN=" '{print prefix $0}'
```

***Benchmark aligners for DEL (deletion) between genome A and simulated genome B'***

```
## Select DELs ≥ 50 bp
$ awk '$8>49' CleanSV/A.B.rawSV/AvsB.chain.filter.tnet.synnet.refiltered.synnet.out.indel.sort > AvsB.DELs.bed
$ awk '$8>49' CleanSV/A.simulatedB.rawSV/AvssimulatedB.chain.filter.tnet.synnet.refiltered.synnet.out.indel.sort \
> AvssimulatedB.DELs.bed
## Extract the different DELs set between A vs. B and A vs. simulatedB
$ bedtools intersect -a AvssimulatedB.DELs.bed -b AvsB.DELs.bed -v -f 0.8 -F 0.8 | awk '{print $5"\t"$6"\t"$6+$8}' \
> simulatedBvsB.diff.DELs.bed
## Calculation for TP
$ bedtools intersect -wa -wb -a simulatedB.All.variant.sort.bed.del.bed -b simulatedBvsB.diff.DELs.bed -f 0.8 -F 0.8 | \
cut -f1-3 | sort | uniq | wc -l | awk -v prefix="TP=" '{print prefix $0}'
## Calculation for FP
$ bedtools intersect -b simulatedB.All.variant.sort.bed.del.bed -a simulatedBvsB.diff.DELs.bed -f 0.8 -F 0.8 -v | sort | uniq | \
wc -l | awk -v prefix="FP=" '{print prefix $0}'
## Calculation for FN
$ bedtools intersect -a simulatedB.All.variant.sort.bed.del.bed -b simulatedBvsB.diff.DELs.bed -f 0.8 -F 0.8 -v | sort | uniq | \
wc -l | awk -v prefix="FN=" '{print prefix $0}'
```

***Benchmark aligners for INT (insertion) between genome A and simulated genome B'***

```
## Select INTs ≥50 bp
$ grep -v '#' CleanSV/B.A.rawSV/AvsB.chain.filter.qnet.synnet.refiltered.synnet.indel.sort | \
awk '$8>49 {print $5"\t"$6"\t"$6+$8"\t"$1"\t"$2"\t"$3}' > AvsB.INTs.bed
$ grep -v '#' CleanSV/simulatedB.A.rawSV/AvssimulatedB.chain.filter.qnet.synnet.refiltered.synnet.indel.sort | \
awk '$8>49 {print $5"\t"$6"\t"$6+$8"\t"$1"\t"$2"\t"$3}' > AvssimulatedB.INTs.bed
## Extract the different INTs set between A vs. B and A vs. simulatedB
$ bedtools intersect -a AvssimulatedB.INTs.bed -b AvsB.INTs.bed -v -f 0.8 -F 0.8 | cut -f4-6 > simulatedBvsB.diff.INTs.bed
## Calculation for TP
$ bedtools intersect -a simulatedB.All.variant.sort.bed.ins.bed -b simulatedBvsB.diff.INTs.bed -f 0.8 -F 0.8 -wa -wb | \
cut -f1-3 | sort | uniq | wc -l | awk -v prefix="TP=" '{print prefix $0}'
## Calculation for FP
$ bedtools intersect -b simulatedB.All.variant.sort.bed.ins.bed -a simulatedBvsB.diff.INTs.bed -f 0.8 -F 0.8 -v | sort | uniq | \
wc -l | awk -v prefix="FP=" '{print prefix $0}'
## Calculation for FN
$ bedtools intersect -a simulatedB.All.variant.sort.bed.ins.bed -b simulatedBvsB.diff.INTs.bed -f 0.8 -F 0.8 -v | sort | uniq | \
wc -l | awk -v prefix="FN=" '{print prefix $0}'
```

**Step 5: Normalization**

We used the alignable proportion to normalize the metrics, ensuring that the precision, recall, and F1-score are calculated only from the alignable regions. These metrics were further adjusted using the genome alignment rate, as described below:

### $average\_AlignmentRate = \ / N_{(\text{The number of aligners})}$

### $adjustTP = TP / average\_AlignmentRate$

### $adjustFP = FP * average\_AlignmentRate$

### $adjustFN = N_{(\text{SV number})} - adjustTP$

### Supplementary Tables

**Supplementary Table 1: Species selected for testing and comparison of aligners in whole genome alignment of plant genomes.**

| Species | Genome size | Repeat content | Assemblies used | Reference |
|---|---|---|---|---|
| Fly (*Drosophila melanogaster)* | ~140 Mb | 22 % | iso-1 *versus* A4 | [2–4] |
| Human (*Homo sapiens*) | ~3.1 Gb | ~54 % | CHM13 *versus* GRCh38 | [5,6] |
| Rice (*Oryza sativa*) | ~396 Mb | ~46 % | MH63 *versus* ZS97 | [7] |
| Tomato (*Solanum lycopersicum*) | ~782 Mb | 71.77 % | SL4.0 *versus* M82 | [8,9] |
| Maize (*Zea mays*) | ~2.2 Gb | ~85 % | B73 *versus* Nc350 | [10] |
| Pepper (*Capsicum annuum*) | ~3.1 Gb | 84.71 % | Ca59 *versus* Zunla-1 | [11,12] |

**Supplementary Table 2: Parameters and commands used for different aligners to evaluate whole genome alignments across genomes of varying complexity.**

| Aligner | Version | Parameters | Reference |
|---|---|---|---|
| lastz | 1.04.22 | lastz $REF[multiple] $QRY --format=axt --output=REFvsQRY.axt | Harris 2007 |
| last | 1406 | lastdb -P8 -uNEAR REFdat $REF | [13] |
| | | lastal REFdat QRY > REFvsQRY.maf | |
| MUMmer | v4 | nucmer --prefix=REFvsQRY.mm $REF $QRY | [14] |
| | | nucmer --maxmatch --prefix=REFvsQRY.max $REF $QRY | |
| minimap2 | 2.24-r1122 | minimap2 -c --cs=long -x asm5 $REF $QRY > REFvsQRY.asm5.paf | [15] |
| | | minimap2 -c --cs=long -x asm10 $REF $QRY > REFvsQRY.asm10.paf | |
| | | minimap2 -c --cs=long -x asm20 $REF $QRY > REFvsQRY.asm20.paf | |
| | | minimap2 -c --cs=long $REF $QRY > REFvsQRY.def.paf | |
| AnchorWave | v1.0.1/ v1.2.3 | anchorwave genoAli -i REF.gff3 -as cds.fa -r $REF -a cds.sam -ar REF.sam -s $QRY -n | [16] |
| | | anchors -o anchorwave.maf -f anchorwave.f.maf -v SV.vcf -m 0 -t 4 | |
| | | anchorwave proali -i REF.gff3 -as cds.fa -r $REF -a cds.sam -ar REF.sam -s $QRY -n | |
| | | anchors | |
| GSAlign | v1.0.22 | -o anchorwave.maf -f anchorwave.f.maf -R 1 -Q 1 -m 0 -t 4 | [17] |
| | | GSAlign -r $REF -q $QRY -o REFvsQRY | |

**Supplementary Table 3: Metrics for evaluating different aligners on whole genome alignments across genomes of varying complexity.**

| Species | Compared genomes | Lastz default | Last default | MUMmer4 default | MUMmer4 maxmatch | minimap2 default | minimap2 asm5 | minimap2 asm10 | minimap2 asm20 | AnchorWave genoAli | AnchorWave proali | GSAlign |
|---|---|---|---|---|---|---|---|---|---|---|---|---|
| **Execution time (CPU-time hh:mm:ss)** | | | | | | | | | | | | |
| Fly | ISO1 versus A4 | 3:28:03 | 0:16:05 | 0:10:17 | 1:11:33 | 0:04:44 | 0:03:55 | 0:02:45 | 0:04:43 | 1:04:03 | 1:13:33 | 0:02:07 |
| Rice | MH63 versus ZH97 | 28:35:04 | 2:47:24 | 6:54:34 | 489:37:29 | 1:22:07 | 0:22:20 | 0:19:43 | 0:41:09 | 5:48:20 | 3:44:18 | 0:07:58 |
| Tomato | SL4.0 versus M82 | > 2 weeks | 7:27:13 | 7:58:48 | NA | 4:00:07 | 0:23:49 | 0:22:05 | 0:41:22 | 6:29:18 | 1:29:57 | 0:14:53 |
| Maize | B73 versus Nc350 | NA | 48:07:32 | 147:10:12 | NA | 164:18:02 | 7:50:44 | 7:15:59 | 13:38:19 | 91:38:35 | 85:39:06 | 1:14:12 |
| Pepper | CA59 versus Zunla-1 | NA | 45:52:15 | 29:58:14 | NA | 23:09:53 | 2:40:44 | 2:38:05 | 5:06:50 | NA | 81:05:16 | 1:05:57 |
| Human | CHM13 versus GRCh38 | NA | 29:28:26 | 17:23:35 | NA | 16:20:02 | 0:43:04 | 0:42:11 | 1:17:29 | 91:28:32 | 55:02:40 | 1:13:54 |
| **Peak memory usage (Gb)** | | | | | | | | | | | | |
| Fly | ISO1 versus A4 | 0.95 | 1.18 | 2.79 | 8.34 | 4.65 | 4.004 | 3.992 | 4.43 | 117.8 | 90.153 | 1.17 |
| Rice | MH63 versus ZH97 | 2.91 | 5.76 | 7.63 | 457.4 | 28 | 10.84 | 10.83 | 19.49 | 145.7 | 116.06 | 2.65 |
| Tomato | SL4.0 versus M82 | 10.3 | 8.75 | 10.78 | NA | 55.9 | 15.15 | 15.02 | 26.1 | 96.1 | 58.57 | 4.4 |
| Maize | B73 versus Nc350 | NA | 38.8 | 30.921 | NA | 280.5 | 38.42 | 36.62 | 66 | 202.2 | 200.78 | 14.19 |
| Pepper | CA59 versus Zunla-1 | NA | 41.32 | 50.61 | NA | 64.49 | 16.17 | 15.92 | 28.97 | NA | 203.31 | 15 |
| Human | CHM13 versus GRCh38 | NA | 31.48 | 48.3 | NA | 147.41 | 25.68 | 25.72 | 36.6 | 210.04 | 209.6 | 14.63 |
| **Number/storage space (Gb) of preliminary alignment blocks** | | | | | | | | | | | | |
| Fly | ISO1 versus A4 | 0.49 | 2.765 | 0.383 | 2.24 | 0.262 | 0.255 | 0.258 | 0.26 | 0.266 | 0.26 | 0.235 |
| Rice | MH63 versus ZH97 | 4.38 | 128.61 | 5.074 | 182.99 | 0.754 | 0.70 | 0.714 | 0.718 | 0.845 | 0.826 | 0.645 |
| Tomato | SL4.0 versus M82 | 93.17 | 450.68 | 8.28 | NA | 1.49 | 1.452 | 1.47 | 1.48 | 1.55 | 1.51 | 1.38 |
| Maize | B73 versus Nc350 | NA | 5,020.0 | 137.00 | NA | 4.87 | 3.50 | 3.834 | 3.95 | 5.67 | 5.6 | 3.04 |
| Pepper | CA59 versus Zunla-1 | NA | 3185.29 | 28.89 | NA | 6.56 | 6.41 | 6.48 | 6.51 | NA | 5.20 | 2.55 |
| Human | CHM13 versus GRCh38 | NA | 1308.86 | 18.25 | NA | 5.56 | 5.48 | 5.54 | 5.63 | 5.63 | 5.39 | 5.19 |
| **Genomic occupancy of the inferred one-to-one orthologous alignments** | | | | | | | | | | | | |
| Fly | ISO1 versus A4 | 0.9522 | 0.9925 | 0.9820 | 0.9926 | 0.9500 | 0.9394 | 0.9434 | 0.9462 | 0.9337 | 0.9228 | 0.9225 |
| Rice | MH63 versus ZH97 | 0.8728 | 0.9827 | 0.9641 | 0.9769 | 0.8724 | 0.8470 | 0.8553 | 0.8558 | 0.8541 | 0.8498 | 0.8305 |
| Tomato | SL4.0 versus M82 | NA | 0.9937 | 0.9895 | NA | 0.9615 | 0.9524 | 0.9548 | 0.9543 | 0.9480 | 0.9474 | 0.9326 |
| Maize | B73 versus Nc350 | NA | NA | NA | NA | 0.6528 | 0.5578 | 0.5789 | 0.5808 | 0.5883 | 0.5910 | 0.6461 |

| | | | | | | | | | | | | |
|---|---|---|---|---|---|---|---|---|---|---|---|---|
| Pepper | CA59 versus Zunla-1 | NA | NA | NA | NA | 0.8905 | 0.8829 | 0.8867 | 0.8873 | NA | 0.3520 | 0.4395 |
| Human | CHM13 versus GRCh38 | NA | NA | NA | NA | 0.9232 | 0.9192 | 0.9212 | 0.9234 | 0.8385 | 0.8350 | 0.8883 |

**Supplementary Table 4: SVs (deletions and insertions) and their overlap across different aligners, as detected by SVGAP between two real genomes within species: rice, *Drosophila*, and tomato.**

| Aligners | AnchorWave genoAli | AnchorWave proAli | GSAlign | last | lastz | minimap2 asm10 | minimap2 asm20 | minimap2 asm5 | minimap2 default | MUMmer4 maxmatch | MUMmer4 default |
|---|---|---|---|---|---|---|---|---|---|---|---|
| **The number of deletions (≥ 50 bp) detected between two rice genomes: MH63 and ZS97.** | | | | | | | | | | | |
| AnchorWave_genoAli | 11,642 | 9,777 | 3,187 | 8,094 | 7,205 | 6,196 | 6,646 | 5,178 | 7,512 | 6,856 | 6,684 |
| AnchorWave_proAli | 9,777 | 10,800 | 3,096 | 7,711 | 6,937 | 5,937 | 6,386 | 5,001 | 7,250 | 6,666 | 6,516 |
| GSAlign | 3,187 | 3,096 | 3,735 | 3,064 | 2,820 | 2,870 | 2,912 | 2,645 | 3,067 | 2,951 | 3,028 |
| last | 8,094 | 7,711 | 3,064 | 10,757 | 6,896 | 6,371 | 6,648 | 5,447 | 7,481 | 7,105 | 6,825 |
| lastz | 7,205 | 6,937 | 2,820 | 6,896 | 9,600 | 5,630 | 5,986 | 4,790 | 6,809 | 5,960 | 5,955 |
| minimap2_asm10 | 6,196 | 5,937 | 2,870 | 6,371 | 5,630 | 8,118 | 7,447 | 6,520 | 7,238 | 6,093 | 6,050 |
| minimap2_asm20 | 6,646 | 6,386 | 2,912 | 6,648 | 5,986 | 7,447 | 8,860 | 6,319 | 7,776 | 6,309 | 6,232 |
| minimap2_asm5 | 5,178 | 5,001 | 2,645 | 5,447 | 4,790 | 6,520 | 6,319 | 6,683 | 6,147 | 5,417 | 5,409 |
| minmap2_default | 7,512 | 7,250 | 3,067 | 7,481 | 6,809 | 7,238 | 7,776 | 6,147 | 10,427 | 6,908 | 6,850 |
| MUMmer4_max | 6,856 | 6,666 | 2,951 | 7,105 | 5,960 | 6,093 | 6,309 | 5,417 | 6,908 | 8,969 | 7,310 |
| MUMmer4_mm | 6,684 | 6,516 | 3,028 | 6,825 | 5,955 | 6,050 | 6,232 | 5,409 | 6,850 | 7,310 | 7,970 |
| **Overlap of deletion calls (≥ 50 bp) across different aligners, as detected by SVGAP between two rice genomes: MH63 and ZS97.** | | | | | | | | | | | |
| AnchorWave_genoAli | 1.0000 | 0.8398 | 0.2738 | 0.6952 | 0.6189 | 0.5322 | 0.5709 | 0.4448 | 0.6452 | 0.5889 | 0.5741 |
| AnchorWave_proAli | 0.9053 | 1.0000 | 0.2867 | 0.7140 | 0.6423 | 0.5497 | 0.5913 | 0.4631 | 0.6713 | 0.6172 | 0.6033 |
| GSAlign | 0.8533 | 0.8289 | 1.0000 | 0.8203 | 0.7550 | 0.7684 | 0.7797 | 0.7082 | 0.2840 | 0.2732 | 0.8107 |
| last | 0.7524 | 0.7168 | 0.2848 | 1.0000 | 0.6411 | 0.5923 | 0.6180 | 0.5064 | 0.6955 | 0.6605 | 0.6345 |
| lastz | 0.7505 | 0.7226 | 0.2938 | 0.7183 | 1.0000 | 0.5865 | 0.6235 | 0.4990 | 0.7093 | 0.6208 | 0.6203 |
| minimap2_asm10 | 0.7632 | 0.7313 | 0.3535 | 0.7848 | 0.6935 | 1.0000 | 0.9173 | 0.8032 | 0.8916 | 0.7506 | 0.7453 |
| minimap2_asm20 | 0.7501 | 0.7208 | 0.3287 | 0.7503 | 0.6756 | 0.8405 | 1.0000 | 0.7132 | 0.8777 | 0.7121 | 0.7034 |
| minimap2_asm5 | 0.7748 | 0.7483 | 0.3958 | 0.8151 | 0.7167 | 0.9756 | 0.9455 | 1.0000 | 0.9198 | 0.8106 | 0.8094 |
| minmap2_default | 0.7204 | 0.6953 | 0.2941 | 0.7175 | 0.6530 | 0.6942 | 0.7458 | 0.5895 | 1.0000 | 0.6625 | 0.6569 |
| MUMmer4_max | 0.7644 | 0.7432 | 0.3290 | 0.7922 | 0.6645 | 0.6793 | 0.7034 | 0.6040 | 0.7702 | 1.0000 | 0.8150 |
| MUMmer4_mm | 0.8386 | 0.8176 | 0.3799 | 0.8563 | 0.7472 | 0.7591 | 0.7819 | 0.6787 | 0.8595 | 0.9172 | 1.0000 |
| **The number of insertions (≥ 50 bp) detected between two rice genomes: MH63 and ZS97.** | | | | | | | | | | | |

| | | | | | | | | | | | |
|---|---|---|---|---|---|---|---|---|---|---|---|
| AnchorWave_genoAli | 11,542 | 9,899 | 3,124 | 8,169 | 7,400 | 6,269 | 6,696 | 5,226 | 7,471 | 6,877 | 6,723 |
| AnchorWave_proAli | 9,899 | 10,879 | 3,067 | 7,810 | 7,158 | 6,048 | 6,486 | 5,099 | 7,250 | 6,763 | 6,626 |
| GSAlign | 3,124 | 3,067 | 3,821 | 3,176 | 2,891 | 3,047 | 3,092 | 2,825 | 3,229 | 3,068 | 3,120 |
| last | 8,169 | 7,810 | 3,176 | 10,484 | 7,084 | 6,509 | 6,801 | 5,543 | 7,514 | 7,181 | 6,914 |
| lastz | 7,400 | 7,158 | 2,891 | 7,084 | 9,860 | 5,813 | 6,211 | 4,908 | 6,864 | 6,202 | 6,092 |
| minimap2_asm10 | 6,269 | 6,048 | 3,047 | 6,509 | 5,813 | 8,213 | 7,584 | 6,599 | 7,342 | 6,227 | 6,197 |
| minimap2_asm20 | 6,696 | 6,486 | 3,092 | 6,801 | 6,211 | 7,584 | 8,891 | 6,397 | 7,849 | 6,416 | 6,384 |
| minimap2_asm5 | 5,226 | 5,099 | 2,825 | 5,543 | 4,908 | 6,599 | 6,397 | 6,708 | 6,222 | 5,515 | 5,527 |
| minmap2_default | 7,471 | 7,250 | 3,229 | 7,514 | 6,864 | 7,342 | 7,849 | 6,222 | 10,076 | 6,954 | 6,929 |
| MUMmer4_max | 6,877 | 6,763 | 3,068 | 7,181 | 6,202 | 6,227 | 6,416 | 5,515 | 6,954 | 8,665 | 7,533 |
| MUMmer4_mm | 6,723 | 6,626 | 3,120 | 6,914 | 6,092 | 6,197 | 6,384 | 5,527 | 6,929 | 7,533 | 8,097 |
| **Overlap of insertion calls (≥ 50 bp) between different aligners, as detected by SVGAP between two rice genomes: MH63 and ZS97.** | | | | | | | | | | | |
| AnchorWave_genoAli | 1.0000 | 0.8577 | 0.2707 | 0.7078 | 0.6411 | 0.5431 | 0.5801 | 0.4528 | 0.6473 | 0.5958 | 0.5825 |
| AnchorWave_proAli | 0.9099 | 1.0000 | 0.2819 | 0.7179 | 0.6580 | 0.5559 | 0.5962 | 0.4687 | 0.6664 | 0.6217 | 0.6091 |
| GSAlign | 0.8176 | 0.8027 | 1.0000 | 0.8312 | 0.7566 | 0.7974 | 0.8092 | 0.7393 | 0.8451 | 0.8029 | 0.8165 |
| last | 0.7792 | 0.7449 | 0.3029 | 1.0000 | 0.6757 | 0.6209 | 0.6487 | 0.5287 | 0.7167 | 0.6849 | 0.6595 |
| lastz | 0.7505 | 0.7260 | 0.2932 | 0.7185 | 1.0000 | 0.5896 | 0.6299 | 0.4978 | 0.6961 | 0.6290 | 0.6178 |
| minimap2_asm10 | 0.7633 | 0.7364 | 0.3710 | 0.7925 | 0.7078 | 1.0000 | 0.9234 | 0.8035 | 0.8939 | 0.7582 | 0.7545 |
| minimap2_asm20 | 0.7531 | 0.7295 | 0.3478 | 0.7649 | 0.6986 | 0.8530 | 1.0000 | 0.7195 | 0.8828 | 0.7216 | 0.7180 |
| minimap2_asm5 | 0.7791 | 0.7601 | 0.4211 | 0.8263 | 0.7317 | 0.9838 | 0.9536 | 1.0000 | 0.9275 | 0.8222 | 0.8239 |
| minmap2_default | 0.7415 | 0.7195 | 0.3205 | 0.7457 | 0.6812 | 0.7287 | 0.7790 | 0.6175 | 1.0000 | 0.6902 | 0.6877 |
| MUMmer4_max | 0.7937 | 0.7805 | 0.3541 | 0.8287 | 0.7158 | 0.7186 | 0.7405 | 0.6365 | 0.8025 | 1.0000 | 0.8694 |
| MUMmer4_mm | 0.8303 | 0.8183 | 0.3853 | 0.8539 | 0.7524 | 0.7653 | 0.7884 | 0.6826 | 0.8557 | 0.9303 | 1.0000 |
| **The number of deletions (≥ 50 bp) detected between two *Drosophila* genomes: iso-1 and A4.** | | | | | | | | | | | |
| AnchorWave_genoAli | 2,203 | 1,930 | 819 | 1,742 | 1,603 | 1,766 | 1,812 | 1,551 | 1,777 | 1,609 | 1,546 |
| AnchorWave_proAli | 1,930 | 1,999 | 786 | 1,632 | 1,529 | 1,649 | 1,686 | 1,479 | 1,670 | 1,544 | 1,501 |
| GSAlign | 819 | 786 | 984 | 823 | 791 | 836 | 833 | 784 | 840 | 804 | 792 |
| last | 1,742 | 1,632 | 823 | 2,214 | 1,694 | 1,853 | 1,840 | 1,667 | 1,816 | 1,788 | 1,646 |

| | | | | | | | | | | | |
|---|---|---|---|---|---|---|---|---|---|---|---|
| lastz | 1,603 | 1,529 | 791 | 1,694 | 2,156 | 1,661 | 1,682 | 1,495 | 1,670 | 1,568 | 1,497 |
| minimap2_asm10 | 1,766 | 1,649 | 836 | 1,853 | 1,661 | 2,199 | 2,061 | 1,864 | 1,951 | 1,728 | 1,645 |
| minimap2_asm20 | 1,812 | 1,686 | 833 | 1,840 | 1,682 | 2,061 | 2,248 | 1,808 | 2,019 | 1,735 | 1,645 |
| minimap2_asm5 | 1,551 | 1,479 | 784 | 1,667 | 1,495 | 1,864 | 1,808 | 1,905 | 1,747 | 1,629 | 1,552 |
| minmap2_default | 1,777 | 1,670 | 840 | 1,816 | 1,670 | 1,951 | 2,019 | 1,747 | 2,230 | 1,724 | 1,647 |
| MUMmer4_max | 1,609 | 1,544 | 804 | 1,788 | 1,568 | 1,728 | 1,735 | 1,629 | 1,724 | 1,987 | 1,738 |
| MUMmer4_mm | 1,546 | 1,501 | 792 | 1,646 | 1,497 | 1,645 | 1,645 | 1,552 | 1,647 | 1,738 | 1,819 |

**Overlap of deletion calls (≥ 50 bp) between different aligners, as detected by SVGAP between two *Drosophila* genomes: iso-1 and A4.**

| | | | | | | | | | | | |
|---|---|---|---|---|---|---|---|---|---|---|---|
| AnchorWave_genoAli | 1.0000 | 0.8761 | 0.3718 | 0.7907 | 0.7276 | 0.8016 | 0.8225 | 0.7040 | 0.8066 | 0.7304 | 0.7018 |
| AnchorWave_proAli | 0.9655 | 1.0000 | 0.3932 | 0.8164 | 0.7649 | 0.8249 | 0.8434 | 0.7399 | 0.8354 | 0.7724 | 0.7509 |
| GSAlign | 0.8323 | 0.7988 | 1.0000 | 0.8364 | 0.8039 | 0.8496 | 0.8465 | 0.7967 | 0.8537 | 0.8171 | 0.8049 |
| last | 0.7868 | 0.7371 | 0.3717 | 1.0000 | 0.7651 | 0.8369 | 0.8311 | 0.7529 | 0.8202 | 0.8076 | 0.7435 |
| lastz | 0.7435 | 0.7092 | 0.3669 | 0.7857 | 1.0000 | 0.7704 | 0.7801 | 0.6934 | 0.7746 | 0.7273 | 0.6943 |
| minimap2_asm10 | 0.8031 | 0.7499 | 0.3802 | 0.8427 | 0.7553 | 1.0000 | 0.9372 | 0.8477 | 0.8872 | 0.7858 | 0.7481 |
| minimap2_asm20 | 0.8060 | 0.7500 | 0.3706 | 0.8185 | 0.7482 | 0.9168 | 1.0000 | 0.8043 | 0.8981 | 0.7718 | 0.7318 |
| minimap2_asm5 | 0.8142 | 0.7764 | 0.4115 | 0.8751 | 0.7848 | 0.9785 | 0.9491 | 1.0000 | 0.9171 | 0.8551 | 0.8147 |
| minmap2_default | 0.7969 | 0.7489 | 0.3767 | 0.8143 | 0.7489 | 0.8749 | 0.9054 | 0.7834 | 1.0000 | 0.7731 | 0.7386 |
| MUMmer4_max | 0.8098 | 0.7771 | 0.4046 | 0.8998 | 0.7891 | 0.8697 | 0.8732 | 0.8198 | 0.8676 | 1.0000 | 0.8747 |
| MUMmer4_mm | 0.8499 | 0.8252 | 0.4354 | 0.9049 | 0.8230 | 0.9043 | 0.9043 | 0.8532 | 0.9054 | 0.9555 | 1.0000 |

**The number of insertions (≥ 50 bp) detected between two Drosophila genomes: iso-1 and A4.**

| | | | | | | | | | | | |
|---|---|---|---|---|---|---|---|---|---|---|---|
| AnchorWave_genoAli | 2,185 | 1,914 | 766 | 1,737 | 1,609 | 1,752 | 1,778 | 1,543 | 1,758 | 1,595 | 1,533 |
| AnchorWave_proAli | 1,914 | 1,999 | 726 | 1,623 | 1,525 | 1,628 | 1,659 | 1,453 | 1,654 | 1,527 | 1,481 |
| GSAlign | 766 | 726 | 957 | 808 | 767 | 810 | 810 | 755 | 810 | 776 | 778 |
| last | 1,737 | 1,623 | 808 | 2,456 | 1,715 | 1,831 | 1,834 | 1,638 | 1,812 | 1,837 | 1,671 |
| lastz | 1,609 | 1,525 | 767 | 1,715 | 2,245 | 1,683 | 1,715 | 1,494 | 1,699 | 1,574 | 1,509 |
| minimap2_asm10 | 1,752 | 1,628 | 810 | 1,831 | 1,683 | 2,273 | 2,088 | 1,895 | 1,960 | 1,692 | 1,614 |
| minimap2_asm20 | 1,778 | 1,659 | 810 | 1,834 | 1,715 | 2,088 | 2,326 | 1,819 | 2,019 | 1,703 | 1,623 |
| minimap2_asm5 | 1,543 | 1,453 | 755 | 1,638 | 1,494 | 1,895 | 1,819 | 1,958 | 1,736 | 1,554 | 1,500 |

| | | | | | | | | | | | |
|---|---|---|---|---|---|---|---|---|---|---|---|
| minmap2_default | 1,758 | 1,654 | 810 | 1,812 | 1,699 | 1,960 | 2,019 | 1,736 | 2,349 | 1,681 | 1,618 |
| MUMmer4_max | 1,595 | 1,527 | 776 | 1,837 | 1,574 | 1,692 | 1,703 | 1,554 | 1,681 | 2,171 | 1,745 |
| MUMmer4_mm | 1,533 | 1,481 | 778 | 1,671 | 1,509 | 1,614 | 1,623 | 1,500 | 1,618 | 1,745 | 1,908 |
| **Overlap of insertion calls (≥ 50 bp) between different aligners, as detected by SVGAP between two *Drosophila* genomes: iso-1 and A4.** | | | | | | | | | | | |
| AnchorWave_genoAli | 1.0000 | 0.8760 | 0.3506 | 0.7950 | 0.7364 | 0.8018 | 0.8137 | 0.7062 | 0.8046 | 0.7300 | 0.7016 |
| AnchorWave_proAli | 0.9575 | 1.0000 | 0.3632 | 0.8119 | 0.7629 | 0.8144 | 0.8299 | 0.7269 | 0.8274 | 0.7639 | 0.7409 |
| GSAlign | 0.8004 | 0.7586 | 1.0000 | 0.8443 | 0.8015 | 0.8464 | 0.8464 | 0.7889 | 0.8464 | 0.8109 | 0.8130 |
| last | 0.7072 | 0.6608 | 0.3290 | 1.0000 | 0.6983 | 0.7455 | 0.7467 | 0.6669 | 0.7378 | 0.7480 | 0.6804 |
| lastz | 0.7167 | 0.6793 | 0.3416 | 0.7639 | 1.0000 | 0.7497 | 0.7639 | 0.6655 | 0.7568 | 0.7011 | 0.6722 |
| minimap2_asm10 | 0.7708 | 0.7162 | 0.3564 | 0.8055 | 0.7404 | 1.0000 | 0.9186 | 0.8337 | 0.8623 | 0.7444 | 0.7101 |
| minimap2_asm20 | 0.7644 | 0.7132 | 0.3482 | 0.7885 | 0.7373 | 0.8977 | 1.0000 | 0.7820 | 0.8680 | 0.7322 | 0.6978 |
| minimap2_asm5 | 0.7880 | 0.7421 | 0.3856 | 0.8366 | 0.7630 | 0.9678 | 0.9290 | 1.0000 | 0.8866 | 0.7937 | 0.7661 |
| minmap2_default | 0.7484 | 0.7041 | 0.3448 | 0.7714 | 0.7233 | 0.8344 | 0.8595 | 0.7390 | 1.0000 | 0.7156 | 0.6888 |
| MUMmer4_max | 0.7347 | 0.7034 | 0.3574 | 0.8462 | 0.7250 | 0.7794 | 0.7844 | 0.7158 | 0.7743 | 1.0000 | 0.8038 |
| MUMmer4_mm | 0.8035 | 0.7762 | 0.4078 | 0.8758 | 0.7909 | 0.8459 | 0.8506 | 0.7862 | 0.8480 | 0.9146 | 1.0000 |
| **The number of deletions (≥ 50 bp) detected between two tomato genomes: SL4 and M82.** | | | | | | | | | | | |
| AnchorWave_genoAli | 4,654 | 4,036 | 1,160 | 2,448 | 93 | 2,600 | 2,766 | 2,263 | 2,631 | NA | 2,209 |
| AnchorWave_proAli | 4,036 | 4,398 | 1,144 | 2,381 | 91 | 2,491 | 2,660 | 2,198 | 2,548 | NA | 2,178 |
| GSAlign | 1,160 | 1,144 | 1,504 | 1,124 | 35 | 1,159 | 1,154 | 1,106 | 1,157 | NA | 1,138 |
| last | 2,448 | 2,381 | 1,124 | 4,135 | 104 | 2,511 | 2,517 | 2,266 | 2,520 | NA | 2,363 |
| lastz | 93 | 91 | 35 | 104 | 213 | 114 | 121 | 94 | 122 | NA | 85 |
| minimap2_asm10 | 2,600 | 2,491 | 1,159 | 2,511 | 114 | 4,540 | 4,112 | 3,533 | 3,809 | NA | 2,450 |
| minimap2_asm20 | 2,766 | 2,660 | 1,154 | 2,517 | 121 | 4,112 | 5,020 | 3,406 | 4,210 | NA | 2,475 |
| minimap2_asm5 | 2,263 | 2,198 | 1,106 | 2,266 | 94 | 3,533 | 3,406 | 3,618 | 3,205 | NA | 2,280 |
| minmap2_default | 2,631 | 2,548 | 1,157 | 2,520 | 122 | 3,809 | 4,210 | 3,205 | 5,398 | NA | 2,483 |
| MUMmer4_default | 2,209 | 2,178 | 1,138 | 2,363 | 85 | 2,450 | 2,475 | 2,280 | 2,483 | NA | 3,083 |
| **Overlap of deletion calls (≥ 50 bp) between different aligners, as detected by SVGAP between two tomato genomes: SL4 and M82.** | | | | | | | | | | | |
| AnchorWave_genoAli | 1.0000 | 0.8672 | 0.2492 | 0.5260 | 0.0200 | 0.5587 | 0.5943 | 0.4862 | 0.5653 | NA | 0.4746 |

| | | | | | | | | | | | |
|---|---|---|---|---|---|---|---|---|---|---|---|
| AnchorWave_proAli | 0.9177 | 1.0000 | 0.2601 | 0.5414 | 0.0207 | 0.5664 | 0.6048 | 0.4998 | 0.5794 | NA | 0.4952 |
| GSAlign | 0.7713 | 0.7606 | 1.0000 | 0.7473 | 0.0233 | 0.7706 | 0.7673 | 0.7354 | 0.7693 | NA | 0.7566 |
| last | 0.5920 | 0.5758 | 0.2718 | 1.0000 | 0.0252 | 0.6073 | 0.6087 | 0.5480 | 0.6094 | NA | 0.5715 |
| lastz | 0.4366 | 0.4272 | 0.1643 | 0.4883 | 1.0000 | 0.5352 | 0.5681 | 0.4413 | 0.5728 | NA | 0.3991 |
| minimap2_asm10 | 0.5727 | 0.5487 | 0.2553 | 0.5531 | 0.0251 | 1.0000 | 0.9057 | 0.7782 | 0.8390 | NA | 0.5396 |
| minimap2_asm20 | 0.5510 | 0.5299 | 0.2299 | 0.5014 | 0.0241 | 0.8191 | 1.0000 | 0.6785 | 0.8386 | NA | 0.4930 |
| minimap2_asm5 | 0.6255 | 0.6075 | 0.3057 | 0.6263 | 0.0260 | 0.9765 | 0.9414 | 1.0000 | 0.8858 | NA | 0.6302 |
| minmap2_default | 0.4874 | 0.4720 | 0.2143 | 0.4668 | 0.0226 | 0.7056 | 0.7799 | 0.5937 | 1.0000 | NA | 0.4600 |
| MUMmer4_default | 0.7165 | 0.7065 | 0.3691 | 0.7665 | 0.0276 | 0.7947 | 0.8028 | 0.7395 | 0.8054 | NA | 1.0000 |

**The number of insertions (≥ 50 bp) detected between two tomato genomes: SL4 and M82.**

| | | | | | | | | | | | |
|---|---|---|---|---|---|---|---|---|---|---|---|
| AnchorWave_genoAli | 4,814 | 4,282 | 1,309 | 3,209 | 98 | 3,108 | 3,229 | 2,829 | 3,228 | NA | 2,934 |
| AnchorWave_proAli | 4,282 | 4,588 | 1,287 | 3,109 | 92 | 3,011 | 3,120 | 2,769 | 3,136 | NA | 2,900 |
| GSAlign | 1,309 | 1,287 | 1,754 | 1,429 | 46 | 1,441 | 1,461 | 1,385 | 1,457 | NA | 1,426 |
| last | 3,209 | 3,109 | 1,429 | 6,100 | 125 | 3,256 | 3,290 | 3,045 | 3,331 | NA | 3,159 |
| lastz | 98 | 92 | 46 | 125 | 194 | 103 | 110 | 85 | 106 | NA | 89 |
| minimap2_asm10 | 3,108 | 3,011 | 1,441 | 3,256 | 103 | 4,223 | 3,941 | 3,644 | 3,709 | NA | 3,161 |
| minimap2_asm20 | 3,229 | 3,120 | 1,461 | 3,290 | 110 | 3,941 | 4,562 | 3,538 | 3,865 | NA | 3,199 |
| minimap2_asm5 | 2,829 | 2,769 | 1,385 | 3,045 | 85 | 3,644 | 3,538 | 3,713 | 3,378 | NA | 3,008 |
| minmap2_default | 3,228 | 3,136 | 1,457 | 3,331 | 106 | 3,709 | 3,865 | 3,378 | 5,181 | NA | 3,238 |
| MUMmer4_default | 2,934 | 2,900 | 1,426 | 3,159 | 89 | 3,161 | 3,199 | 3,008 | 3,238 | NA | 3,796 |

**Overlap of insertion calls (≥ 50 bp) between different aligners, as detected by SVGAP between two tomato genomes: SL4 and M82.**

| | | | | | | | | | | | |
|---|---|---|---|---|---|---|---|---|---|---|---|
| AnchorWave_genoAli | 1.0000 | 0.8895 | 0.2719 | 0.6666 | 0.0204 | 0.6456 | 0.6708 | 0.5877 | 0.6705 | NA | 0.6095 |
| AnchorWave_proAli | 0.9333 | 1.0000 | 0.2805 | 0.6776 | 0.0201 | 0.6563 | 0.6800 | 0.6035 | 0.6835 | NA | 0.6321 |
| GSAlign | 0.7463 | 0.7338 | 1.0000 | 0.8147 | 0.0262 | 0.8216 | 0.8330 | 0.7896 | 0.8307 | NA | 0.8130 |
| last | 0.5261 | 0.5097 | 0.2343 | 1.0000 | 0.0205 | 0.5338 | 0.5393 | 0.4992 | 0.5461 | NA | 0.5179 |
| lastz | 0.5052 | 0.4742 | 0.2371 | 0.6443 | 1.0000 | 0.5309 | 0.5670 | 0.4381 | 0.5464 | NA | 0.4588 |
| minimap2_asm10 | 0.7360 | 0.7130 | 0.3412 | 0.7710 | 0.0244 | 1.0000 | 0.9332 | 0.8629 | 0.8783 | NA | 0.7485 |
| minimap2_asm20 | 0.7078 | 0.6839 | 0.3203 | 0.7212 | 0.0241 | 0.8639 | 1.0000 | 0.7755 | 0.8472 | NA | 0.7012 |

| | | | | | | | | | | | |
|---|---|---|---|---|---|---|---|---|---|---|---|
| minimap2_asm5 | 0.7619 | 0.7458 | 0.3730 | 0.8201 | 0.0229 | 0.9814 | 0.9529 | 1.0000 | 0.9098 | NA | 0.8101 |
| minmap2_default | 0.6230 | 0.6053 | 0.2812 | 0.6429 | 0.0205 | 0.7159 | 0.7460 | 0.6520 | 1.0000 | NA | 0.6250 |
| MUMmer4_default | 0.7729 | 0.7640 | 0.3757 | 0.8322 | 0.0234 | 0.8327 | 0.8427 | 0.7924 | 0.8530 | NA | 1.0000 |

**Supplementary Table 5: Benchmark results with reference-based simulated SVs in tomato, rice, and *Drosophila*.**

| Aligner | # of total calls | # of accurate calls | Recall (Sensitivity) | Precision | FDR | F1 |
|---|---|---|---|---|---|---|
| *Simulated **deletions** in tomato (assembly = SL4; n = 28,440; size ≥ 50 bp)* | | | | | | |
| AnchorWave_genoAli | 28,060 | 27,490 | 0.9666 | 0.9797 | 0.0203 | 0.9731 |
| AnchroWave_proAli | 27,693 | 26,230 | 0.9223 | 0.9472 | 0.0528 | 0.9346 |
| GSAlign | 26,215 | 26,167 | 0.9201 | 0.9982 | 0.0018 | 0.9575 |
| **last** | **28,225** | **28,102** | **0.9881** | **0.9956** | **0.0044** | **0.9919** |
| lastz | 21,521 | 19,909 | 0.7000 | 0.9251 | 0.0749 | 0.7970 |
| minimap2_asm10 | 27,562 | 27,466 | 0.9658 | 0.9965 | 0.0035 | 0.9809 |
| minimap2_asm20 | 27,573 | 27,443 | 0.9649 | 0.9953 | 0.0047 | 0.9799 |
| minimap2_asm5 | 27,542 | 27,452 | 0.9653 | 0.9967 | 0.0033 | 0.9807 |
| minimap2_default | 28,323 | 27,304 | 0.9601 | 0.9640 | 0.0360 | 0.9620 |
| MUMmer4_default | 27,913 | 27,857 | 0.9795 | 0.9980 | 0.0020 | 0.9887 |
| *Simulated **insertions** in tomato (assembly = SL4; n = 15,245; size ≥ 50 bp)* | | | | | | |
| AnchorWave_genoAli | 15,271 | 14,594 | 0.9573 | 0.9557 | 0.0443 | 0.9565 |
| AnchroWave_proAli | 15,293 | 14,318 | 0.9392 | 0.9362 | 0.0638 | 0.9377 |
| GSAlign | 14,387 | 14,385 | 0.9436 | 0.9999 | 0.0001 | 0.9709 |
| **last** | **15,137** | **15,098** | **0.9904** | **0.9974** | **0.0026** | **0.9939** |
| lastz | 12,644 | 11,918 | 0.7818 | 0.9426 | 0.0574 | 0.8547 |
| minimap2_asm10 | 14,651 | 14,627 | 0.9595 | 0.9984 | 0.0016 | 0.9785 |
| minimap2_asm20 | 14,662 | 14,614 | 0.9586 | 0.9967 | 0.0033 | 0.9773 |
| minimap2_asm5 | 14,641 | 14,620 | 0.9590 | 0.9986 | 0.0014 | 0.9784 |
| minimap2_default | 14,931 | 14,645 | 0.9606 | 0.9808 | 0.0192 | 0.9706 |
| MUMmer4_default | 15,066 | 15,042 | 0.9867 | 0.9984 | 0.0016 | 0.9925 |
| *Simulated **deletions** in rice (assembly = Nipponbare; n = 20,328; size ≥ 50 bp)* | | | | | | |
| AnchorWave_genoAli | 19,805 | 19,452 | 0.9569 | 0.9822 | 0.0178 | 0.9694 |
| AnchroWave_proAli | 19,086 | 18,308 | 0.9006 | 0.9592 | 0.0408 | 0.9290 |
| GSAlign | 17,796 | 17,737 | 0.8725 | 0.9967 | 0.0033 | 0.9305 |
| last | **19,974** | **19,828** | **0.9754** | **0.9927** | **0.0073** | **0.9840** |
| lastz | 15,648 | 14,908 | 0.7334 | 0.9527 | 0.0473 | 0.8288 |
| minimap2_asm10 | 18,980 | 18,882 | 0.9289 | 0.9948 | 0.0052 | 0.9607 |
| minimap2_asm20 | 18,956 | 18,830 | 0.9263 | 0.9934 | 0.0066 | 0.9587 |

| | | | | | | |
|---|---|---|---|---|---|---|
| minimap2_asm5 | 18,941 | 18,848 | 0.9272 | 0.9951 | 0.0049 | 0.9599 |
| minimap2_default | 19,673 | 19,197 | 0.9444 | 0.9758 | 0.0242 | 0.9598 |
| MUMmer4_default | 19,344 | 19,279 | 0.9484 | 0.9966 | 0.0034 | 0.9719 |
| MUMmer4_maxmatch | 19,502 | 19,359 | 0.9523 | 0.9927 | 0.0073 | 0.9721 |
| unimap | 17,785 | 17,693 | 0.8704 | 0.9948 | 0.0052 | 0.9284 |
| winnowmap | 15,870 | 15,801 | 0.7773 | 0.9957 | 0.0043 | 0.8730 |
| *Simulated* ***insertions*** *in rice (assembly* = *Nipponbare; n* = 10,046*; size ≥ 50 bp)* | | | | | | |
| AnchorWave_genoAli | 9,987 | 9,641 | 0.9597 | 0.9654 | 0.0346 | 0.9625 |
| AnchorWave_proAli | 9,971 | 9,384 | 0.9341 | 0.9411 | 0.0589 | 0.9376 |
| GSAlign | 9,350 | 9,336 | 0.9293 | 0.9985 | 0.0015 | 0.9627 |
| last | **9,887** | **9,855** | **0.9810** | **0.9968** | **0.0032** | **0.9888** |
| lastz | 8,241 | 8,088 | 0.8051 | 0.9814 | 0.0186 | 0.8846 |
| minimap2_asm10 | 9,249 | 9,229 | 0.9187 | 0.9978 | 0.0022 | 0.9566 |
| minimap2_asm20 | 9,239 | 9,205 | 0.9163 | 0.9963 | 0.0037 | 0.9546 |
| minimap2_asm5 | 9,240 | 9,226 | 0.9184 | 0.9985 | 0.0015 | 0.9568 |
| minimap2_default | 9,589 | 9,416 | 0.9373 | 0.9820 | 0.0180 | 0.9591 |
| MUMmer4_default | 9,757 | 9,744 | 0.9699 | 0.9987 | 0.0013 | 0.9841 |
| MUMmer4_maxmatch | 9,757 | 9,712 | 0.9668 | 0.9954 | 0.0046 | 0.9809 |
| unimap | 8,470 | 8,457 | 0.8418 | 0.9985 | 0.0015 | 0.9135 |
| winnowmap | 8,276 | 8,264 | 0.8226 | 0.9986 | 0.0014 | 0.9021 |
| *Simulated* ***deletions*** *in Drosophila (assembly* = iso-1*; n* = 7,264*; size ≥ 50 bp)* | | | | | | |
| AnchorWave_genoAli | 7,001 | 6,935 | 0.9547 | 0.9906 | 0.0094 | 0.9723 |
| AnchorWave_proAli | 6,777 | 6,588 | 0.9069 | 0.9721 | 0.0279 | 0.9384 |
| GSAlign | 6,415 | 6,399 | 0.8809 | 0.9975 | 0.0025 | 0.9356 |
| last | **7,200** | **7,125** | **0.9809** | **0.9896** | **0.0104** | **0.9852** |
| lastz | 6,743 | 6,541 | 0.9005 | 0.9700 | 0.0300 | 0.9340 |
| minimap2_asm10 | 6,994 | 6,951 | 0.9569 | 0.9939 | 0.0061 | 0.9750 |
| minimap2_asm20 | 6,981 | 6,916 | 0.9521 | 0.9907 | 0.0093 | 0.9710 |
| minimap2_asm5 | 6,985 | 6,945 | 0.9561 | 0.9943 | 0.0057 | 0.9748 |
| minimap2_default | 7,112 | 7,015 | 0.9657 | 0.9864 | 0.0136 | 0.9759 |
| MUMmer4_default | 6,957 | 6,941 | 0.9555 | 0.9977 | 0.0023 | 0.9762 |
| MUMmer4_maxmatch | 7,117 | 7,042 | 0.9694 | 0.9895 | 0.0105 | 0.9793 |

| *Simulated **insertions** in Drosophila (assembly* = iso-1*; n* = 4,290*; size ≥ 50 bp)* | | | | | | |
|---|---|---|---|---|---|---|
| AnchorWave_genoAli | 4,156 | 3,874 | 0.9030 | 0.9321 | 0.0679 | 0.9174 |
| AnchorWave_proAli | 4,099 | 3,760 | 0.8765 | 0.9173 | 0.0827 | 0.8964 |
| GSAlign | 3,928 | 3,923 | 0.9145 | 0.9987 | 0.0013 | 0.9547 |
| last | **4,246** | **4,211** | **0.9816** | **0.9918** | **0.0082** | **0.9866** |
| lastz | 4,083 | 3,978 | 0.9273 | 0.9743 | 0.0257 | 0.9502 |
| minimap2_asm10 | 4,099 | 4,070 | 0.9487 | 0.9929 | 0.0071 | 0.9703 |
| minimap2_asm20 | 4,080 | 4,039 | 0.9415 | 0.9900 | 0.0100 | 0.9651 |
| minimap2_asm5 | 4,086 | 4,066 | 0.9478 | 0.9951 | 0.0049 | 0.9709 |
| minimap2_default | 4,171 | 4,062 | 0.9469 | 0.9739 | 0.0261 | 0.9602 |
| MUMmer4_default | 4,165 | 4,154 | 0.9683 | 0.9974 | 0.0026 | 0.9826 |
| MUMmer4_maxmatch | 4,230 | 4,198 | 0.9786 | 0.9924 | 0.0076 | 0.9854 |

**Supplementary Table 6: Benchmark results across aligners and callers with phylogeny-based simulated SVs in rice and tomato.**

| Rice | MH63 | R498 | 9311 | ZS97 | Nipp | Oruf | CJ14 | Opun |
|---|---|---|---|---|---|---|---|---|
| ***Recall without normalization for simulated 10,000 deletions in different rice genomes (size ≥ 50 bp)*** | | | | | | | | |
| minimap2 | 0.9803 | 0.8828 | 0.8266 | 0.8089 | 0.6862 | 0.5645 | 0.6464 | 0.1602 |
| MUMmer4 | 0.976 | 0.8811 | 0.8221 | 0.8035 | 0.6793 | 0.5637 | 0.6342 | 0.1283 |
| AnchorWave | 0.9853 | 0.8887 | 0.8343 | 0.8196 | 0.6873 | 0.5785 | 0.6632 | 0.1961 |
| last | 0.9915 | 0.8922 | 0.8384 | 0.8216 | 0.6897 | 0.5786 | 0.6687 | 0.205 |
| last-split | 0.9869 | 0.8891 | 0.837 | 0.8175 | 0.6914 | 0.5791 | 0.6572 | 0.1985 |
| Anchorwave | 0.9852 | 0.8888 | 0.8344 | 0.8195 | 0.6868 | 0.579 | 0.6622 | 0.1966 |
| SVIM-asm | 0.9711 | 0.8737 | 0.8183 | 0.7992 | 0.6712 | 0.5549 | 0.6324 | 0.1427 |
| paftools | 0.9655 | 0.8562 | 0.792 | 0.7692 | 0.6093 | 0.4784 | 0.5558 | 0.0168 |
| Assemblytics | 0.5934 | 0.5464 | 0.5173 | 0.4988 | 0.4417 | 0.3756 | 0.4149 | 0.0863 |
| SyRI | 0.5857 | 0.526 | 0.5089 | 0.4844 | 0.4223 | 0.3584 | 0.3858 | 0.0252 |
| GSAlign | 0.0018 | 0.0132 | 0.0178 | 0.0222 | 0.0426 | 0.0408 | 0.0527 | 0.0152 |
| MUM&CO | 0.7468 | 0.6285 | 0.5597 | 0.546 | 0.3715 | 0.287 | 0.2378 | 0 |
| ***Precision without normalization for simulated 10,000 deletions in different rice genomes (size ≥ 50 bp)*** | | | | | | | | |
| minimap2 | 0.9849 | 0.9540 | 0.9304 | 0.8871 | 0.8078 | 0.7923 | 0.7991 | 0.3518 |
| MUMmer4 | 0.9950 | 0.9911 | 0.9850 | 0.9807 | 0.9031 | 0.9461 | 0.9573 | 0.7752 |
| Anchorwave | 0.9842 | 0.9578 | 0.9369 | 0.9020 | 0.8451 | 0.8410 | 0.8629 | 0.5324 |
| last | 0.9961 | 0.9646 | 0.9100 | 0.8730 | 0.8614 | 0.8784 | 0.8376 | 0.5963 |
| last-split | 0.9962 | 0.9821 | 0.9636 | 0.9649 | 0.9283 | 0.9316 | 0.9070 | 0.6803 |
| Anchorwave | 0.9822 | 0.9489 | 0.9275 | 0.8885 | 0.8396 | 0.8085 | 0.8337 | 0.4458 |
| SVIM-asm | 0.9720 | 0.8349 | 0.8182 | 0.7624 | 0.6642 | 0.5666 | 0.6130 | 0.2039 |
| paftools | 0.9698 | 0.9240 | 0.8979 | 0.8591 | 0.7896 | 0.6706 | 0.7065 | 0.0680 |
| Assemblytics | 0.8716 | 0.9476 | 0.9287 | 0.9188 | 0.8613 | 0.8268 | 0.8028 | 0.6426 |
| SyRI | 0.9894 | 0.9781 | 0.9658 | 0.9636 | 0.9302 | 0.9057 | 0.8820 | 0.7500 |
| GSAlign | 0.0455 | 1.0000 | 1.0000 | 1.0000 | 1.0000 | 1.0000 | 1.0000 | 1.0000 |
| MUM&co | 0.9624 | 0.9190 | 0.8698 | 0.8370 | 0.6834 | 0.5879 | 0.5141 | 0.0000 |
| ***F1 score without normalization for simulated 10,000 deletions in different rice genomes (size ≥ 50 bp)*** | | | | | | | | |
| minimap2 | 0.9826 | 0.9170 | 0.8755 | 0.8462 | 0.7420 | 0.6593 | 0.7147 | 0.2201 |
| MUMmer4 | 0.9854 | 0.9329 | 0.8962 | 0.8833 | 0.7754 | 0.7065 | 0.7629 | 0.2202 |
| Anchorwave | 0.9848 | 0.9219 | 0.8826 | 0.8588 | 0.7581 | 0.6855 | 0.7500 | 0.2866 |
| last | 0.9938 | 0.9270 | 0.8727 | 0.8465 | 0.7660 | 0.6977 | 0.7437 | 0.3051 |
| last-split | 0.9915 | 0.9333 | 0.8959 | 0.8851 | 0.7925 | 0.7142 | 0.7621 | 0.3073 |
| Anchorwave | 0.9837 | 0.9178 | 0.8785 | 0.8526 | 0.7556 | 0.6748 | 0.7381 | 0.2729 |
| SVIM-asm | 0.9715 | 0.8538 | 0.8183 | 0.7804 | 0.6677 | 0.5607 | 0.6226 | 0.1679 |
| paftools | 0.9676 | 0.8888 | 0.8416 | 0.8116 | 0.6878 | 0.5584 | 0.6222 | 0.0269 |
| Assemblytics | 0.7061 | 0.6931 | 0.6645 | 0.6466 | 0.5840 | 0.5165 | 0.5471 | 0.1522 |
| SyRI | 0.7358 | 0.6841 | 0.6666 | 0.6447 | 0.5809 | 0.5136 | 0.5368 | 0.0488 |
| GSAlign | 0.0035 | 0.0261 | 0.0350 | 0.0434 | 0.0817 | 0.0784 | 0.1001 | 0.0299 |
| MUM&co | 0.8410 | 0.7465 | 0.6811 | 0.6609 | 0.4813 | 0.3857 | 0.3252 | 0.0000 |
| ***Recall without normalization for simulated 10,000 insertions in different rice genomes (size ≥ 50 bp)*** | | | | | | | | |
| minimap2 | 0.9847 | 0.9101 | 0.8679 | 0.847 | 0.7714 | 0.6571 | 0.7477 | 0.2795 |
| MUMmer4 | 0.9945 | 0.9243 | 0.8829 | 0.8662 | 0.7736 | 0.6855 | 0.7781 | 0.2959 |

| | | | | | | | | |
|---|---|---|---|---|---|---|---|---|
| Anchorwave | 0.9975 | 0.9287 | 0.8842 | 0.8709 | 0.7751 | 0.6897 | 0.7889 | 0.3718 |
| last | 0.9984 | 0.9314 | 0.8897 | 0.8751 | 0.7845 | 0.695 | 0.7958 | 0.3991 |
| last-split | 0.9969 | 0.9289 | 0.8851 | 0.8698 | 0.7799 | 0.6917 | 0.7851 | 0.3885 |
| Anchorwave | 0.9975 | 0.9284 | 0.884 | 0.8709 | 0.776 | 0.6901 | 0.7905 | 0.1966 |
| SVIM-asm | 0.9716 | 0.8911 | 0.8491 | 0.8201 | 0.7173 | 0.6168 | 0.6991 | 0.1803 |
| paftools | 0.8425 | 0.7615 | 0.7169 | 0.6937 | 0.5817 | 0.4834 | 0.5474 | 0.023 |
| Assemblytics | 0.8499 | 0.7904 | 0.7519 | 0.7286 | 0.6569 | 0.5817 | 0.663 | 0.1749 |
| SYRI | 0.9314 | 0.8403 | 0.7909 | 0.7644 | 0.6484 | 0.5512 | 0.5956 | 0.0461 |
| GSalign | 0.0014 | 0.0032 | 0.0034 | 0.0041 | 0.0069 | 0.005 | 0.0095 | 0.0036 |
| MUM&co | 0.8722 | 0.7421 | 0.6849 | 0.6635 | 0.5171 | 0.287 | 0.3978 | 0 |
| ***Precision without normalization for simulated 10,000 insertions in different rice genomes (size ≥ 50 bp)*** | | | | | | | | |
| minimap2 | 0.9767 | 0.9289 | 0.9086 | 0.8672 | 0.7957 | 0.7392 | 0.7884 | 0.4026 |
| MUMmer4 | 0.9996 | 0.9940 | 0.9878 | 0.9833 | 0.9716 | 0.9606 | 0.9749 | 0.8950 |
| Anchorwave | 0.9909 | 0.9680 | 0.9514 | 0.9301 | 0.8888 | 0.8865 | 0.9099 | 0.7461 |
| last | 0.9990 | 0.9796 | 0.9259 | 0.8982 | 0.8445 | 0.9042 | 0.9193 | 0.7821 |
| last-split | 0.9996 | 0.9871 | 0.9721 | 0.9700 | 0.9136 | 0.9401 | 0.9408 | 0.8674 |
| Anchorwave | 0.9879 | 0.9614 | 0.9453 | 0.9201 | 0.8927 | 0.8617 | 0.8902 | 0.5175 |
| SVIM-asm | 0.9605 | 0.8427 | 0.8022 | 0.7472 | 0.6294 | 0.5307 | 0.6182 | 0.2079 |
| paftools | 0.9536 | 0.8761 | 0.8562 | 0.7928 | 0.6988 | 0.6178 | 0.6833 | 0.0939 |
| Assemblytics | 0.8924 | 0.9643 | 0.9529 | 0.9444 | 0.9222 | 0.8914 | 0.9052 | 0.9081 |
| SYRI | 0.9806 | 0.9867 | 0.9793 | 0.9752 | 0.9566 | 0.9435 | 0.9652 | 0.9147 |
| GSalign | 0.0376 | 1.0000 | 1.0000 | 1.0000 | 1.0000 | 1.0000 | 1.0000 | 1.0000 |
| MUM&co | 0.9726 | 0.8689 | 0.8048 | 0.7622 | 0.6190 | 0.5874 | 0.5449 | 0.0000 |
| ***F1 score without normalization for simulated 10,000 insertions in different rice genomes (size ≥ 50 bp)*** | | | | | | | | |
| minimap2 | 0.9807 | 0.9194 | 0.8878 | 0.8570 | 0.7834 | 0.6957 | 0.7675 | 0.3299 |
| MUMmer4 | 0.9970 | 0.9579 | 0.9324 | 0.9210 | 0.8614 | 0.8001 | 0.8655 | 0.4448 |
| Anchorwave | 0.9942 | 0.9479 | 0.9166 | 0.8995 | 0.8281 | 0.7758 | 0.8451 | 0.4963 |
| last | 0.9987 | 0.9549 | 0.9074 | 0.8865 | 0.8134 | 0.7859 | 0.8531 | 0.5285 |
| last-split | 0.9982 | 0.9571 | 0.9266 | 0.9172 | 0.8415 | 0.7970 | 0.8559 | 0.5366 |
| Anchorwave | 0.9927 | 0.9446 | 0.9136 | 0.8948 | 0.8303 | 0.7664 | 0.8374 | 0.2849 |
| SVIM-asm | 0.9660 | 0.8662 | 0.8250 | 0.7820 | 0.6705 | 0.5705 | 0.6562 | 0.1931 |
| paftools | 0.8946 | 0.8148 | 0.7804 | 0.7399 | 0.6349 | 0.5424 | 0.6079 | 0.0370 |
| Assemblytics | 0.8706 | 0.8687 | 0.8405 | 0.8226 | 0.7673 | 0.7040 | 0.7654 | 0.2933 |
| SYRI | 0.9554 | 0.9076 | 0.8751 | 0.8570 | 0.7729 | 0.6959 | 0.7366 | 0.0878 |
| GSalign | 0.0027 | 0.0064 | 0.0068 | 0.0082 | 0.0137 | 0.0100 | 0.0188 | 0.0072 |
| MUM&co | 0.9197 | 0.8005 | 0.7400 | 0.7094 | 0.5635 | 0.3856 | 0.4599 | 0.0000 |

| **Tomato** | **SL4** | **M82** | **ZY65** | **ZY56** | **ZY57** | **ZY60** | **ZY61** | **ZY59** |
|---|---|---|---|---|---|---|---|---|
| ***Recall without normalization for simulated ~13,200 deletions in different tomato genomes (size ≥ 50 bp)*** | | | | | | | | |
| minimap2 | 0.9594 | 0.8840 | 0.9274 | 0.8060 | 0.8077 | 0.4149 | 0.3080 | 0.2597 |
| MUMmer4 | 0.9877 | 0.9090 | 0.9572 | 0.7833 | 0.8075 | 0.4001 | 0.3045 | 0.2483 |
| AnchorWave | 0.9832 | 0.9113 | 0.9540 | 0.7582 | 0.7831 | 0.3429 | 0.2518 | 0.2248 |
| last-split | 0.9935 | 0.9076 | 0.9612 | 0.8177 | 0.8479 | 0.4409 | 0.3299 | 0.2853 |
| SVIM-asm | 0.8780 | 0.8106 | 0.8446 | 0.7431 | 0.7593 | 0.4235 | 0.3149 | 0.2584 |
| SyRI | 0.5277 | 0.4896 | 0.5193 | 0.4775 | 0.4813 | 0.2403 | 0.1615 | 0.0900 |

| | | | | | | | | |
|---|---|---|---|---|---|---|---|---|
| paftools | 0.8638 | 0.7953 | 0.8312 | 0.7292 | 0.7377 | 0.3593 | 0.2426 | 0.1824 |
| Assemblytics | 0.4652 | 0.4161 | 0.4472 | 0.3929 | 0.3905 | 0.2439 | 0.1896 | 0.1454 |
| AnchorWave | 0.9831 | 0.9114 | 0.9548 | 0.7583 | 0.6540 | 0.3436 | 0.2526 | 0.2254 |
| MUM&co | 0.8612 | 0.7052 | 0.7972 | 0.4110 | 0.4328 | 0.0039 | 0.0004 | 0.0001 |
| ***Precision without normalization for simulated ~13,200 deletions in different tomato genomes (size ≥ 50 bp)*** | | | | | | | | |
| minimap2 | 0.9076 | 0.8532 | 0.8865 | 0.8041 | 0.8220 | 0.5206 | 0.4475 | 0.4092 |
| MUMmer4 | 0.9962 | 0.9884 | 0.9950 | 0.9708 | 0.9747 | 0.8101 | 0.7350 | 0.7079 |
| AnchorWave | 0.9785 | 0.8941 | 0.9638 | 0.8929 | 0.9098 | 0.6805 | 0.5494 | 0.5518 |
| last-split | 0.9972 | 0.9820 | 0.9903 | 0.9565 | 0.9663 | 0.8476 | 0.7663 | 0.5646 |
| SVIM-asm | 0.7715 | 0.6842 | 0.5255 | 0.3318 | 0.5873 | 0.0750 | 0.0513 | 0.0396 |
| SyRI | 0.9863 | 0.9642 | 0.9644 | 0.9380 | 0.9438 | 0.8501 | 0.7848 | 0.7428 |
| paftools | 0.8363 | 0.7553 | 0.7911 | 0.5464 | 0.5593 | 0.3564 | 0.2914 | 0.2474 |
| Assemblytics | 0.9321 | 0.9063 | 0.9091 | 0.8620 | 0.8634 | 0.6228 | 0.5508 | 0.4956 |
| AnchorWave | 0.9744 | 0.8717 | 0.9559 | 0.8574 | 0.8716 | 0.5352 | 0.4050 | 0.3919 |
| Mumandco | 0.9778 | 0.9162 | 0.9645 | 0.6634 | 0.7004 | 0.0584 | 0.0230 | 0.0122 |
| ***F1 score without normalization for simulated ~13,200 deletions in different tomato genomes (size ≥ 50 bp)*** | | | | | | | | |
| minimap2 | 0.9328 | 0.8683 | 0.9065 | 0.8051 | 0.8148 | 0.4618 | 0.3649 | 0.3177 |
| MUMmer4 | 0.9919 | 0.9470 | 0.9757 | 0.8670 | 0.8833 | 0.5357 | 0.4306 | 0.3676 |
| AnchorWave | 0.9808 | 0.9026 | 0.9589 | 0.8201 | 0.8417 | 0.4560 | 0.3454 | 0.3195 |
| last-split | 0.9953 | 0.9433 | 0.9755 | 0.8816 | 0.9032 | 0.5801 | 0.4613 | 0.3791 |
| SVIM-asm | 0.8213 | 0.7421 | 0.6479 | 0.4587 | 0.6624 | 0.1275 | 0.0882 | 0.0687 |
| SyRI | 0.6875 | 0.6495 | 0.6751 | 0.6329 | 0.6375 | 0.3747 | 0.2679 | 0.1605 |
| paftools | 0.8498 | 0.7748 | 0.8107 | 0.6247 | 0.6362 | 0.3579 | 0.2648 | 0.2100 |
| Assemblytics | 0.6207 | 0.5703 | 0.5995 | 0.5398 | 0.5377 | 0.3505 | 0.2820 | 0.2249 |
| AnchorWave | 0.9787 | 0.8911 | 0.9554 | 0.8048 | 0.7473 | 0.4185 | 0.3111 | 0.2862 |
| MUM&co | 0.9158 | 0.7969 | 0.8729 | 0.5075 | 0.5350 | 0.0074 | 0.0007 | 0.0002 |
| ***Recall without normalization for simulated ~14,800 insertions in different tomato genomes (size ≥ 50 bp)*** | | | | | | | | |
| minimap2 | 0.9776 | 0.9289 | 0.9514 | 0.8338 | 0.8611 | 0.5593 | 0.4609 | 0.4038 |
| MUMmer4 | 0.9961 | 0.9495 | 0.9747 | 0.8375 | 0.8622 | 0.5542 | 0.4567 | 0.4088 |
| AnchorWave | 0.9956 | 0.9503 | 0.9738 | 0.8141 | 0.8365 | 0.4545 | 0.3505 | 0.3266 |
| last-split | 0.9978 | 0.9468 | 0.9745 | 0.8482 | 0.8826 | 0.5896 | 0.4888 | 0.4396 |
| SVIM-asm | 0.9560 | 0.9088 | 0.9310 | 0.8494 | 0.8672 | 0.5761 | 0.4620 | 0.3958 |
| SyRI | 0.9248 | 0.8560 | 0.8849 | 0.7709 | 0.7803 | 0.4032 | 0.2767 | 0.1672 |
| paftools | 0.7428 | 0.7066 | 0.7257 | 0.6635 | 0.6733 | 0.4077 | 0.3068 | 0.2374 |
| Assemblytics | 0.8815 | 0.8159 | 0.8415 | 0.7393 | 0.7563 | 0.5271 | 0.4348 | 0.3650 |
| anchorwave | 0.9959 | 0.9507 | 0.9745 | 0.8151 | 0.7045 | 0.4578 | 0.3533 | 0.3310 |
| MUM&co | 0.8633 | 0.7472 | 0.8228 | 0.5045 | 0.5184 | 0.0084 | 0.0010 | 0.0005 |
| ***Precision without normalization for simulated ~14,800 insertions in different tomato genomes (size ≥ 50 bp)*** | | | | | | | | |
| minimap2 | 0.9780 | 0.9129 | 0.9584 | 0.8748 | 0.9009 | 0.6438 | 0.5562 | 0.5238 |
| Mummer4 | 1.0000 | 0.9868 | 0.9952 | 0.9795 | 0.9862 | 0.8742 | 0.8431 | 0.8433 |
| AnchorWave | 0.9881 | 0.9185 | 0.9697 | 0.9117 | 0.9296 | 0.7545 | 0.6326 | 0.6698 |
| last-split | 0.9998 | 0.9858 | 0.9897 | 0.9628 | 0.9728 | 0.9022 | 0.8611 | 0.7793 |
| SVIM-asm | 0.8623 | 0.7880 | 0.7668 | 0.7213 | 0.6937 | 0.3510 | 0.2748 | 0.2524 |
| SyRI | 0.9921 | 0.9817 | 0.9818 | 0.9675 | 0.9642 | 0.9225 | 0.8561 | 0.8704 |
| paftools | 0.9232 | 0.8440 | 0.8885 | 0.7809 | 0.6226 | 0.4349 | 0.3609 | 0.3046 |
| Assemblytics | 0.9577 | 0.9430 | 0.9461 | 0.9172 | 0.9211 | 0.8078 | 0.7341 | 0.7296 |

| | | | | | | | | |
|---|---|---|---|---|---|---|---|---|
| AnchorWave | 0.9860 | 0.9045 | 0.9655 | 0.8788 | 0.8977 | 0.6214 | 0.4984 | 0.5122 |
| MUM&co | 0.9727 | 0.8756 | 0.9243 | 0.6659 | 0.6943 | 0.1249 | 0.0526 | 0.0825 |
| ***F1 score without normalization for simulated ~14,800 insertions in different tomato genomes (size ≥ 50 bp)*** | | | | | | | | |
| minimap2 | 0.9778 | 0.9208 | 0.9548 | 0.8538 | 0.8806 | 0.5986 | 0.5041 | 0.4561 |
| MUMmer4 | 0.9981 | 0.9678 | 0.9849 | 0.9029 | 0.9200 | 0.6783 | 0.5925 | 0.5507 |
| AnchorWave | 0.9919 | 0.9341 | 0.9718 | 0.8601 | 0.8806 | 0.5673 | 0.4511 | 0.4391 |
| last-split | 0.9988 | 0.9659 | 0.9821 | 0.9019 | 0.9255 | 0.7132 | 0.6236 | 0.5621 |
| SVIM-asm | 0.9067 | 0.8441 | 0.8410 | 0.7801 | 0.7708 | 0.4362 | 0.3447 | 0.3082 |
| SyRI | 0.9573 | 0.9146 | 0.9309 | 0.8581 | 0.8625 | 0.5612 | 0.4182 | 0.2805 |
| paftools | 0.8233 | 0.7692 | 0.7989 | 0.7174 | 0.6470 | 0.4209 | 0.3317 | 0.2668 |
| Assemblytics | 0.9180 | 0.8749 | 0.8908 | 0.8187 | 0.8306 | 0.6379 | 0.5461 | 0.4866 |
| AnchorWave | 0.9910 | 0.9270 | 0.9700 | 0.8458 | 0.7895 | 0.5272 | 0.4135 | 0.4021 |
| MUM&co | 0.9148 | 0.8063 | 0.8706 | 0.5741 | 0.5936 | 0.0157 | 0.0020 | 0.0011 |

**Supplementary Table 7: Commands and parameters for testing different methods to call SVs in plant genomes.**

| Methods | Aligners | Parameters | Reference |
|---|---|---|---|
| Assemblytics | MUMmer4.0 | Assemblytics REFvsQRY.mm.delta REF **100**/500/1000/10000 1 40000<br>Assemblytics REFvsQRY.max.delta REF **100**/500/1000/10000 1 40000 | 18 |
| SyRI | minimap2 | minimap2 -ax asm5 --eqx -t 4 $REF $QRY > REF.QRY.sam<br>syri -c REFvsQRY.sam -r $REF -q $QRY -k -F S | 19 |
| | MUMmer4.0 | nucmer -t 4 --prefix=REF.QRY.mm $REF $QRY<br>show-coords -THrd REF.QRY.mm.delta > REF.QRY.mm.coords<br>syri -d REFvsQRY.mm.delta -c REFvsQRY4.mm.coords -r $REF -q $QRY | |
| SVIM-asm | minimap2 | minimap2 -a -x asm5 --cs -r2k -t 4 $REF $QRY > REF.QRY.sam<br>samtools sort -m4G -@4 -o REF.QRY.sorted.bam REF.QRY.sam<br>samtools index REF.QRY.sorted.bam<br>svim-asm haploid ./SV REF.QRY.sorted.bam $REF | 20 |
| paftools | minimap2 | minimap2 -c --cs=long $REF $QRY > REFvsQRY.def.paf | 15 |
| GSAlign | GSAlign | GSAlign -r $REF -q $QRY -o REFvsQRY | 17 |
| MUM&Co | MUMmer4.0 | mumandco_v3.8.sh -r $REF -q $QRY -g genome.size -o REFvsQRY -t 4 | 21 |
| AnchorWave | AnchorWave | anchorwave genoAli -i REF.gff3 -as cds.fa -r $REF -a cds.sam -ar REF.sam -s $QRY -n anchors -o anchorwave.maf -f anchorwave.f.maf -v SV.vcf -m 0 -t 4 | 16 |
| SVGAP | minimap2/last | see manual | This study |

#### Supplementary Table 8: Pairwise comparison of SV calls between different methods.

##### SVs detected between two rice genome assemblies: MH63 and ZS97.

| | AnchorWave | Assemblytics | GSAlign | paftools | MUM&co | SVGAP last | SVGAP minimap2 | SVIM-asm | SyRI minimap2 | SyRI MUMmer4 |
|---|---|---|---|---|---|---|---|---|---|---|
| AnchorWave | **24,960** | 10,261 | 1,399 | 12,741 | 6,287 | 16,207 | 14,985 | 9,854 | 4,482 | 2,830 |
| Assemblytics | 10,249 | **11,993** | 985 | 8,566 | 4,708 | 9,689 | 9,832 | 7,517 | 3,538 | 3,159 |
| GSAlign | 1,394 | 985 | **2,557** | 1,328 | 152 | 1,383 | 1,497 | 1,208 | 805 | 724 |
| paftools | 12,741 | 8,576 | 1,334 | **19,889** | 5,541 | 12,697 | 16,952 | 10,912 | 5,532 | 2,528 |
| MUM&co | 6,280 | 4,711 | 152 | 5,537 | **19,514** | 5,883 | 5,893 | 5,258 | 2,253 | 762 |
| SVGAPS/ last | 16,213 | 9,700 | 1,388 | 12,700 | 5,888 | **21,241** | 15,008 | 10,198 | 4,820 | 2,735 |
| SVGAPS/ minimap2 | 14,988 | 9,843 | 1,503 | 16,992 | 5,897 | 15,008 | **20,503** | 11,650 | 5,804 | 2,812 |
| SVIM-asm | 9,849 | 7,525 | 1,211 | 10,905 | 5,257 | 10,189 | 11,640 | **13,402** | 6,258 | 2,377 |
| SyRI/ minimap2 | 4,479 | 3,544 | 807 | 5,531 | 2,252 | 4,817 | 5,802 | 6,269 | **6,571** | 1,510 |
| SyRI/ MUMmer4 | 2,831 | 3,167 | 727 | 2,530 | 762 | 2,736 | 2,814 | 2,381 | 1,511 | **3,606** |

##### SVs detected between two tomato genome assemblies: SL4 and M82.

| | AnchorWave | Assemblytics | GSAlign | paftools | MUM&co | SVGAP last | SVGAP minimap2 | SVIM-asm | SyRI minimap2 | SyRI MUMmer4 |
|---|---|---|---|---|---|---|---|---|---|---|
| AnchorWave | **10,494** | 3,231 | 703 | 1,985 | 5,850 | 5,600 | 6,347 | 4,601 | 3,455 | 2,103 |
| Assemblytics | 3,229 | **4,146** | 526 | 994 | 3,284 | 2,958 | 3,304 | 3,093 | 2,413 | 2,193 |
| GSAlign | 702 | 527 | **1,152** | 41 | 809 | 625 | 797 | 765 | 701 | 476 |
| MUM&co | 1,986 | 995 | 41 | **4,695** | 1,967 | 2,131 | 2,173 | 1,501 | 1,280 | 506 |
| paftools | 5,842 | 3,285 | 810 | 1,966 | **13,170** | 5,684 | 9,772 | 5,914 | 4,627 | 2,113 |
| SVGAP/ last | 5,604 | 2,961 | 626 | 2,131 | 5,691 | **8,619** | 6,260 | 4,614 | 3,499 | 1,795 |
| SVGAP/ minimap2 | 6,347 | 3,306 | 798 | 2,173 | 9,778 | 6,260 | **11,449** | 5,883 | 4,594 | 2,105 |
| SVIM-asm | 4,598 | 3,096 | 766 | 1,499 | 5,913 | 4,611 | 5,879 | **7,346** | 5,246 | 2,045 |
| SyRI/ minimap2 | 3,453 | 2,414 | 702 | 1,279 | 4,628 | 3,497 | 4,592 | 5,250 | **5,435** | 1,761 |
| SyRI/ MUMmer4 | 2,103 | 2,196 | 476 | 506 | 2,116 | 1,795 | 2,105 | 2,045 | 1,762 | **2,680** |

**Supplementary Table 9: Recall of the merging step in SVGAP evaluated using population-scale simulation data based on *Oryza* and *Solanum* phylogeny.**

| **Benchmarking in rice** | | **SVs retrieved from the merging step in SVGAP** | | | | | | | |
|---|---|---|---|---|---|---|---|---|---|
| SV types | # of simulated SVs | Nipp | MH63 | R498 | 9311 | ZS97 | Oruf | CJ14 | Opun |
| Deletions (< 50 bp) | 1,266 | 1,265 | 894 | 900 | 871 | 906 | 839 | 802 | 208 |
| Deletions (≥ 50 bp) | 1,711 | 1,707 | 1,265 | 1,270 | 1,196 | 1,245 | 1,146 | 1,155 | 377 |
| Insertions (< 50 bp) | 864 | 798 | 620 | 618 | 596 | 619 | 559 | 546 | 163 |
| Insertions (≥ 50 bp) | 1,368 | 1,367 | 1,013 | 1,019 | 975 | 1,012 | 932 | 918 | 390 |
| **Benchmarking in tomato** | | **SVs retrieved from the merging step in SVGAP** | | | | | | | |
| | # of simulated SVs | SL4 | M82 | ZY65 | ZY56 | ZY57 | ZY60 | ZY61 | ZY59 |
| Deletions (≥ 50 bp) | 3,017 | 2,978 | 2,815 | 2,774 | 2,755 | 2,566 | 1,748 | 1,406 | 1,034 |
| Insertions (≥ 50 bp) | 3,382 | 3,356 | 3,245 | 3,149 | 3,036 | 2,906 | 1,695 | 1,340 | 1,106 |

**Supplementary Table 10: Performance of the genotyping step in SVGAP evaluated using population-scale simulation data based on *Oryza* and *Solanum* phylogeny.**

**Accuracy, error, and missing rate for benchmarking analysis for insertions using *Oryza* phylogeny-based data.**

| | Size | Nipp | MH63 | R498 | 9311 | ZS97 | Oruf | CJ14 | Opun | Size | Nipp | MH63 | R498 | 9311 | ZS97 | Oruf | CJ14 | Opun |
|---|---|---|---|---|---|---|---|---|---|---|---|---|---|---|---|---|---|---|
| **# of retrieved calls** | | **798** | **620** | **618** | **596** | **619** | **559** | **546** | **163** | **1,367** | **1,013** | **1,019** | **975** | **1,012** | **932** | **918** | **390** | **798** |
| **Accuracy** | | | | | | | | | | | | | | | | | | |
| SIM0 | <50 bp | 730 | 611 | 614 | 592 | 613 | 557 | 542 | 160 | ≥ 50 bp | 1,348 | 1,012 | 1,018 | 974 | 1,010 | 929 | 917 | 386 |
| SIM1 | <50 bp | 783 | 580 | 586 | 559 | 583 | 515 | 506 | 132 | ≥ 50 bp | 1,341 | 967 | 982 | 935 | 970 | 887 | 869 | 316 |
| SIM2 | <50 bp | 783 | 577 | 579 | 560 | 583 | 518 | 499 | 125 | ≥ 50 bp | 1,339 | 975 | 985 | 936 | 973 | 879 | 859 | 305 |
| SIM3 | <50 bp | 778 | 589 | 588 | 570 | 592 | 526 | 509 | 134 | ≥ 50 bp | 1,338 | 970 | 985 | 935 | 979 | 889 | 864 | 309 |
| SIM4 | <50 bp | 784 | 589 | 596 | 571 | 594 | 530 | 507 | 127 | ≥ 50 bp | 1,347 | 974 | 983 | 944 | 977 | 872 | 862 | 317 |
| SIM5 | <50 bp | 778 | 589 | 593 | 567 | 591 | 528 | 504 | 132 | ≥ 50 bp | 1,338 | 973 | 982 | 937 | 971 | 880 | 872 | 307 |
| SIM6 | <50 bp | 782 | 579 | 583 | 562 | 588 | 520 | 503 | 116 | ≥ 50 bp | 1,342 | 980 | 988 | 948 | 974 | 891 | 869 | 319 |
| SIM7 | <50 bp | 779 | 586 | 591 | 565 | 589 | 528 | 514 | 130 | ≥ 50 bp | 1,342 | 974 | 982 | 936 | 968 | 888 | 863 | 316 |
| SIM8 | <50 bp | 783 | 587 | 591 | 564 | 587 | 516 | 500 | 115 | ≥ 50 bp | 1,329 | 979 | 987 | 938 | 974 | 886 | 866 | 317 |
| SIM9 | <50 bp | 781 | 589 | 594 | 569 | 589 | 516 | 504 | 131 | ≥ 50 bp | 1,346 | 972 | 986 | 937 | 977 | 885 | 875 | 311 |
| SIM10 | <50 bp | 783 | 582 | 580 | 560 | 585 | 519 | 505 | 139 | ≥ 50 bp | 1,342 | 968 | 979 | 934 | 966 | 876 | 877 | 320 |
| SIM11 | <50 bp | 786 | 584 | 586 | 562 | 581 | 525 | 503 | 126 | ≥ 50 bp | 1,335 | 973 | 985 | 940 | 967 | 881 | 865 | 309 |
| SIM12 | <50 bp | 776 | 582 | 586 | 561 | 582 | 523 | 511 | 132 | ≥ 50 bp | 1,349 | 973 | 981 | 929 | 969 | 882 | 861 | 321 |
| SIM13 | <50 bp | 778 | 585 | 590 | 570 | 591 | 524 | 506 | 125 | ≥ 50 bp | 1,349 | 979 | 992 | 936 | 972 | 890 | 860 | 322 |
| SIM14 | <50 bp | 783 | 592 | 590 | 567 | 592 | 528 | 506 | 130 | ≥ 50 bp | 1,338 | 973 | 983 | 932 | 975 | 887 | 872 | 314 |
| SIM15 | <50 bp | 772 | 584 | 588 | 568 | 583 | 522 | 506 | 131 | ≥ 50 bp | 1,353 | 972 | 982 | 930 | 967 | 881 | 874 | 313 |
| SIM16 | <50 bp | 780 | 584 | 587 | 566 | 585 | 524 | 502 | 130 | ≥ 50 bp | 1,342 | 968 | 983 | 935 | 963 | 878 | 863 | 314 |
| SIM17 | <50 bp | 782 | 585 | 591 | 565 | 589 | 524 | 502 | 133 | ≥ 50 bp | 1,343 | 973 | 987 | 936 | 971 | 891 | 873 | 311 |
| SIM18 | <50 bp | 778 | 585 | 589 | 571 | 594 | 514 | 504 | 128 | ≥ 50 bp | 1,347 | 977 | 987 | 936 | 970 | 877 | 873 | 320 |
| SIM19 | <50 bp | 782 | 586 | 586 | 567 | 590 | 523 | 502 | 127 | ≥ 50 bp | 1,344 | 970 | 980 | 935 | 969 | 886 | 868 | 303 |
| SIM20 | <50 bp | 780 | 585 | 589 | 566 | 581 | 518 | 503 | 126 | ≥ 50 bp | 1,345 | 972 | 989 | 938 | 973 | 884 | 867 | 312 |
| **Average** | | 778 | 586 | 589 | 567 | 589 | 524 | 507 | 130 | | 1,343 | 975 | 986 | 938 | 973 | 886 | 870 | 317 |
| **Accuracy** | | 97.5% | 94.5% | 95.4% | 95.1% | 95.1% | 93.7% | 92.8% | 79.7% | | 98.2% | 96.2% | 96.8% | 96.2% | 96.2% | 95.0% | 94.8% | 81.3% |

| | | | | | | | | | | | | | | | | | | |
|---|---|---|---|---|---|---|---|---|---|---|---|---|---|---|---|---|---|---|
| **Error rate** | | | | | | | | | | | | | | | | | | |
| SIM0 | <50 bp | 35 | 6 | 2 | 1 | 2 | 0 | 0 | 0 | ≥ 50 bp | 8 | 0 | 0 | 0 | 0 | 0 | 0 | 1 |
| SIM1 | <50 bp | 9 | 18 | 13 | 14 | 14 | 22 | 19 | 9 | ≥ 50 bp | 8 | 28 | 20 | 18 | 20 | 24 | 33 | 22 |
| SIM2 | <50 bp | 9 | 24 | 21 | 13 | 15 | 18 | 18 | 10 | ≥ 50 bp | 8 | 24 | 17 | 22 | 22 | 27 | 36 | 25 |
| SIM3 | <50 bp | 15 | 15 | 14 | 7 | 9 | 15 | 12 | 4 | ≥ 50 bp | 10 | 27 | 20 | 24 | 20 | 27 | 33 | 23 |
| SIM4 | <50 bp | 3 | 18 | 11 | 13 | 13 | 13 | 15 | 7 | ≥ 50 bp | 7 | 25 | 22 | 14 | 21 | 39 | 35 | 20 |
| SIM5 | <50 bp | 14 | 20 | 14 | 13 | 15 | 17 | 16 | 7 | ≥ 50 bp | 7 | 23 | 22 | 20 | 22 | 31 | 32 | 21 |
| SIM6 | <50 bp | 9 | 21 | 17 | 15 | 14 | 20 | 17 | 2 | ≥ 50 bp | 7 | 20 | 19 | 17 | 25 | 26 | 35 | 4 |
| SIM7 | <50 bp | 11 | 20 | 14 | 14 | 16 | 17 | 15 | 8 | ≥ 50 bp | 10 | 23 | 17 | 15 | 25 | 23 | 34 | 21 |
| SIM8 | <50 bp | 8 | 17 | 13 | 15 | 15 | 21 | 20 | 1 | ≥ 50 bp | 12 | 19 | 17 | 21 | 21 | 28 | 35 | 5 |
| SIM9 | <50 bp | 12 | 16 | 15 | 14 | 14 | 21 | 19 | 6 | ≥ 50 bp | 7 | 21 | 12 | 18 | 18 | 24 | 29 | 22 |
| SIM10 | <50 bp | 8 | 21 | 19 | 18 | 16 | 19 | 19 | 6 | ≥ 50 bp | 7 | 29 | 21 | 23 | 26 | 38 | 25 | 19 |
| SIM11 | <50 bp | 5 | 19 | 15 | 16 | 18 | 18 | 17 | 6 | ≥ 50 bp | 12 | 26 | 19 | 21 | 26 | 30 | 33 | 18 |
| SIM12 | <50 bp | 11 | 19 | 16 | 16 | 17 | 18 | 13 | 6 | ≥ 50 bp | 5 | 23 | 18 | 22 | 22 | 28 | 31 | 15 |
| SIM13 | <50 bp | 9 | 19 | 16 | 13 | 14 | 17 | 21 | 6 | ≥ 50 bp | 7 | 22 | 14 | 21 | 20 | 25 | 37 | 16 |
| SIM14 | <50 bp | 10 | 14 | 16 | 15 | 13 | 13 | 20 | 9 | ≥ 50 bp | 10 | 29 | 21 | 25 | 24 | 28 | 31 | 17 |
| SIM15 | <50 bp | 15 | 22 | 15 | 11 | 17 | 20 | 17 | 10 | ≥ 50 bp | 3 | 28 | 23 | 28 | 27 | 30 | 30 | 23 |
| SIM16 | <50 bp | 9 | 19 | 17 | 11 | 15 | 15 | 18 | 8 | ≥ 50 bp | 10 | 27 | 21 | 22 | 26 | 35 | 37 | 25 |
| SIM17 | <50 bp | 11 | 17 | 13 | 14 | 13 | 15 | 20 | 7 | ≥ 50 bp | 5 | 24 | 17 | 22 | 23 | 23 | 27 | 22 |
| SIM18 | <50 bp | 7 | 19 | 16 | 11 | 11 | 22 | 20 | 7 | ≥ 50 bp | 11 | 20 | 15 | 16 | 22 | 34 | 25 | 19 |
| SIM19 | <50 bp | 10 | 18 | 17 | 12 | 11 | 17 | 17 | 8 | ≥ 50 bp | 7 | 23 | 19 | 17 | 24 | 29 | 28 | 27 |
| SIM20 | <50 bp | 10 | 19 | 14 | 14 | 18 | 22 | 19 | 10 | ≥ 50 bp | 7 | 27 | 16 | 23 | 27 | 26 | 33 | 24 |
| **Average** | | 11 | 18 | 15 | 13 | 14 | 17 | 17 | 7 | | 8 | 23 | 18 | 19 | 22 | 27 | 30 | 19 |
| **Error rate** | | 1.37% | 2.93% | 2.37% | 2.16% | 2.23% | 3.07% | 3.07% | 4.00% | | 0.59% | 2.29% | 1.73% | 2.00% | 2.17% | 2.94% | 3.31% | 4.75% |
| **Missing rate** | | | | | | | | | | | | | | | | | | |
| SIM0 | <50 bp | 33 | 3 | 2 | 3 | 4 | 2 | 4 | 3 | ≥ 50 bp | 11 | 1 | 1 | 1 | 2 | 3 | 1 | 3 |
| SIM1 | <50 bp | 6 | 22 | 19 | 23 | 22 | 22 | 21 | 22 | ≥ 50 bp | 18 | 18 | 17 | 22 | 22 | 21 | 16 | 52 |
| SIM2 | <50 bp | 6 | 19 | 18 | 23 | 21 | 23 | 29 | 28 | ≥ 50 bp | 20 | 14 | 17 | 17 | 17 | 26 | 23 | 60 |
| SIM3 | <50 bp | 5 | 16 | 16 | 19 | 18 | 18 | 25 | 25 | ≥ 50 bp | 19 | 16 | 14 | 16 | 13 | 16 | 21 | 58 |
| SIM4 | <50 bp | 11 | 13 | 11 | 12 | 12 | 16 | 24 | 29 | ≥ 50 bp | 13 | 14 | 14 | 17 | 14 | 21 | 21 | 53 |
| SIM5 | <50 bp | 6 | 11 | 11 | 16 | 13 | 14 | 26 | 24 | ≥ 50 bp | 22 | 17 | 15 | 18 | 19 | 21 | 14 | 62 |

| | | | | | | | | | | | | | | | | | | |
|---|---|---|---|---|---|---|---|---|---|---|---|---|---|---|---|---|---|---|
| SIM6 | <50 bp | 7 | 20 | 18 | 19 | 17 | 19 | 26 | 45 | ≥ 50 bp | 18 | 13 | 12 | 10 | 13 | 15 | 14 | 67 |
| SIM7 | <50 bp | 8 | 14 | 13 | 17 | 14 | 14 | 17 | 25 | ≥ 50 bp | 15 | 16 | 20 | 24 | 19 | 21 | 21 | 53 |
| SIM8 | <50 bp | 7 | 16 | 14 | 17 | 17 | 22 | 26 | 47 | ≥ 50 bp | 26 | 15 | 15 | 16 | 17 | 18 | 17 | 68 |
| SIM9 | <50 bp | 5 | 15 | 9 | 13 | 16 | 22 | 23 | 26 | ≥ 50 bp | 14 | 20 | 21 | 20 | 17 | 23 | 14 | 57 |
| SIM10 | <50 bp | 7 | 17 | 19 | 18 | 18 | 21 | 22 | 18 | ≥ 50 bp | 18 | 16 | 19 | 18 | 20 | 18 | 16 | 51 |
| SIM11 | <50 bp | 7 | 17 | 17 | 18 | 20 | 16 | 26 | 31 | ≥ 50 bp | 20 | 14 | 15 | 14 | 19 | 21 | 20 | 63 |
| SIM12 | <50 bp | 11 | 19 | 16 | 19 | 20 | 18 | 22 | 25 | ≥ 50 bp | 13 | 17 | 20 | 24 | 21 | 22 | 26 | 54 |
| SIM13 | <50 bp | 11 | 16 | 12 | 13 | 14 | 18 | 19 | 32 | ≥ 50 bp | 11 | 12 | 13 | 18 | 20 | 17 | 21 | 52 |
| SIM14 | <50 bp | 5 | 14 | 12 | 14 | 14 | 18 | 20 | 24 | ≥ 50 bp | 19 | 11 | 15 | 18 | 13 | 17 | 15 | 59 |
| SIM15 | <50 bp | 11 | 14 | 15 | 17 | 19 | 17 | 23 | 22 | ≥ 50 bp | 11 | 13 | 14 | 17 | 18 | 21 | 14 | 54 |
| SIM16 | <50 bp | 9 | 17 | 14 | 19 | 19 | 20 | 26 | 25 | ≥ 50 bp | 15 | 18 | 15 | 18 | 23 | 19 | 18 | 51 |
| SIM17 | <50 bp | 5 | 18 | 14 | 17 | 17 | 20 | 24 | 23 | ≥ 50 bp | 19 | 16 | 15 | 17 | 18 | 18 | 18 | 57 |
| SIM18 | <50 bp | 13 | 16 | 13 | 14 | 14 | 23 | 22 | 28 | ≥ 50 bp | 9 | 16 | 17 | 23 | 20 | 21 | 20 | 51 |
| SIM19 | <50 bp | 6 | 16 | 15 | 17 | 18 | 19 | 27 | 28 | ≥ 50 bp | 16 | 20 | 20 | 23 | 19 | 17 | 22 | 60 |
| SIM20 | <50 bp | 8 | 16 | 15 | 16 | 20 | 19 | 24 | 27 | ≥ 50 bp | 15 | 14 | 14 | 14 | 12 | 22 | 18 | 54 |
| Average | | 9 | 16 | 14 | 16 | 17 | 18 | 23 | 27 | | 16 | 15 | 15 | 17 | 17 | 19 | 18 | 54 |
| Missing rate | | 1.12% | 2.53% | 2.26% | 2.75% | 2.67% | 3.25% | 4.15% | 16.25% | | 1.19% | 1.46% | 1.51% | 1.78% | 1.68% | 2.03% | 1.92% | 13.91% |

**Accuracy, error, and missing rate for benchmarking analysis for deletions using *Oryza* phylogeny-based data.**

| # of retrieved calls | | **1,265** | **894** | **900** | **871** | **906** | **839** | **802** | **208** | | **1,707** | **1,265** | **1,270** | **1,196** | **1,245** | **1,146** | **1,155** | **377** |
|---|---|---|---|---|---|---|---|---|---|---|---|---|---|---|---|---|---|---|
| **Accuracy** | | **Nipp** | **MH63** | **R498** | **9311** | **ZS97** | **Oruf** | **CJ14** | **Opun** | | **Nipp** | **MH63** | **R498** | **9311** | **ZS97** | **Oruf** | **CJ14** | **Opun** |
| SIM0 | <50 bp | 1,182 | 828 | 832 | 803 | 845 | 768 | 733 | 139 | ≥ 50 bp | 1,690 | 1,264 | 1,270 | 1,195 | 1,243 | 1,143 | 1,153 | 376 |
| SIM1 | <50 bp | 1,242 | 825 | 831 | 804 | 847 | 762 | 732 | 146 | ≥ 50 bp | 1,693 | 1,248 | 1,257 | 1,179 | 1,228 | 1,125 | 1,137 | 329 |
| SIM2 | <50 bp | 1,251 | 838 | 841 | 812 | 852 | 783 | 737 | 138 | ≥ 50 bp | 1,700 | 1,246 | 1,252 | 1,175 | 1,223 | 1,127 | 1,133 | 331 |
| SIM3 | <50 bp | 1,244 | 836 | 841 | 810 | 851 | 769 | 738 | 151 | ≥ 50 bp | 1,689 | 1,247 | 1,257 | 1,180 | 1,225 | 1,121 | 1,135 | 340 |
| SIM4 | <50 bp | 1,228 | 828 | 834 | 798 | 841 | 764 | 734 | 145 | ≥ 50 bp | 1,689 | 1,244 | 1,252 | 1,176 | 1,223 | 1,125 | 1,129 | 335 |
| SIM5 | <50 bp | 1,242 | 827 | 834 | 799 | 839 | 755 | 733 | 143 | ≥ 50 bp | 1,691 | 1,245 | 1,252 | 1,172 | 1,219 | 1,117 | 1,134 | 336 |
| SIM6 | <50 bp | 1,243 | 839 | 841 | 813 | 851 | 774 | 735 | 144 | ≥ 50 bp | 1,687 | 1,248 | 1,255 | 1,180 | 1,223 | 1,120 | 1,132 | 338 |
| SIM7 | <50 bp | 1,240 | 839 | 839 | 809 | 848 | 773 | 739 | 148 | ≥ 50 bp | 1,686 | 1,246 | 1,254 | 1,173 | 1,220 | 1,121 | 1,132 | 342 |
| SIM8 | <50 bp | 1,241 | 838 | 840 | 810 | 844 | 770 | 733 | 147 | ≥ 50 bp | 1,685 | 1,246 | 1,255 | 1,184 | 1,227 | 1,125 | 1,142 | 342 |
| SIM9 | <50 bp | 1,237 | 838 | 845 | 806 | 849 | 760 | 728 | 136 | ≥ 50 bp | 1,691 | 1,244 | 1,249 | 1,176 | 1,215 | 1,112 | 1,132 | 336 |

| | | | | | | | | | | | | | | | | | | |
|---|---|---|---|---|---|---|---|---|---|---|---|---|---|---|---|---|---|---|
| SIM10 | <50 bp | 1,247 | 825 | 830 | 802 | 842 | 766 | 724 | 142 | ≥ 50 bp | 1,684 | 1,243 | 1,253 | 1,174 | 1,225 | 1,121 | 1,136 | 340 |
| SIM11 | <50 bp | 1,248 | 838 | 834 | 814 | 851 | 765 | 730 | 147 | ≥ 50 bp | 1,691 | 1,244 | 1,250 | 1,167 | 1,214 | 1,124 | 1,132 | 331 |
| SIM12 | <50 bp | 1,239 | 835 | 836 | 806 | 845 | 769 | 738 | 148 | ≥ 50 bp | 1,690 | 1,250 | 1,255 | 1,178 | 1,224 | 1,123 | 1,139 | 337 |
| SIM13 | <50 bp | 1,241 | 833 | 838 | 810 | 849 | 763 | 736 | 139 | ≥ 50 bp | 1,689 | 1,249 | 1,258 | 1,179 | 1,227 | 1,124 | 1,142 | 336 |
| SIM14 | <50 bp | 1,245 | 833 | 838 | 804 | 847 | 768 | 737 | 152 | ≥ 50 bp | 1,693 | 1,246 | 1,254 | 1,172 | 1,220 | 1,122 | 1,135 | 338 |
| SIM15 | <50 bp | 1,244 | 833 | 835 | 807 | 851 | 770 | 730 | 146 | ≥ 50 bp | 1,696 | 1,247 | 1,253 | 1,180 | 1,225 | 1,124 | 1,132 | 339 |
| SIM16 | <50 bp | 1,243 | 829 | 829 | 799 | 838 | 774 | 734 | 144 | ≥ 50 bp | 1,693 | 1,240 | 1,252 | 1,177 | 1,221 | 1,119 | 1,131 | 344 |
| SIM17 | <50 bp | 1,238 | 833 | 846 | 806 | 846 | 771 | 735 | 141 | ≥ 50 bp | 1,684 | 1,250 | 1,260 | 1,180 | 1,229 | 1,125 | 1,135 | 336 |
| SIM18 | <50 bp | 1,242 | 847 | 856 | 817 | 853 | 771 | 735 | 143 | ≥ 50 bp | 1,689 | 1,239 | 1,250 | 1,173 | 1,212 | 1,116 | 1,128 | 336 |
| SIM19 | <50 bp | 1,239 | 829 | 839 | 802 | 843 | 770 | 731 | 146 | ≥ 50 bp | 1,694 | 1,246 | 1,256 | 1,177 | 1,223 | 1,124 | 1,129 | 334 |
| SIM20 | <50 bp | 1,241 | 831 | 838 | 803 | 847 | 767 | 731 | 151 | ≥ 50 bp | 1,685 | 1,246 | 1,253 | 1,176 | 1,221 | 1,123 | 1,133 | 331 |
| Average | | 1,239 | 833 | 838 | 806 | 847 | 768 | 733 | 145 | | 1,690 | 1,247 | 1,255 | 1,177 | 1,223 | 1,123 | 1,135 | 338 |
| Accuracy | | 97.94% | 93.22% | 93.11% | 92.58% | 93.45% | 91.56% | 91.46% | 69.51% | | 99.00% | 98.54% | 98.79% | 98.44% | 98.25% | 97.98% | 98.25% | 89.77% |
| **Error rate** | | | | | | | | | | | | | | | | | | |
| SIM0 | <50 bp | 1 | 0 | 0 | 0 | 0 | 0 | 0 | 0 | ≥ 50 bp | 7 | 0 | 0 | 1 | 1 | 0 | 0 | 0 |
| SIM1 | <50 bp | 1 | 2 | 2 | 1 | 1 | 2 | 2 | 0 | ≥ 50 bp | 8 | 9 | 7 | 6 | 8 | 12 | 9 | 7 |
| SIM2 | <50 bp | 0 | 2 | 1 | 1 | 1 | 1 | 1 | 0 | ≥ 50 bp | 3 | 9 | 10 | 8 | 11 | 10 | 13 | 15 |
| SIM3 | <50 bp | 0 | 2 | 1 | 0 | 0 | 1 | 1 | 0 | ≥ 50 bp | 11 | 10 | 8 | 8 | 10 | 16 | 9 | 6 |
| SIM4 | <50 bp | 1 | 2 | 1 | 1 | 0 | 1 | 1 | 0 | ≥ 50 bp | 11 | 11 | 10 | 7 | 10 | 14 | 16 | 9 |
| SIM5 | <50 bp | 2 | 2 | 1 | 2 | 1 | 1 | 1 | 0 | ≥ 50 bp | 11 | 9 | 10 | 8 | 14 | 17 | 13 | 9 |
| SIM6 | <50 bp | 0 | 1 | 1 | 0 | 0 | 1 | 1 | 0 | ≥ 50 bp | 12 | 12 | 10 | 9 | 14 | 16 | 14 | 11 |
| SIM7 | <50 bp | 0 | 2 | 1 | 1 | 1 | 1 | 1 | 0 | ≥ 50 bp | 10 | 9 | 8 | 7 | 12 | 10 | 11 | 8 |
| SIM8 | <50 bp | 0 | 0 | 0 | 0 | 0 | 0 | 0 | 0 | ≥ 50 bp | 14 | 8 | 8 | 5 | 8 | 13 | 6 | 8 |
| SIM9 | <50 bp | 0 | 1 | 1 | 0 | 0 | 2 | 1 | 0 | ≥ 50 bp | 8 | 11 | 12 | 13 | 17 | 21 | 14 | 12 |
| SIM10 | <50 bp | 0 | 1 | 2 | 2 | 2 | 2 | 2 | 0 | ≥ 50 bp | 14 | 13 | 10 | 10 | 11 | 16 | 12 | 5 |
| SIM11 | <50 bp | 0 | 3 | 2 | 1 | 1 | 2 | 2 | 0 | ≥ 50 bp | 10 | 14 | 12 | 15 | 19 | 13 | 13 | 8 |
| SIM12 | <50 bp | 0 | 1 | 0 | 0 | 0 | 0 | 0 | 0 | ≥ 50 bp | 12 | 11 | 11 | 9 | 12 | 14 | 12 | 6 |
| SIM13 | <50 bp | 0 | 2 | 2 | 1 | 2 | 2 | 2 | 0 | ≥ 50 bp | 13 | 10 | 9 | 10 | 14 | 14 | 9 | 14 |
| SIM14 | <50 bp | 1 | 4 | 2 | 2 | 1 | 2 | 2 | 0 | ≥ 50 bp | 8 | 12 | 11 | 13 | 16 | 15 | 10 | 8 |
| SIM15 | <50 bp | 1 | 2 | 1 | 1 | 1 | 2 | 1 | 0 | ≥ 50 bp | 5 | 14 | 13 | 9 | 13 | 16 | 12 | 5 |

| | | | | | | | | | | | | | | | | | |
|---|---|---|---|---|---|---|---|---|---|---|---|---|---|---|---|---|---|
| SIM16 | <50 bp | 0 | 3 | 2 | 1 | 2 | 2 | 2 | 0 | ≥ 50 bp | 8 | 13 | 9 | 7 | 10 | 14 | 13 | 8 |
| SIM17 | <50 bp | 1 | 1 | 0 | 0 | 0 | 0 | 0 | 0 | ≥ 50 bp | 18 | 8 | 6 | 7 | 6 | 13 | 12 | 9 |
| SIM18 | <50 bp | 0 | 1 | 0 | 0 | 0 | 1 | 0 | 0 | ≥ 50 bp | 12 | 19 | 14 | 15 | 21 | 20 | 17 | 10 |
| SIM19 | <50 bp | 0 | 1 | 2 | 1 | 2 | 3 | 2 | 0 | ≥ 50 bp | 7 | 10 | 9 | 10 | 13 | 11 | 16 | 13 |
| SIM20 | <50 bp | 1 | 1 | 1 | 1 | 2 | 1 | 1 | 0 | ≥ 50 bp | 14 | 10 | 9 | 9 | 11 | 13 | 11 | 12 |
| Average | | 0 | 2 | 1 | 1 | 1 | 1 | 1 | 0 | | 10 | 11 | 9 | 9 | 12 | 14 | 12 | 9 |
| Error rate | | 0.03% | 0.18% | 0.12% | 0.09% | 0.09% | 0.15% | 0.14% | 0.00% | | 0.60% | 0.84% | 0.73% | 0.74% | 0.96% | 1.20% | 1.00% | 2.31% |
| **Missing rate** | | | | | | | | | | | | | | | | | | |
| SIM0 | <50 bp | 82 | 66 | 68 | 68 | 61 | 71 | 69 | 69 | ≥ 50 bp | 10 | 1 | 0 | 0 | 1 | 3 | 2 | 1 |
| SIM1 | <50 bp | 22 | 67 | 67 | 66 | 58 | 75 | 68 | 62 | ≥ 50 bp | 6 | 8 | 6 | 11 | 9 | 9 | 9 | 41<br>3 |
| SIM2 | <50 bp | 14 | 54 | 58 | 58 | 53 | 55 | 64 | 70 | ≥ 50 bp | 4 | 10 | 8 | 13 | 11 | 9 | 9 | 1 |
| SIM3 | <50 bp | 21 | 56 | 58 | 61 | 55 | 69 | 63 | 57 | ≥ 50 bp | 7 | 8 | 5 | 8 | 10 | 9 | 11 | 31 |
| SIM4 | <50 bp | 36 | 64 | 65 | 72 | 65 | 74 | 67 | 63 | ≥ 50 bp | 7 | 10 | 8 | 13 | 12 | 7 | 10 | 33 |
| SIM5 | <50 bp | 21 | 65 | 65 | 70 | 66 | 83 | 68 | 65 | ≥ 50 bp | 5 | 11 | 8 | 16 | 12 | 12 | 8 | 32 |
| SIM6 | <50 bp | 22 | 54 | 58 | 58 | 55 | 64 | 66 | 64 | ≥ 50 bp | 8 | 5 | 5 | 7 | 8 | 10 | 9 | 28 |
| SIM7 | <50 bp | 25 | 53 | 60 | 61 | 57 | 65 | 62 | 60 | ≥ 50 bp | 11 | 10 | 8 | 16 | 13 | 15 | 12 | 27 |
| SIM8 | <50 bp | 24 | 56 | 60 | 61 | 62 | 69 | 69 | 61 | ≥ 50 bp | 8 | 11 | 7 | 7 | 10 | 8 | 7 | 27 |
| SIM9 | <50 bp | 28 | 55 | 54 | 65 | 57 | 77 | 73 | 72 | ≥ 50 bp | 8 | 10 | 9 | 7 | 13 | 13 | 9 | 29 |
| SIM10 | <50 bp | 18 | 68 | 68 | 67 | 62 | 71 | 76 | 66 | ≥ 50 bp | 9 | 9 | 7 | 12 | 9 | 9 | 7 | 32 |
| SIM11 | <50 bp | 17 | 53 | 64 | 56 | 54 | 72 | 70 | 61 | ≥ 50 bp | 6 | 7 | 8 | 14 | 12 | 9 | 10 | 38 |
| SIM12 | <50 bp | 26 | 58 | 64 | 65 | 61 | 70 | 64 | 60 | ≥ 50 bp | 5 | 4 | 4 | 9 | 9 | 9 | 4 | 34 |
| SIM13 | <50 bp | 24 | 59 | 60 | 60 | 55 | 74 | 64 | 69 | ≥ 50 bp | 5 | 6 | 3 | 7 | 4 | 8 | 4 | 27 |
| SIM14 | <50 bp | 19 | 57 | 60 | 65 | 58 | 69 | 63 | 56 | ≥ 50 bp | 6 | 7 | 5 | 11 | 9 | 9 | 10 | 31 |
| SIM15 | <50 bp | 20 | 59 | 64 | 63 | 54 | 67 | 71 | 62 | ≥ 50 bp | 6 | 4 | 4 | 7 | 7 | 6 | 11 | 33 |
| SIM16 | <50 bp | 22 | 62 | 69 | 71 | 66 | 63 | 66 | 64 | ≥ 50 bp | 6 | 12 | 9 | 12 | 14 | 13 | 11 | 25 |
| SIM17 | <50 bp | 26 | 60 | 54 | 65 | 60 | 68 | 67 | 67 | ≥ 50 bp | 5 | 7 | 4 | 9 | 10 | 8 | 8 | 32 |
| SIM18 | <50 bp | 23 | 46 | 44 | 54 | 53 | 67 | 67 | 65 | ≥ 50 bp | 6 | 7 | 6 | 8 | 12 | 10 | 10 | 31 |
| SIM19 | <50 bp | 26 | 64 | 59 | 68 | 61 | 66 | 69 | 62 | ≥ 50 bp | 6 | 9 | 5 | 9 | 9 | 11 | 10 | 30 |
| SIM20 | <50 bp | 23 | 62 | 61 | 67 | 57 | 71 | 70 | 57 | ≥ 50 bp | 8 | 9 | 8 | 11 | 13 | 10 | 11 | 34 |

| | | | | | | | | | | | | | | | | | | |
|---|---|---|---|---|---|---|---|---|---|---|---|---|---|---|---|---|---|---|
| **Average** | | 26 | 59 | 61 | 64 | 59 | 70 | 67 | 63 | | 7 | 8 | 6 | 10 | 10 | 9 | 9 | 30 |
| **Missing rate** | | 2.03% | 6.59% | 6.77% | 7.33% | 6.46% | 8.29% | 8.41% | 30.49% | | 0.40% | 0.62% | 0.48% | 0.82% | 0.79% | 0.82% | 0.75% | 7.92% |

**Accuracy, error, and missing rate for benchmarking analysis for deletions and insertions using *Solanum* phylogeny-based data.**

| | | Insertions | | | | | | | | | Deletions | | | | | | | |
|---|---|---|---|---|---|---|---|---|---|---|---|---|---|---|---|---|---|---|
| | | **SL4** | **M82** | **ZY65** | **ZY56** | **ZY57** | **ZY60** | **ZY61** | **ZY59** | | **SL4** | **M82** | **ZY65** | **ZY56** | **ZY57** | **ZY60** | **ZY61** | **ZY59** |
| **# of retrieved calls** | | **3,356** | **3,245** | **3,149** | **3,036** | **2,906** | **1,695** | **1,340** | **1,106** | | **2,978** | **2,815** | **2,774** | **2,755** | **2,566** | **1,748** | **1,406** | **1,034** |
| **Accuracy** | | | | | | | | | | | | | | | | | | |
| SIM0 | ≥50 bp | 3,351 | 3,241 | 3,144 | 3,031 | 2,900 | 1,691 | 1,338 | 1,104 | ≥ 50 bp | 2,978 | 2,815 | 2,774 | 2,755 | 2,566 | 1,747 | 1,405 | 1,032 |
| SIM1 | ≥50 bp | 3,353 | 3,240 | 3,145 | 3,020 | 2,883 | 1,630 | 1,249 | 1,060 | ≥ 50 bp | 2,977 | 2,808 | 2,770 | 2,735 | 2,555 | 1,716 | 1,355 | 1,000 |
| SIM2 | ≥50 bp | 3,351 | 3,237 | 3,143 | 3,017 | 2,887 | 1,629 | 1,269 | 1,042 | ≥ 50 bp | 2,977 | 2,808 | 2,771 | 2,740 | 2,554 | 1,721 | 1,357 | 1,001 |
| SIM3 | ≥50 bp | 3,354 | 3,240 | 3,147 | 3,021 | 2,880 | 1,630 | 1,272 | 1,058 | ≥ 50 bp | 2,977 | 2,807 | 2,773 | 2,738 | 2,557 | 1,716 | 1,365 | 1,000 |
| SIM4 | ≥50 bp | 3,350 | 3,236 | 3,141 | 3,021 | 2,885 | 1,631 | 1,279 | 1,051 | ≥ 50 bp | 2,977 | 2,809 | 2,772 | 2,738 | 2,559 | 1,715 | 1,348 | 1,003 |
| SIM5 | ≥50 bp | 3,352 | 3,237 | 3,145 | 3,022 | 2,884 | 1,633 | 1,269 | 1,054 | ≥ 50 bp | 2,978 | 2,810 | 2,773 | 2,739 | 2,557 | 1,713 | 1,365 | 1,007 |
| SIM6 | ≥50 bp | 3,346 | 3,231 | 3,140 | 3,014 | 2,878 | 1,626 | 1,266 | 1,047 | ≥ 50 bp | 2,978 | 2,813 | 2,771 | 2,734 | 2,556 | 1,710 | 1,358 | 1,006 |
| SIM7 | ≥50 bp | 3,351 | 3,234 | 3,144 | 3,020 | 2,885 | 1,624 | 1,269 | 1,045 | ≥ 50 bp | 2,977 | 2,807 | 2,771 | 2,741 | 2,556 | 1,718 | 1,352 | 993 |
| SIM8 | ≥50 bp | 3,353 | 3,239 | 3,146 | 3,023 | 2,890 | 1,629 | 1,271 | 1,052 | ≥ 50 bp | 2,977 | 2,808 | 2,772 | 2,732 | 2,555 | 1,713 | 1,359 | 1,003 |
| SIM9 | ≥50 bp | 3,352 | 3,237 | 3,144 | 3,021 | 2,886 | 1,628 | 1,262 | 1,048 | ≥ 50 bp | 2,977 | 2,810 | 2,769 | 2,734 | 2,555 | 1,714 | 1,362 | 1,004 |
| SIM10 | ≥50 bp | 3,349 | 3,238 | 3,142 | 3,019 | 2,878 | 1,621 | 1,263 | 1,042 | ≥ 50 bp | 2,977 | 2,810 | 2,770 | 2,741 | 2,558 | 1,717 | 1,360 | 1,005 |
| SIM11 | ≥50 bp | 3,353 | 3,239 | 3,146 | 3,020 | 2,885 | 1,634 | 1,269 | 1,047 | ≥ 50 bp | 2,978 | 2,808 | 2,771 | 2,736 | 2,558 | 1,714 | 1,359 | 1,002 |
| SIM12 | ≥50 bp | 3,350 | 3,238 | 3,144 | 3,024 | 2,886 | 1,637 | 1,263 | 1,048 | ≥ 50 bp | 2,978 | 2,807 | 2,772 | 2,739 | 2,554 | 1,721 | 1,356 | 1,012 |
| SIM13 | ≥50 bp | 3,350 | 3,238 | 3,143 | 3,010 | 2,883 | 1,624 | 1,266 | 1,053 | ≥ 50 bp | 2,978 | 2,810 | 2,771 | 2,740 | 2,552 | 1,718 | 1,363 | 1,007 |
| SIM14 | ≥50 bp | 3,352 | 3,240 | 3,145 | 3,021 | 2,886 | 1,641 | 1,266 | 1,050 | ≥ 50 bp | 2,977 | 2,806 | 2,772 | 2,741 | 2,554 | 1,715 | 1,363 | 1,003 |
| SIM15 | ≥50 bp | 3,351 | 3,236 | 3,144 | 3,017 | 2,883 | 1,633 | 1,269 | 1,054 | ≥ 50 bp | 2,977 | 2,811 | 2,771 | 2,733 | 2,555 | 1,717 | 1,357 | 1,010 |
| SIM16 | ≥50 bp | 3,351 | 3,237 | 3,143 | 3,012 | 2,881 | 1,626 | 1,262 | 1,051 | ≥ 50 bp | 2,977 | 2,807 | 2,768 | 2,736 | 2,551 | 1,714 | 1,365 | 1,007 |
| SIM17 | ≥50 bp | 3,348 | 3,234 | 3,140 | 3,018 | 2,883 | 1,622 | 1,272 | 1,045 | ≥ 50 bp | 2,977 | 2,808 | 2,771 | 2,738 | 2,552 | 1,719 | 1,363 | 1,002 |
| SIM18 | ≥50 bp | 3,350 | 3,236 | 3,144 | 3,014 | 2,885 | 1,622 | 1,270 | 1,049 | ≥ 50 bp | 2,977 | 2,808 | 2,771 | 2,737 | 2,555 | 1,708 | 1,359 | 1,008 |
| SIM19 | ≥50 bp | 3,351 | 3,234 | 3,142 | 3,012 | 2,882 | 1,623 | 1,264 | 1,043 | ≥ 50 bp | 2,978 | 2,810 | 2,770 | 2,741 | 2,557 | 1,716 | 1,363 | 1,008 |
| SIM20 | ≥50 bp | 3,349 | 3,234 | 3,142 | 3,014 | 2,880 | 1,622 | 1,253 | 1,052 | ≥ 50 bp | 2,977 | 2,810 | 2,772 | 2,742 | 2,557 | 1,712 | 1,361 | 998 |
| **Average** | | 3,351 | 3,237 | 3,144 | 3,019 | 2,884 | 1,631 | 1,270 | 1,052 | | 2,977 | 2,809 | 2,771 | 2,739 | 2,556 | 1,717 | 1,362 | 1,005 |
| **Accuracy** | | 99.85% | 99.75% | 99.83% | 99.43% | 99.25% | 96.24% | 94.74% | 95.13% | | 99.98% | 99.79% | 99.90% | 99.40% | 99.60% | 98.22% | 96.85% | 97.22% |

| | | | | | | | | | | | | | | | | | | |
|---|---|---|---|---|---|---|---|---|---|---|---|---|---|---|---|---|---|---|
| **Error rate** | | | | | | | | | | | | | | | | | | |
| **SIM0** | ≥50 bp | 0 | 0 | 0 | 0 | 1 | 0 | 0 | 0 | ≥ 50 bp | 0 | 0 | 0 | 0 | 0 | 0 | 0 | 0 |
| **SIM1** | ≥50 bp | 0 | 0 | 0 | 0 | 3 | 1 | 6 | 0 | ≥ 50 bp | 1 | 1 | 1 | 6 | 1 | 2 | 3 | 6 |
| **SIM2** | ≥50 bp | 0 | 0 | 0 | 2 | 2 | 0 | 3 | 1 | ≥ 50 bp | 1 | 2 | 1 | 6 | 1 | 2 | 5 | 4 |
| **SIM3** | ≥50 bp | 0 | 0 | 0 | 0 | 2 | 0 | 6 | 0 | ≥ 50 bp | 1 | 2 | 1 | 8 | 1 | 1 | 3 | 6 |
| **SIM4** | ≥50 bp | 0 | 0 | 0 | 1 | 4 | 1 | 2 | 1 | ≥ 50 bp | 1 | 1 | 1 | 8 | 1 | 1 | 3 | 3 |
| **SIM5** | ≥50 bp | 0 | 0 | 0 | 0 | 3 | 0 | 6 | 0 | ≥ 50 bp | 0 | 0 | 0 | 5 | 0 | 1 | 2 | 3 |
| **SIM6** | ≥50 bp | 0 | 0 | 0 | 2 | 2 | 0 | 2 | 0 | ≥ 50 bp | 0 | 0 | 0 | 6 | 0 | 0 | 4 | 2 |
| **SIM7** | ≥50 bp | 0 | 0 | 0 | 2 | 3 | 1 | 7 | 1 | ≥ 50 bp | 1 | 2 | 1 | 5 | 1 | 2 | 6 | 5 |
| **SIM8** | ≥50 bp | 0 | 0 | 0 | 0 | 2 | 1 | 2 | 0 | ≥ 50 bp | 1 | 1 | 1 | 8 | 1 | 1 | 3 | 4 |
| **SIM9** | ≥50 bp | 0 | 0 | 0 | 0 | 3 | 1 | 2 | 0 | ≥ 50 bp | 1 | 1 | 1 | 6 | 1 | 2 | 2 | 5 |
| **SIM10** | ≥50 bp | 0 | 0 | 0 | 1 | 2 | 0 | 6 | 0 | ≥ 50 bp | 1 | 1 | 1 | 5 | 1 | 1 | 2 | 5 |
| **SIM11** | ≥50 bp | 0 | 0 | 0 | 1 | 3 | 0 | 4 | 1 | ≥ 50 bp | 0 | 1 | 0 | 7 | 0 | 1 | 2 | 3 |
| **SIM12** | ≥50 bp | 0 | 0 | 0 | 0 | 1 | 0 | 2 | 0 | ≥ 50 bp | 0 | 1 | 0 | 6 | 0 | 1 | 2 | 2 |
| **SIM13** | ≥50 bp | 0 | 0 | 0 | 1 | 4 | 1 | 3 | 0 | ≥ 50 bp | 0 | 1 | 0 | 3 | 0 | 0 | 2 | 2 |
| **SIM14** | ≥50 bp | 0 | 0 | 0 | 1 | 2 | 2 | 6 | 0 | ≥ 50 bp | 1 | 1 | 1 | 5 | 1 | 2 | 4 | 3 |
| **SIM15** | ≥50 bp | 0 | 0 | 0 | 0 | 3 | 1 | 3 | 0 | ≥ 50 bp | 1 | 1 | 1 | 10 | 1 | 2 | 2 | 4 |
| **SIM16** | ≥50 bp | 0 | 0 | 0 | 1 | 1 | 1 | 3 | 0 | ≥ 50 bp | 1 | 2 | 1 | 8 | 1 | 2 | 3 | 4 |
| **SIM17** | ≥50 bp | 0 | 0 | 0 | 1 | 3 | 2 | 2 | 0 | ≥ 50 bp | 1 | 1 | 1 | 8 | 1 | 1 | 3 | 3 |
| **SIM18** | ≥50 bp | 0 | 0 | 0 | 1 | 2 | 2 | 2 | 1 | ≥ 50 bp | 1 | 2 | 1 | 8 | 1 | 2 | 4 | 2 |
| **SIM19** | ≥50 bp | 0 | 0 | 0 | 1 | 2 | 0 | 8 | 0 | ≥ 50 bp | 0 | 1 | 0 | 2 | 0 | 1 | 0 | 4 |
| **SIM20** | ≥50 bp | 0 | 0 | 0 | 1 | 1 | 1 | 3 | 0 | ≥ 50 bp | 1 | 1 | 1 | 9 | 1 | 2 | 3 | 6 |
| **Average** | | 0 | 0 | 0 | 1 | 2 | 1 | 4 | 0 | | 1 | 1 | 1 | 6 | 1 | 1 | 3 | 4 |
| **Error rate** | | 0.00% | 0.00% | 0.00% | 0.03% | 0.08% | 0.04% | 0.28% | 0.02% | | 0.02% | 0.04% | 0.02% | 0.22% | 0.03% | 0.07% | 0.20% | 0.35% |
| **Missing rate** | | | | | | | | | | | | | | | | | | |
| **SIM0** | ≥50 bp | 5 | 4 | 5 | 5 | 5 | 4 | 2 | 2 | ≥ 50 bp | 0 | 0 | 0 | 0 | 0 | 1 | 1 | 2 |
| **SIM1** | ≥50 bp | 3 | 5 | 4 | 16 | 20 | 64 | 85 | 46 | ≥ 50 bp | 0 | 6 | 3 | 14 | 10 | 30 | 48 | 28 |
| **SIM2** | ≥50 bp | 5 | 8 | 6 | 17 | 17 | 66 | 68 | 63 | ≥ 50 bp | 0 | 5 | 2 | 9 | 11 | 25 | 44 | 29 |
| **SIM3** | ≥50 bp | 2 | 5 | 2 | 15 | 24 | 65 | 62 | 48 | ≥ 50 bp | 0 | 6 | 0 | 9 | 8 | 31 | 38 | 28 |
| **SIM4** | ≥50 bp | 6 | 9 | 8 | 14 | 17 | 63 | 59 | 54 | ≥ 50 bp | 0 | 5 | 1 | 9 | 6 | 32 | 55 | 28 |

| | | | | | | | | | | | | | | | | | |
|---|---|---|---|---|---|---|---|---|---|---|---|---|---|---|---|---|---|
| **SIM5** | ≥50 bp | 4 | 8 | 4 | 14 | 19 | 62 | 65 | 52 | ≥ 50 bp | 0 | 5 | 1 | 11 | 9 | 34 | 39 | 24 |
| **SIM6** | ≥50 bp | 10 | 14 | 9 | 20 | 26 | 69 | 72 | 59 | ≥ 50 bp | 0 | 2 | 3 | 15 | 10 | 38 | 44 | 26 |
| **SIM7** | ≥50 bp | 5 | 11 | 5 | 14 | 18 | 70 | 64 | 60 | ≥ 50 bp | 0 | 6 | 2 | 9 | 9 | 28 | 48 | 36 |
| **SIM8** | ≥50 bp | 3 | 6 | 3 | 13 | 14 | 65 | 67 | 54 | ≥ 50 bp | 0 | 6 | 1 | 15 | 10 | 34 | 44 | 27 |
| **SIM9** | ≥50 bp | 4 | 8 | 5 | 15 | 17 | 66 | 76 | 58 | ≥ 50 bp | 0 | 4 | 4 | 15 | 10 | 32 | 42 | 25 |
| **SIM10** | ≥50 bp | 7 | 7 | 7 | 16 | 26 | 74 | 71 | 64 | ≥ 50 bp | 0 | 4 | 3 | 9 | 7 | 30 | 44 | 24 |
| **SIM11** | ≥50 bp | 3 | 6 | 3 | 15 | 18 | 61 | 67 | 58 | ≥ 50 bp | 0 | 6 | 3 | 12 | 8 | 33 | 45 | 29 |
| **SIM12** | ≥50 bp | 6 | 7 | 5 | 12 | 19 | 58 | 75 | 58 | ≥ 50 bp | 0 | 7 | 2 | 10 | 12 | 26 | 48 | 20 |
| **SIM13** | ≥50 bp | 6 | 7 | 6 | 25 | 19 | 70 | 71 | 53 | ≥ 50 bp | 0 | 4 | 3 | 12 | 14 | 30 | 41 | 25 |
| **SIM14** | ≥50 bp | 4 | 5 | 4 | 14 | 18 | 52 | 68 | 56 | ≥ 50 bp | 0 | 8 | 1 | 9 | 11 | 31 | 39 | 28 |
| **SIM15** | ≥50 bp | 5 | 9 | 5 | 19 | 20 | 61 | 68 | 52 | ≥ 50 bp | 0 | 3 | 2 | 12 | 10 | 29 | 47 | 20 |
| **SIM16** | ≥50 bp | 5 | 8 | 6 | 23 | 24 | 68 | 75 | 55 | ≥ 50 bp | 0 | 6 | 5 | 11 | 14 | 32 | 38 | 23 |
| **SIM17** | ≥50 bp | 8 | 11 | 9 | 17 | 20 | 71 | 66 | 61 | ≥ 50 bp | 0 | 6 | 2 | 9 | 13 | 28 | 40 | 29 |
| **SIM18** | ≥50 bp | 6 | 9 | 5 | 21 | 19 | 71 | 68 | 56 | ≥ 50 bp | 0 | 5 | 2 | 10 | 10 | 38 | 43 | 24 |
| **SIM19** | ≥50 bp | 5 | 11 | 7 | 23 | 22 | 72 | 68 | 63 | ≥ 50 bp | 0 | 4 | 4 | 12 | 9 | 31 | 43 | 22 |
| **SIM20** | ≥50 bp | 7 | 11 | 7 | 21 | 25 | 72 | 84 | 54 | ≥ 50 bp | 0 | 4 | 1 | 4 | 8 | 34 | 42 | 30 |
| **Average** | | 5 | 8 | 5 | 17 | 19 | 63 | 67 | 54 | | 0 | 5 | 2 | 10 | 9 | 30 | 42 | 25 |
| **Missing rate** | | 0.15% | 0.25% | 0.17% | 0.55% | 0.67% | 3.72% | 4.98% | 4.85% | | 0.00% | 0.17% | 0.08% | 0.37% | 0.37% | 1.71% | 2.96% | 2.43% |

#### Supplementary Table 11: Benchmark SV detection in maize using SVGAP and paftools.

| Methods | # of simulated SVs | # of Predicted SVs | Recall | Accuracy | F1 score |
|---|---|---|---|---|---|
| *Simulated 2000 SVs (deletions and insertions) on chr10 without additional mutations at the flanking region of breakpoints* | | | | | |
| minimap2_default | 2,000 | 2,053 | 1,837/2,000 (91.85%) | 1,837/2,053 (89.5%) | **90.65%** |
| minimap2_asm5 | 2,000 | 2,601 | 1,740/2,000 (87%) | 1,740/2,601 (66.9%) | 75.64% |
| minimap2_asm10 | 2,000 | 2,343 | 1,894/2,000 (94.7%) | 1,894/2,343 (80.8%) | 87.22% |
| minimap2_asm20 | 2,000 | 2,245 | 1,906/2,000 (95.3%) | 1,906/2,245 (84.9%) | 89.80% |
| paftools_default | 2,000 | 1,581 | 1,488/2,000 (74.4%) | 1,488/1,581 (94.1%） | 83.11% |
| paftools_asm5 | 2,000 | 1,110 | 980/2,000 (49%) | 980/1,110 (88.3%) | 63.02% |
| paftools_asm10 | 2,000 | 1,683 | 1,553/2,000 (77.65%) | 1,553/1,683 (92.3%) | 84.33% |
| paftools_asm20 | 2,000 | 1,603 | 1,525/2,000 (76.25%) | 1,525/1,603 (95.1%) | 84.65% |
| *Simulated 3000 SVs (deletions and insertions) on chr10 without additional mutations at the flanking region of breakpoints* | | | | | |
| minimap2_default | 3,000 | 2,979 | 2,944/3,000 (98.1%) | 2,944/2,979 (98.8%) | **98.48%** |
| minimap2_asm5 | 3,000 | 2,963 | 2,931/3,000 (97.7%) | 2,931/2,963 (98.9%) | 98.31% |
| minimap2_asm10 | 3,000 | 2,963 | 2,933/3,000 (97.8%) | 2,933/2,963 (99.0%) | 98.37% |
| minimap2_asm20 | 3,000 | 2,969 | 2,933/3,000 (97.8%) | 2,933/2,969 (98.8%) | 98.27% |
| paftools_default | 3,000 | 4,584 | 1,896/3,000 (63.2%) | 1,896/4,584 (41.4%) | 50.00% |
| paftools_asm5 | 3,000 | 2,058 | 1,907/3,000 (63.6%) | 1,907/2,058 (92.7%) | 75.41% |
| paftools_asm10 | 3,000 | 2,423 | 1,907/3,000 (63.6%) | 1,907/2,423 (78.7%) | 70.33% |
| paftools_asm20 | 3,000 | 2,178 | 1,887/3,000 (62.9%) | 1,887/2,178 (86.6%) | 72.89% |

Recall was determined based on an 80% reciprocal overlap cutoff.

**Supplementary Table 12:** SV detected between 25 NAM maize lines to the reference B73 using SVGAP.

| Pairwise comparison | DEL | INS | DUP | INV | TRL |
|---|---|---|---|---|---|
| **B73 vs. Ab10** | 1,316 | 3,931 | 54 | 28 | 58 |
| **B73 vs. B97** | 28,955 | 34,928 | 427 | 80 | 379 |
| **B73 vs. CML103** | 32,127 | 36,620 | 433 | 87 | 376 |
| **B73 vs. CML228** | 33,229 | 38,509 | 548 | 93 | 349 |
| **B73 vs. CML247** | 33,688 | 38,876 | 511 | 105 | 365 |
| **B73 vs. CML277** | 33,391 | 37,726 | 479 | 91 | 359 |
| **B73 vs. CML322** | 32,660 | 38,546 | 466 | 125 | 389 |
| **B73 vs. CML333** | 32,039 | 38,515 | 484 | 114 | 417 |
| **B73 vs. CML52** | 31,792 | 39,508 | 576 | 102 | 397 |
| **B73 vs. CML69** | 33,093 | 38,463 | 461 | 111 | 377 |
| **B73 vs. HP301** | 31,358 | 34,879 | 461 | 91 | 359 |
| **B73 vs. Il14H** | 32,426 | 36,064 | 469 | 87 | 388 |
| **B73 vs. Ki11** | 33,394 | 41,436 | 498 | 147 | 438 |
| **B73 vs. Ki3** | 33,203 | 37,768 | 479 | 102 | 375 |
| **B73 vs. Ky21** | 29,168 | 32,941 | 466 | 87 | 335 |
| **B73 vs. M162W** | 31,780 | 36,891 | 440 | 83 | 346 |
| **B73 vs. M37W** | 31,663 | 36,690 | 434 | 96 | 344 |
| **B73 vs. Mo18W** | 32,419 | 38,794 | 492 | 115 | 393 |
| **B73 vs. Ms71** | 30,084 | 35,386 | 441 | 81 | 361 |
| **B73 vs. NC350** | 33,096 | 42,040 | 504 | 135 | 473 |
| **B73 vs. NC358** | 32,360 | 38,234 | 499 | 99 | 353 |
| **B73 vs. Oh43** | 29,835 | 34,063 | 457 | 111 | 356 |
| **B73 vs. Oh7b** | 28,266 | 32,372 | 431 | 78 | 304 |
| **B73 vs. P39** | 32,819 | 36,596 | 466 | 119 | 405 |
| **B73 vs. Tx303** | 31,745 | 37,879 | 485 | 115 | 375 |
| **B73 vs. Tzi8** | 32,645 | 38,435 | 558 | 93 | 379 |
| **Total** | 798,551 | 936,090 | 12,019 | 2,575 | 9,450 |
| **Non-redundant** | 192,455 | 312,188 | 6,612 | 2,084 | - |

**Supplementary Table 13: Runtime and memory requirements for each step in SVGAP applied to 49 rice genomes.**

| Pipeline steps | User time (Hour) | System time (Hour) | Elapsed (wall clock) time (Hour) | Maximum memory (Gb) | Threads |
|---|---|---|---|---|---|
| Alignment per sample (minimap2) | 1.28 | 0.02 | 0.16 | 71.29 | 40 |
| 1_Convert2Axt | 0.40 | 0.05 | 0.02 | 0.45 | 40 |
| 2_ChainNetSyn | 0.34 | 0.03 | 0.37 | 0.78 | - |
| 3_SynNetFilter | 0.16 | 0.03 | 0.19 | 0.78 | - |
| 4_PairwiseSV | 136.14 | 0.12 | 4.41 | 1.19 | 40 |
| 5_Combined | 3.11 | 0.05 | 2.47 | 5.51 | - |
| 6_DELGenoptyping | 430.35 | 244.63 | 63.85 | 15.07 | 12 |
| 7_INSGenotyping | 279.25 | 384.91 | 57.18 | 15.16 | 12 |
| 8_InDelGenotyping_del | 10.04 | 0.47 | 1.39 | 15.68 | 12 |
| 8_InDelGenotyping_ins | 9.53 | 0.44 | 1.26 | 15.07 | 12 |
| 9_SNPGenotyping | 11.22 | 0.52 | 1.71 | 15.16 | 12 |

### Supplementary Figures

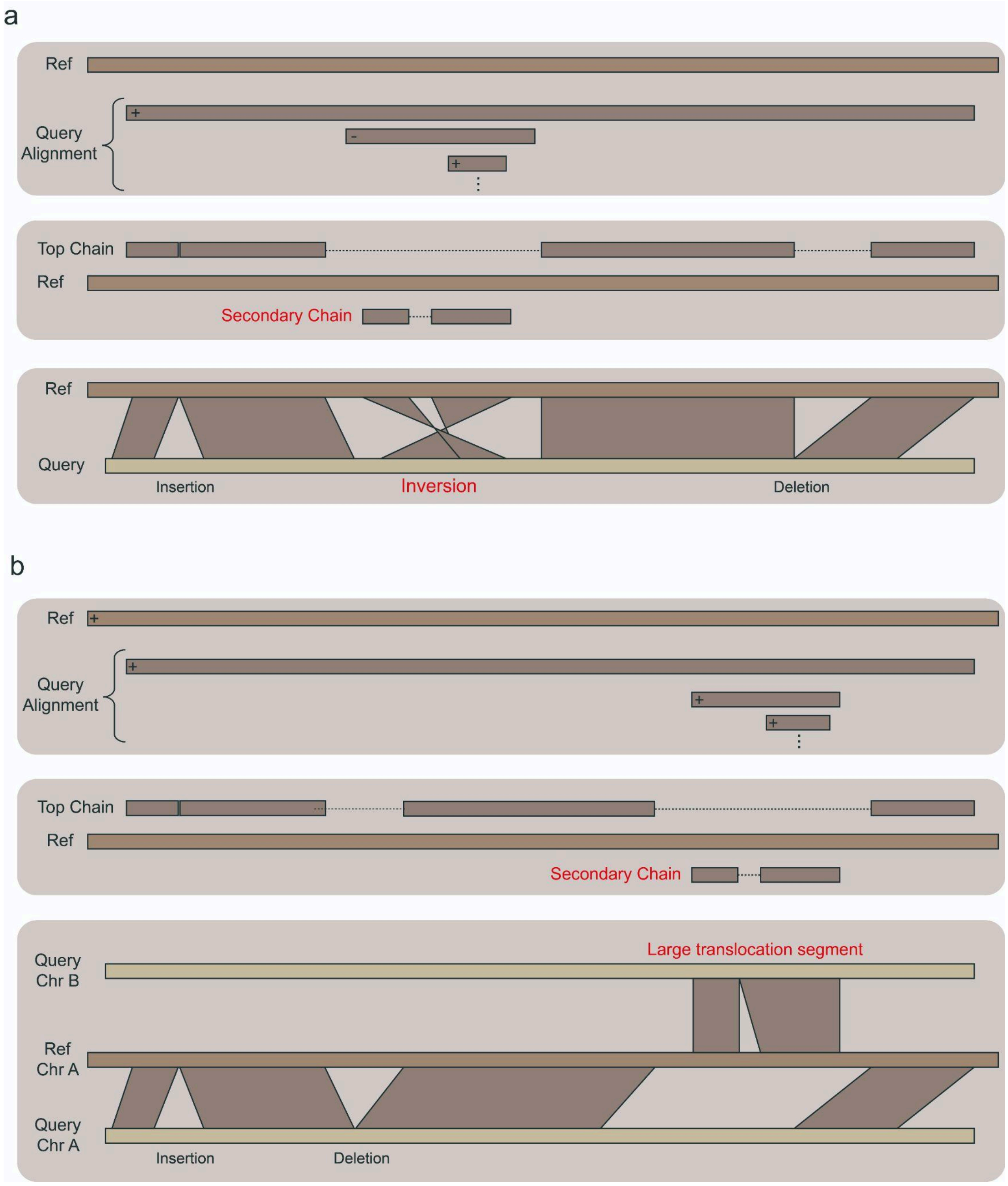


**Supplementary Figure 1: SVGAP facilitates structural variant (SV) identification in inversion and large translocation segments.** In most cases, SVs were detected from the sequence alignments in the top chain. However, some regions exhibited large inversions (**a**) or translocations (**b**), which are more likely to appear as lower (i.e., secondary) chains. These regions are rearranged genomic areas that remain syntenic and orthologous, indicating that the SVs within them should be identified. SVGAP is capable of identifying SVs in these regions and constructing a comprehensive SV dataset.

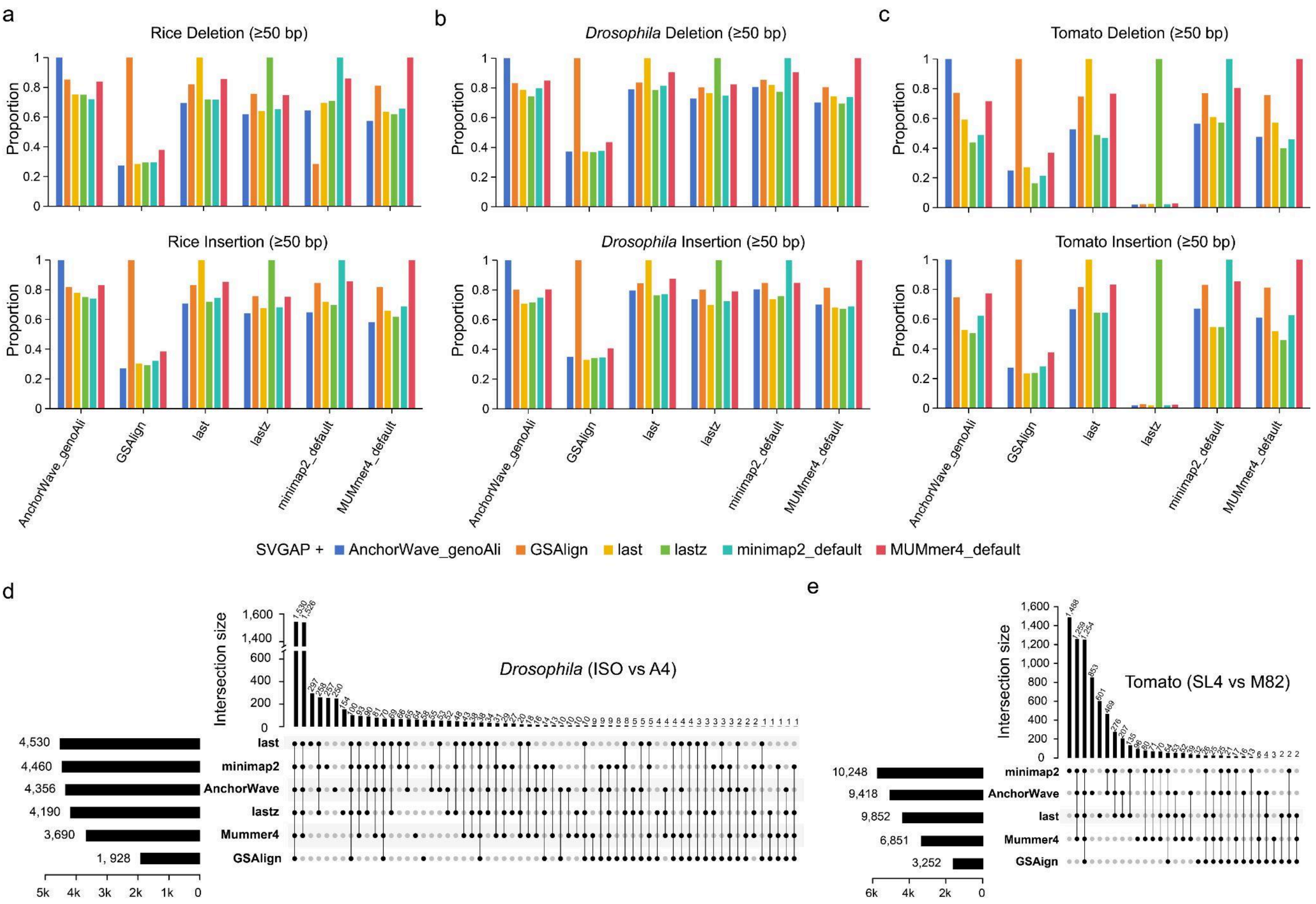


**Supplementary Figure 2: Comparison of SVGAP with different aligners for identifying SVs between real genomes in rice, *Drosophila*, and tomato. a**. Pairwise overlap of SVs (deletions and insertions) identified by SVGAP using alignment results from different aligners between rice genome assemblies: MH63 and ZS97. **b**. Pairwise overlap of SVs identified by SVGAP using alignment results from different aligners between two *Drosophila* genome assemblies: iso-1 and A4. **c**. Pairwise overlap of SVs identified by SVGAP using alignment results from different aligners between two tomato genome assemblies: SL4 and M82. **d**. An upset plot showing the analysis of shared calls between different aligners on two Drosophila data. **e**. An upset plot showing the analysis of shared calls between different aligners on tomato data. The exact numbers are available in Supplementary Table 4.

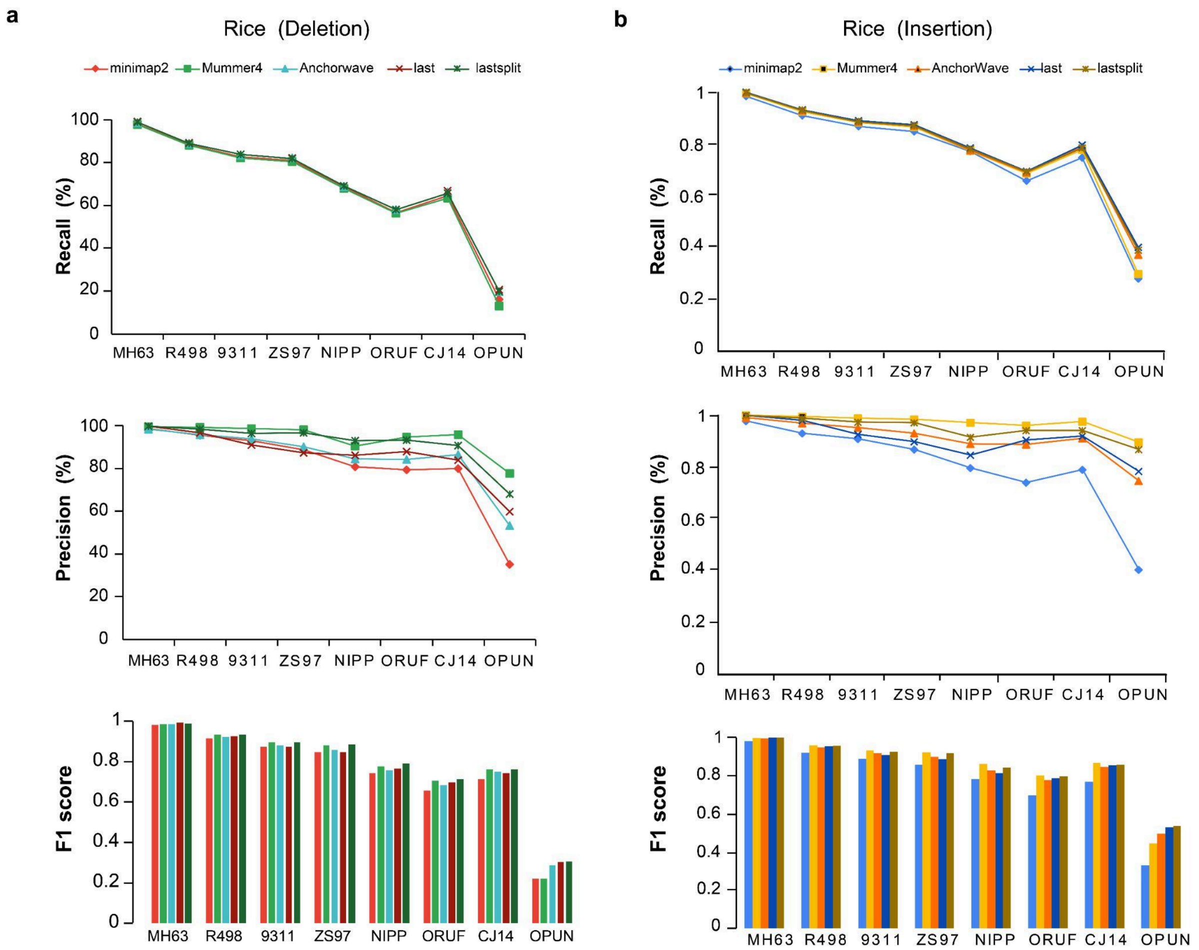


| | | | | | | | | |
|---|---|---|---|---|---|---|---|---|
| minimap2 | 0.9826 | 0.9170 | 0.8755 | 0.8462 | 0.7420 | 0.6593 | 0.7147 | 0.2201 |
| Mummer4 | 0.9854 | 0.9329 | 0.8962 | 0.8833 | 0.7754 | 0.7065 | 0.7629 | 0.2202 |
| Anchorwave | 0.9848 | 0.9219 | 0.8826 | 0.8588 | 0.7581 | 0.6855 | 0.7500 | 0.2866 |
| last | 0.9938 | 0.9270 | 0.8727 | 0.8465 | 0.7660 | 0.6977 | 0.7437 | 0.3051 |
| last + split | 0.9915 | 0.9333 | 0.8959 | 0.8851 | 0.7925 | 0.7142 | 0.7621 | 0.3073 |

| | | | | | | | | |
|---|---|---|---|---|---|---|---|---|
| minimap2 | 0.9807 | 0.9194 | 0.8878 | 0.8570 | 0.7834 | 0.6957 | 0.7675 | 0.3299 |
| Mummer4 | 0.9970 | 0.9579 | 0.9324 | 0.9210 | 0.8614 | 0.8001 | 0.8655 | 0.4448 |
| AnchorWave | 0.9942 | 0.9479 | 0.9166 | 0.8995 | 0.8281 | 0.7758 | 0.8451 | 0.4963 |
| last | 0.9987 | 0.9549 | 0.9074 | 0.8865 | 0.8134 | 0.7859 | 0.8531 | 0.5285 |
| last+split | 0.9982 | 0.9571 | 0.9266 | 0.9172 | 0.8415 | 0.7970 | 0.8559 | 0.5366 |

**Supplementary Figure 3: Benchmarking SVGAP with different aligners for SV calling using *Oryza* phylogeny-based simulation data. a.** Recall, precision, and F1 score for deletions. **b**. Recall, precision, and F1 score for insertions. The exact numbers are available in Supplementary Table 6.

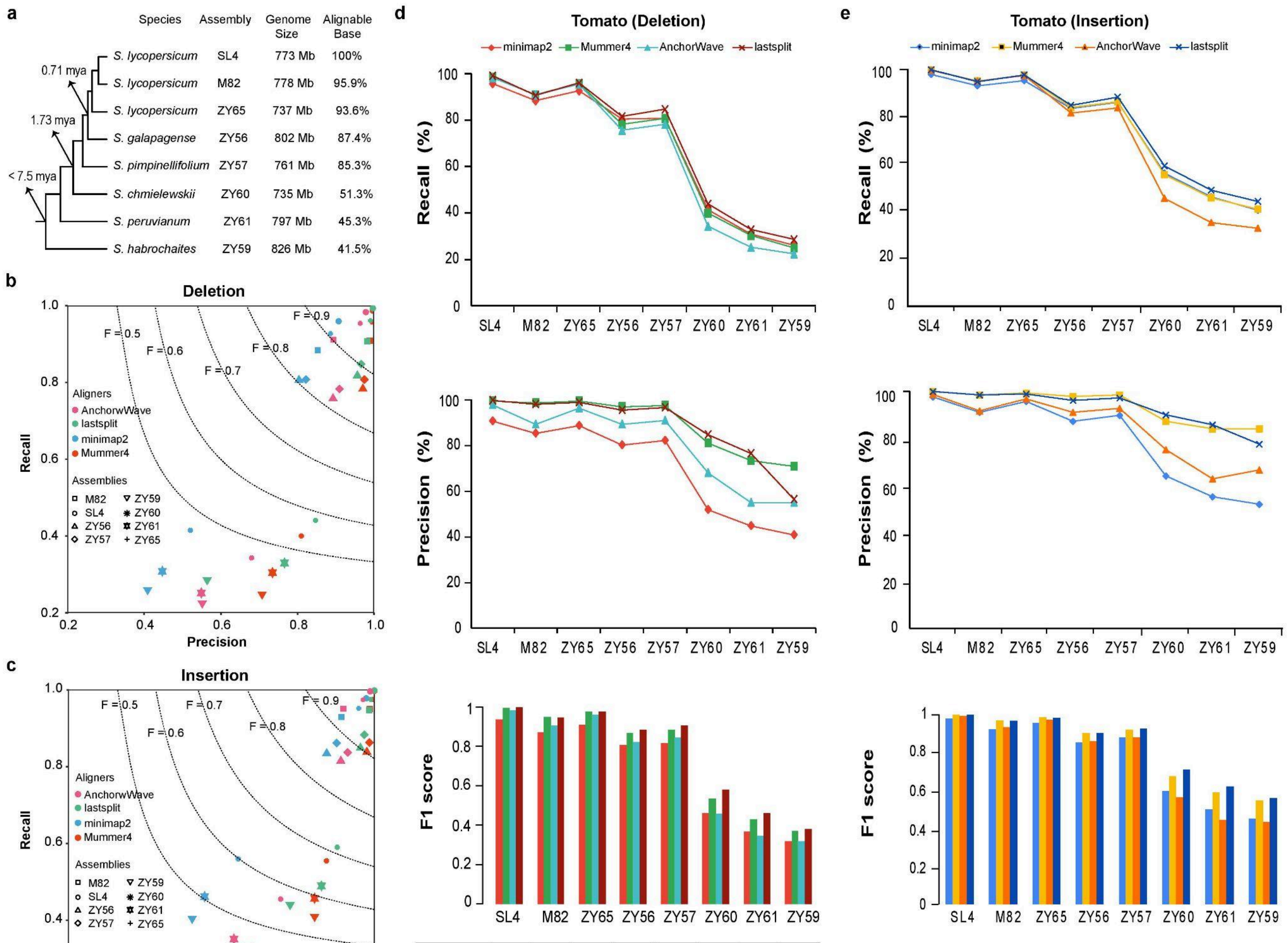


| | SL4 | M82 | ZY65 | ZY56 | ZY57 | ZY60 | ZY61 | ZY59 |
|---|---|---|---|---|---|---|---|---|
| minimap2 | 0.9328 | 0.8683 | 0.9065 | 0.8051 | 0.8148 | 0.4618 | 0.3649 | 0.3177 |
| Mummer4 | 0.9919 | 0.9470 | 0.9757 | 0.8670 | 0.8833 | 0.5357 | 0.4306 | 0.3676 |
| Anchorwave | 0.9808 | 0.9026 | 0.9589 | 0.8201 | 0.8417 | 0.4560 | 0.3454 | 0.3195 |
| last + split | 0.9953 | 0.9433 | 0.9755 | 0.8816 | 0.9032 | 0.5801 | 0.4613 | 0.3791 |

| | SL4 | M82 | ZY65 | ZY56 | ZY57 | ZY60 | ZY61 | ZY59 |
|---|---|---|---|---|---|---|---|---|
| minimap2 | 0.9778 | 0.9208 | 0.9548 | 0.8538 | 0.8806 | 0.5986 | 0.5041 | 0.4561 |
| Mummer4 | 0.9981 | 0.9678 | 0.9849 | 0.9029 | 0.9200 | 0.6783 | 0.5925 | 0.5507 |
| AnchorWave | 0.9919 | 0.9341 | 0.9718 | 0.8601 | 0.8806 | 0.5673 | 0.4511 | 0.4391 |
| last+split | 0.9988 | 0.9659 | 0.9821 | 0.9019 | 0.9255 | 0.7132 | 0.6236 | 0.5621 |

**Supplementary Figure 4: Benchmarking SVGAP with different aligners for SV calling using *Solanum* phylogeny-based simulation data. a.** The *Solanum* genomes, phylogenetic relationship and sequence divergence used for benchmark analysis. **b.** Recall, precision, and F1 score for deletions. **c**. Recall, precision, and F1 score for insertions. **d** and **e** showing the trend of each metric changes as sequence divergence. The exact numbers are available in Supplementary Table 6.

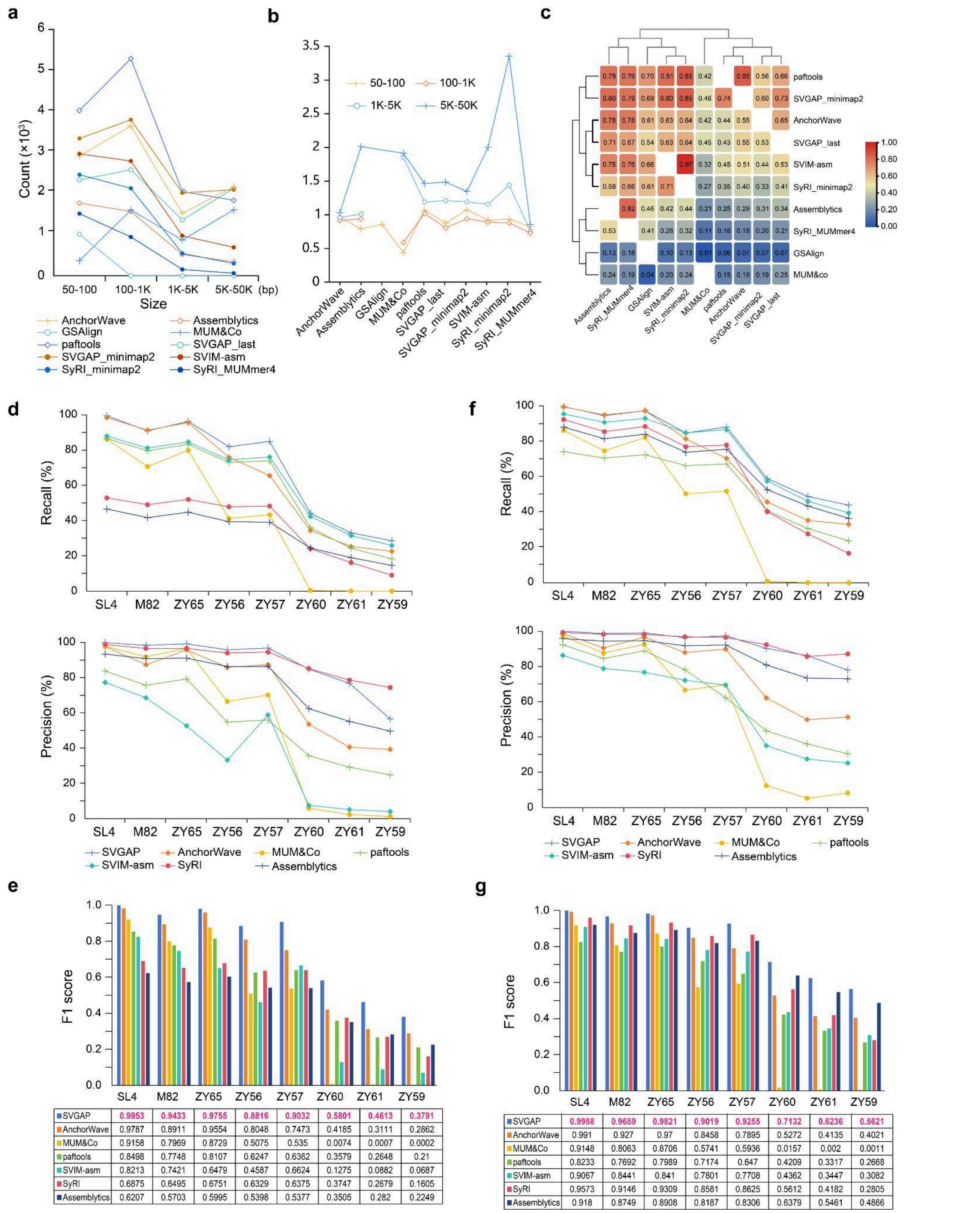


**e**

| | SL4 | M82 | ZY65 | ZY56 | ZY57 | ZY60 | ZY61 | ZY59 |
|---|---|---|---|---|---|---|---|---|
| SVGAP | 0.9953 | 0.9433 | 0.9755 | 0.8816 | 0.9032 | 0.5801 | 0.4613 | 0.3791 |
| AnchorWave | 0.9787 | 0.8911 | 0.9554 | 0.8048 | 0.7473 | 0.4185 | 0.3111 | 0.2862 |
| MUM&Co | 0.9158 | 0.7969 | 0.8729 | 0.5075 | 0.535 | 0.0074 | 0.0007 | 0.0002 |
| paftools | 0.8498 | 0.7748 | 0.8107 | 0.6247 | 0.6362 | 0.3579 | 0.2648 | 0.21 |
| SVIM-asm | 0.8213 | 0.7421 | 0.6479 | 0.4587 | 0.6624 | 0.1275 | 0.0882 | 0.0687 |
| SyRI | 0.6875 | 0.6495 | 0.6751 | 0.6329 | 0.6375 | 0.3747 | 0.2679 | 0.1605 |
| Assemblytics | 0.6207 | 0.5703 | 0.5995 | 0.5398 | 0.5377 | 0.3505 | 0.282 | 0.2249 |

**g**

| | SL4 | M82 | ZY65 | ZY56 | ZY57 | ZY60 | ZY61 | ZY59 |
|---|---|---|---|---|---|---|---|---|
| SVGAP | 0.9988 | 0.9659 | 0.9821 | 0.9019 | 0.9255 | 0.7132 | 0.6236 | 0.5621 |
| AnchorWave | 0.991 | 0.927 | 0.97 | 0.8458 | 0.7895 | 0.5272 | 0.4135 | 0.4021 |
| MUM&Co | 0.9148 | 0.8063 | 0.8706 | 0.5741 | 0.5936 | 0.0157 | 0.002 | 0.0011 |
| paftools | 0.8233 | 0.7692 | 0.7989 | 0.7174 | 0.647 | 0.4209 | 0.3317 | 0.2668 |
| SVIM-asm | 0.9067 | 0.8441 | 0.841 | 0.7801 | 0.7708 | 0.4362 | 0.3447 | 0.3082 |
| SyRI | 0.9573 | 0.9146 | 0.9309 | 0.8581 | 0.8625 | 0.5612 | 0.4182 | 0.2805 |
| Assemblytics | 0.918 | 0.8749 | 0.8908 | 0.8187 | 0.8306 | 0.6379 | 0.5461 | 0.4866 |

**Supplementary Figure 5: Comparison of SVGAP with other methods using benchmark analysis based on simulated data from tomato and *Solanum* phylogeny. a**. Number of SV calls reported by different methods across various length ranges when comparing the two tomato genomes, SL4 and M82. **b**. Ratio of deletions to insertions reported by different methods based on the two tomato genomes. **c**. Overlap of SVs in pairwise comparisons among different methods. **d**. Recall and precision for the benchmark analysis of deletions using simulated data from *Solanum* phylogeny. **e**. F1 score for deletions. **f**. Recall and precision for insertions. **g**. F1 score for insertions.

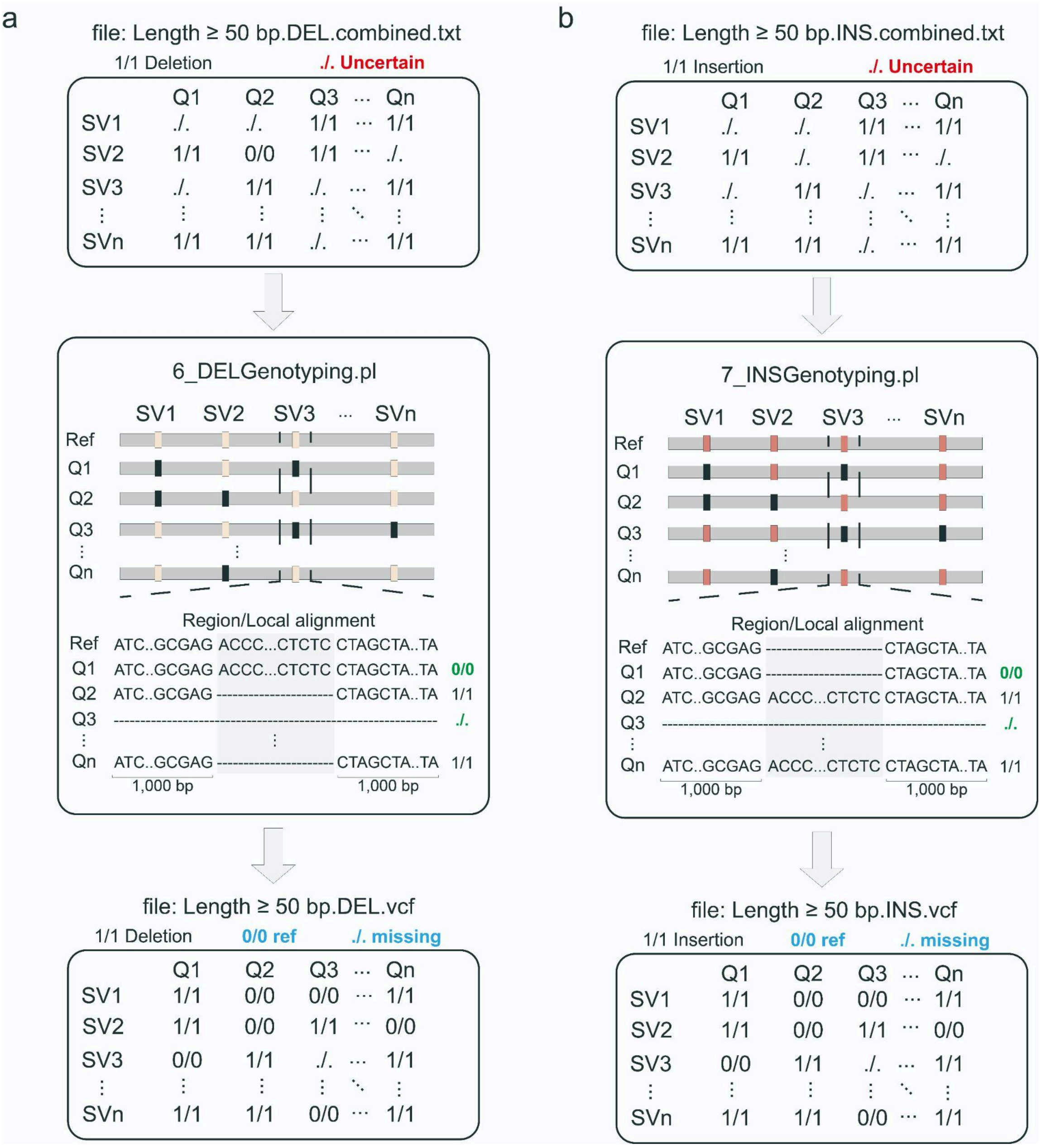


**Supplementary Figure 6: The diagram illustrates the strategy of population genotyping for deletions and insertions.** The genotyping step takes a pre-combined SV dataset as input and re-genotypes each SV call across all samples by extracting local alignments around the SV site from the corresponding pairwise single-coverage genome alignments. For each sample and SV, SVGAP reports three genotypes: 1/1 (alternative type), 0/0 (reference type), and ./. (missing or unable to determine).

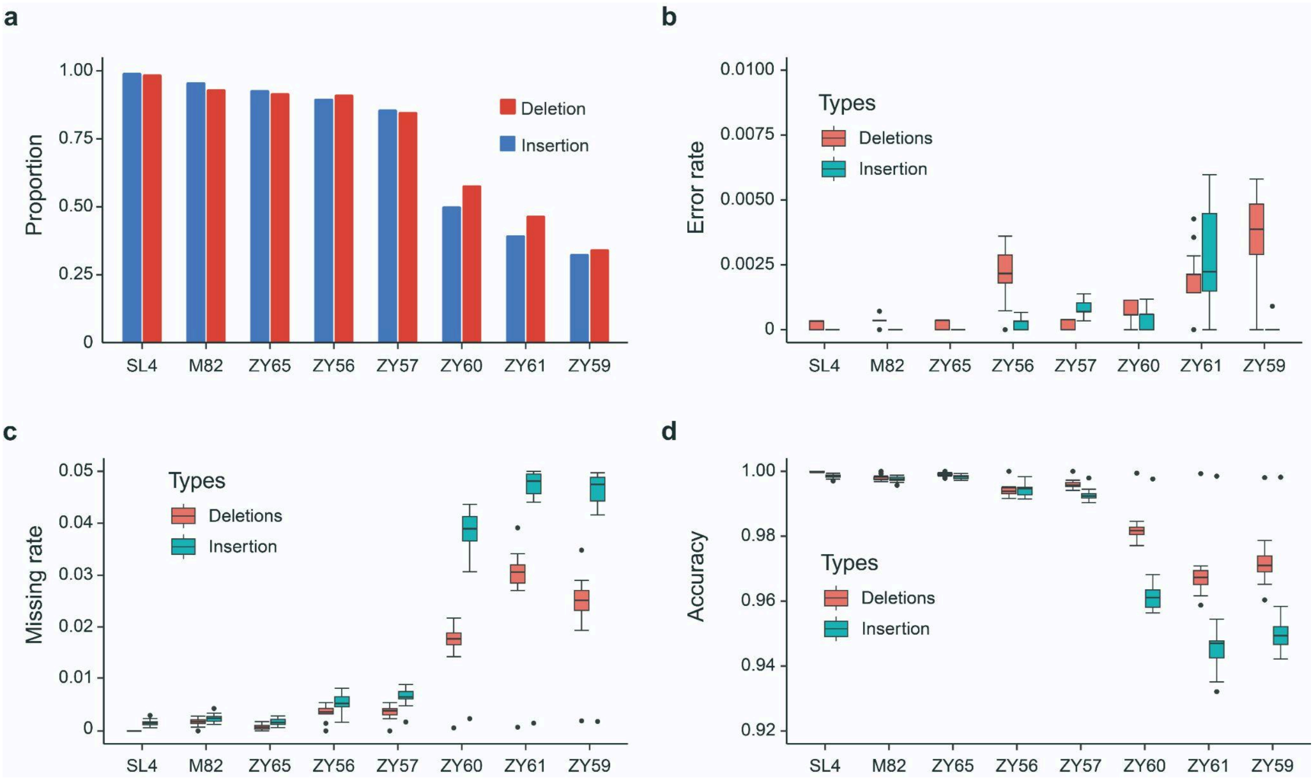


**Supplementary Figure 7: Benchmark analysis of the merging and genotyping functions in SVGAP with tomato data. a**, Proportion of retrieved simulated calls in the merging step by comparing the reference SL4 and eight other *Solanum* genomes of varying sequence divergence. **b**, Error rate of the genotyping step with population-scale simulating data based on *Solanum* phylogeny. **c**, The missing rate of the genotyping step across genomes of varying sequence divergence. **d**. The accuracy of the genotyping step across genomes of varying sequence divergence.

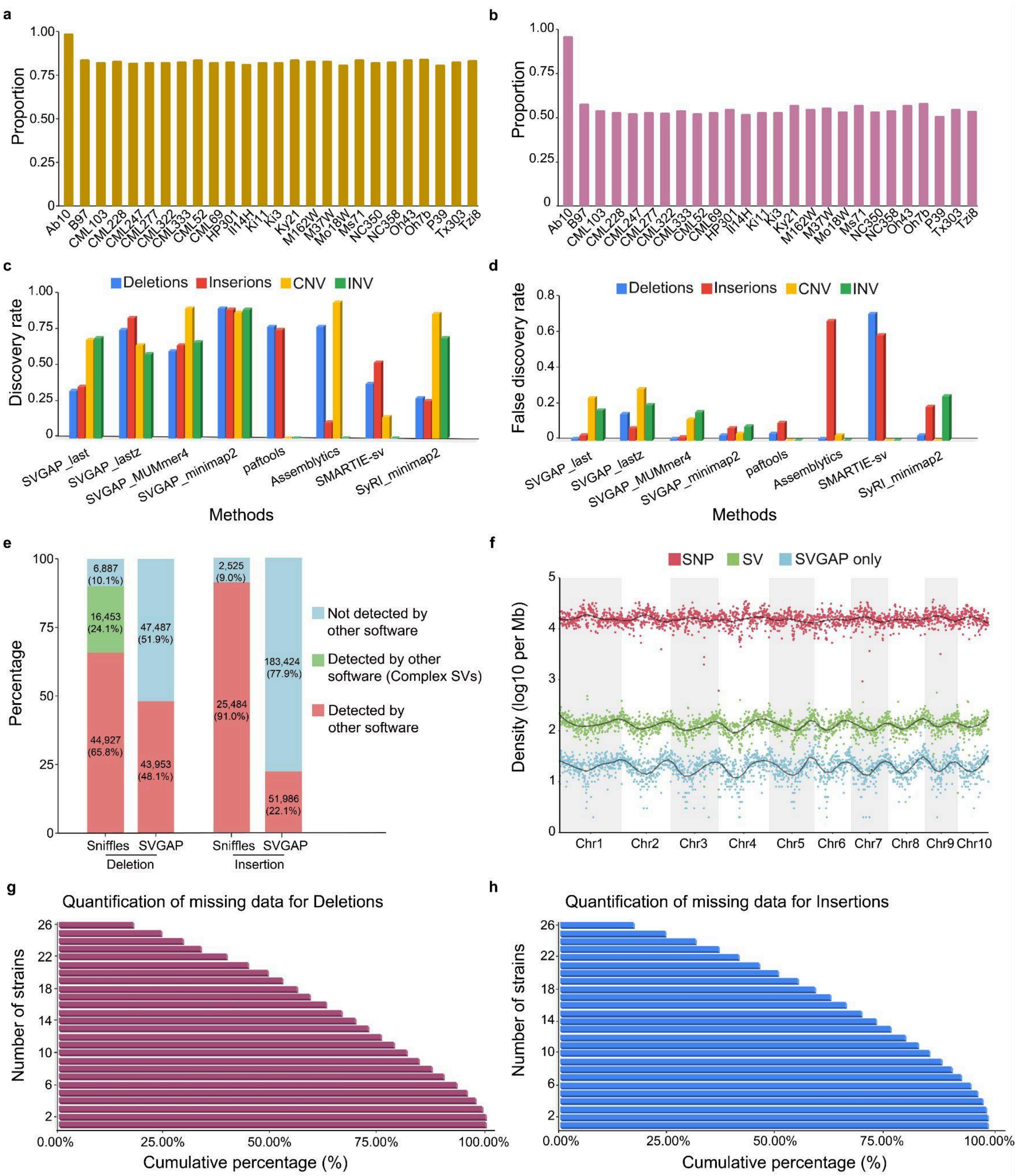


**Supplementary Figure 8: Structural variant (SV) identification across 26 maize genomes. a.** Proportion of genome regions alignable between B73 and other 26 maize genomes. **b.** Proportion of genome regions alignable in synteny between B73 and other 26 maize genomes. **c.** Recall (discovery rate) for benchmarking SV detection across different SV types in maize. **d.** False discovery rate (FDR) for benchmarking SV detection across SV types in maize. **e.** Comparison of SV calls between a previous study [10] using Sniffles and SVGAP calls from this study. **f.** Chromosome-level distribution of SVs and SNPs, highlighting SVs uniquely detected by SVGAP. **g.** Cumulative proportion of deletions identified across 26 maize genomes (based on chromosome 1 deletions). **h.** Cumulative proportion of insertions identified across 26 maize genomes (based on chromosome 2 insertions).

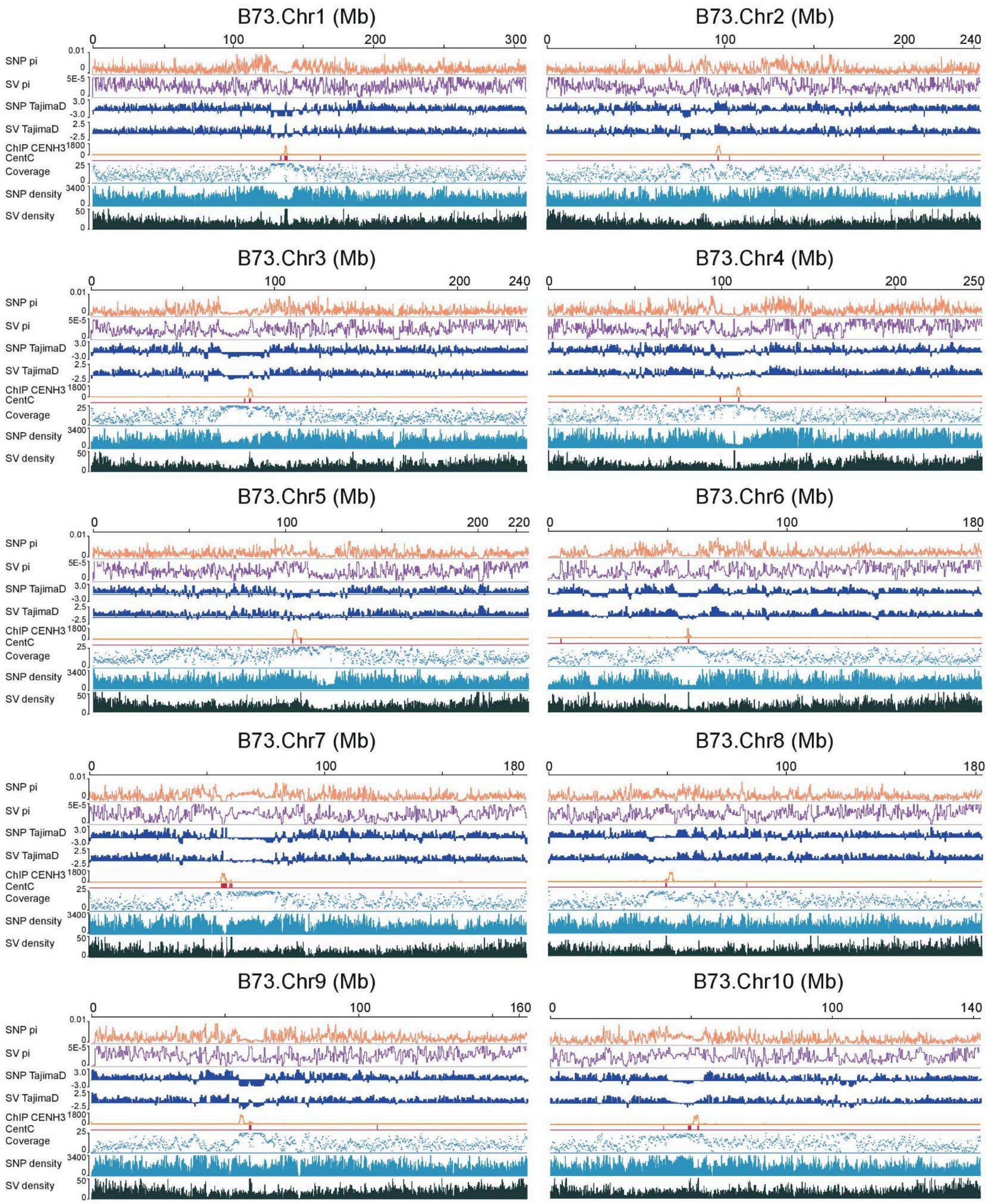


**Supplementary Figure 9: Population properties of genetic variants identified by SVGAP across the maize genome.** The average pairwise diversity (π), Tajima's D, as well as SNP and SV density were calculated in 100-kb non-overlapping windows along each chromosome. Coverage is represented by the number of genomes that can be syntenically aligned to the B73 reference, calculated for each 10-kb window. Notably, an elevated coverage was observed around the functional centromeric regions across most chromosomes, highlighting the structural and evolutionary conservation of these regions.

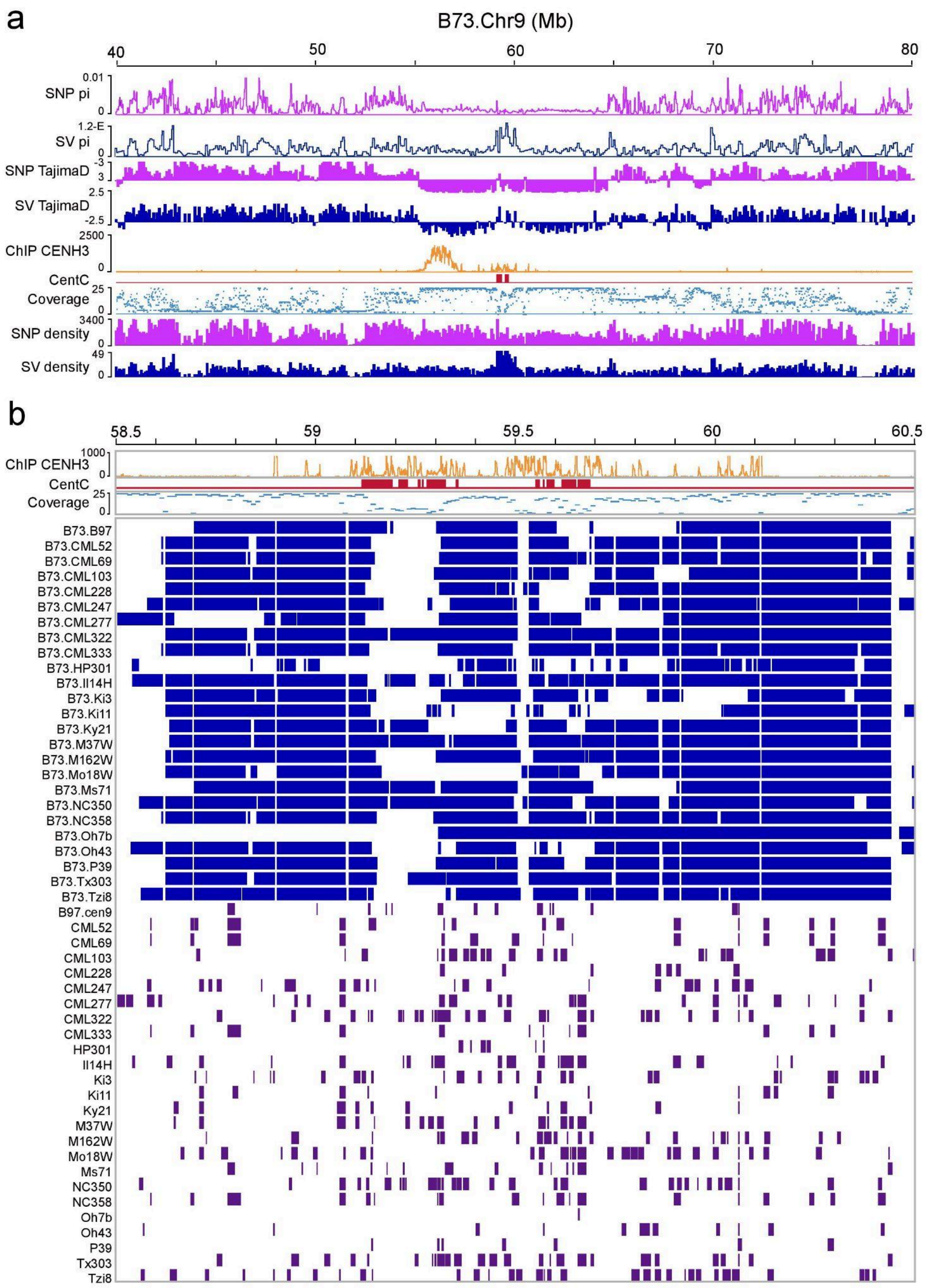


**Supplementary Figure 10: Features of genetic variations and their population properties around the functional centromeric regions of chromosome 9. a**. Panels from top to bottom include: (1) average pairwise diversity (π) for SNPs calculated in non-overlapping 100-bp windows along the chromosome; (2) π calculated for structural variants (SVs); (3) Tajima's D calculated with SNPs; (4) Tajima's D calculated with SVs; (5) CENH3 ChIP-seq peaks; (6) distribution of centromere-specific tandem repeats (CentC); (7) number of genomes aligned to the B73 genome in each 10-kb window; (8) SNP density; and (9) SV density. **b.** Functional centromeric regions are enriched with insertions of centromere-specific retrotransposable elements. Both sequence alignment coverage (blue) and the presence of transposable element (TE) insertions (purple) are shown for 26 maize genomes relative to the B73 reference genome.

### References


1. Bartenhagen, C. & Dugas, M. RSVSim: an R/Bioconductor package for the simulation of structural variations. *Bioinformatics* **29**, 1679–1681 (2013).
2. Kapitonov, V. V. & Jurka, J. Molecular paleontology of transposable elements in the Drosophila melanogaster genome. *Proc Natl Acad Sci USA* **100**, 6569–6574 (2003).
3. Hoskins, R. A. *et al.* The Release 6 reference sequence of the Drosophila melanogaster genome. *Genome Res* **25**, 445–458 (2015).
4. Chakraborty, M. *et al.* Hidden genetic variation shapes the structure of functional elements in Drosophila. *Nat. Genet.* **50**, 20–25 (2018).
5. Hoyt, S. J. *et al.* From telomere to telomere: The transcriptional and epigenetic state of human repeat elements. *Science* **376**, eabk3112 (2022).
6. Nurk, S. *et al.* The complete sequence of a human genome. *Science* **376**, 44–53 (2022).
7. Song, J.-M. *et al.* Two gap-free reference genomes and a global view of the centromere architecture in rice. *Mol. Plant* **14**, 1757–1767 (2021).
8. Hosmani, P. S. *et al.* An improved de novo assembly and annotation of the tomato reference genome using single-molecule sequencing, Hi-C proximity ligation and optical maps. Preprint at https://doi.org/10.1101/767764.
9. Alonge, M. *et al.* Major Impacts of Widespread Structural Variation on Gene Expression and Crop Improvement in Tomato. *Cell* **182**, 145–161.e23 (2020).
10. Hufford, M. B. *et al.* De novo assembly, annotation, and comparative analysis of 26 diverse maize genomes. *Science* **373**, 655–662 (2021).
11. Liao, Y. *et al.* The 3D architecture of the pepper genome and its relationship to function and evolution. *Nat. Commun.* **13**, 3479 (2022).
12. Qin, C. *et al.* Whole-genome sequencing of cultivated and wild peppers provides insights into Capsicum domestication and specialization. *Proc. Natl. Acad. Sci. U. S. A.* **111**, 5135–5140 (2014).
13. Kiełbasa, S. M., Wan, R., Sato, K., Horton, P. & Frith, M. C. Adaptive seeds tame genomic sequence comparison. *Genome Res.* **21**, 487–493 (2011).
14. Marçais, G. *et al.* MUMmer4: A fast and versatile genome alignment system. *PLoS Comput. Biol.* **14**, e1005944 (2018).

15. Li, H. Minimap2: pairwise alignment for nucleotide sequences. *Bioinformatics* **34**, 3094–3100 (2018).
16. Song, B. *et al.* AnchorWave: Sensitive alignment of genomes with high sequence diversity, extensive structural polymorphism, and whole-genome duplication. *Proc. Natl. Acad. Sci. U. S. A.* **119**, (2022).
17. Lin, H.-N. & Hsu, W.-L. GSAlign: an efficient sequence alignment tool for intra-species genomes. *BMC Genomics* **21**, 182 (2020).
18. Nattestad, M. & Schatz, M. C. Assemblytics: a web analytics tool for the detection of variants from an assembly. *Bioinformatics* **32**, 3021–3023 (2016).
19. Goel, M., Sun, H., Jiao, W.-B. & Schneeberger, K. SyRI: finding genomic rearrangements and local sequence differences from whole-genome assemblies. *Genome Biol.* **20**, 277 (2019).
20. Heller, D. & Vingron, M. SVIM-asm: structural variant detection from haploid and diploid genome assemblies. *Bioinformatics* **36**, 5519–5521 (2021).
21. O'Donnell, S. & Fischer, G. MUM&Co: accurate detection of all SV types through whole-genome alignment. *Bioinformatics* **36**, 3242–3243 (2020).
